# Human colorectal pre-cancer atlas identifies distinct molecular programs underlying two major subclasses of pre-malignant tumors

**DOI:** 10.1101/2021.01.11.426044

**Authors:** Bob Chen, Eliot T. McKinley, Alan J. Simmons, Marisol A. Ramirez-Solano, Xiangzhu Zhu, Austin N. Southard-Smith, Nicholas O. Markham, Quanhu Sheng, Julia L. Drewes, Yanwen Xu, Cody N. Heiser, Yuan Zhou, Frank Revetta, Lynne Berry, Wei Zheng, M. Kay Washington, Qiuyin Cai, Cynthia L. Sears, James R. Goldenring, Jeffrey L. Franklin, Simon Vandekar, Joseph T. Roland, Timothy Su, Won Jae Huh, Qi Liu, Robert J. Coffey, Martha J. Shrubsole, Ken Lau

## Abstract

Most colorectal cancers (CRCs) develop from either adenomas (ADs) or sessile serrated lesions (SSLs). The origins and molecular landscapes of these histologically distinct pre-cancerous polyps remain incompletely understood. Here, we present an atlas at single-cell resolution of sporadic conventional tubular/tubulovillous ADs, SSLs, hyperplastic polyps (HPs), microsatellite stable (MSS) and unstable (MSI-H) CRC, and normal colonic mucosa. Using single-cell transcriptomics and multiplex imaging, we studied 69 datasets from 33 participants. We also examined separate sets of 66 and 274 polyps for RNA and targeted gene sequencing, respectively. We performed multiplex imaging on a tissue microarray of 14 ADs and 15 CRCs, and we integrated pre-cancer polyp data with published single-cell and The Cancer Genome Atlas (TCGA) bulk CRC data to establish potential polyp-cancer relationships. Striking differences were observed between ADs and SSLs that extended to MSS and MSI-H CRCs, respectively, reflecting their distinct origins and trajectories. ADs arose from WNT pathway dysregulation in stem cells, which aberrantly expanded and expressed a Hippo and ASCL2 regenerative program. In marked contrast, SSLs were depleted of stem cell-like populations and instead exhibited a program of gastric metaplasia in the setting of elevated cytotoxic inflammation. Using subtype-specific gene regulatory networks and shared genetic variant analysis, we implicated serrated polyps, including some HPs conventionally considered benign, as arising from a metaplastic program in committed absorptive cells. ADs and SSLs displayed distinct patterns of immune cell infiltration that may influence their natural history. Our multi-omic atlas provides novel insights into the malignant potential of colorectal polyps and serves as a framework for precision surveillance and prevention of sporadic CRC.

## Introduction

Colorectal cancer (CRC) is the fourth most common type of cancer and second leading cause of cancer death in the United States (Siegel et al., 2019). Most CRCs develop from pre-cancerous polyps, which are common in adults and targeted for removal during colonoscopy (Baron et al., 2013). However, despite advances in endoscopic techniques, polypectomy may not prevent carcinoma in as many as 24% of cases nor is it fully effective in preventing death from CRC located in the proximal colon (Baron et al., 2013; Baxter et al., 2012; Brenner et al., 2010). For precision targeting, several CRC classification schemes have been adopted based on biological features such as microsatellite-instability (MSI), chromosomal instability (CIN), methylation status (CpG Island Methylator Phenotype or CIMP), and transcriptomic profile (Consensus Molecular Subtypes or CMS) (Dienstmann et al., 2017; Guinney et al., 2015; Ogino and Goel, 2008). Some of these molecular characteristics are evident in polyps, and yet, a comprehensive depiction of pre-cancerous polyps beyond their similarity to CRC has not been described (Komor et al., 2018). Understanding the origins and identifying the unique molecular phenotypes of precursor lesions may facilitate new strategies for precision prevention, surveillance, and therapeutic approaches for CRC.

The conventional adenoma (AD) is the precursor lesion for approximately 65-85% of sporadic CRC (Dickinson et al., 2015; Fearon, 2011). As proposed by Vogelstein and co-workers in a classic model, ADs arise from truncating mutations in *APC*, which result in activation of the WNT pathway and CIN (Fearon and Vogelstein, 1990). As ADs progress, they accumulate additional genetic events, for example, gain-of-function mutations in oncogenes (chiefly *KRAS*) and loss-of-function mutations in tumor suppressor genes such as *p53*. While these key driver mutations and signaling pathways in CIN-type CRC have largely been identified, the single-cell molecular landscape of ADs within their microenvironment remains incompletely resolved.

Although the molecular circuitry associated with AD progression is increasingly described, much less is known about sessile serrated lesions (SSLs), formerly known as sessile serrated adenomas/polyps. SSLs are precursors for as much as 35% of CRCs, although a consensus diagnosis for SSLs was not available until 2010 (Albores S, J.; Adsay, N. V.; Crawford, 2010; Crockett and Nagtegaal, 2019; Longacre and Fenoglio-Preiser, 1990; Rex et al., 2012). They are less common than ADs, representing only 10-20% of polyps, and are more often in the proximal colon, unlike ADs (Crockett and Nagtegaal, 2019; Jones et al., 2008; Markowitz and Bertagnolli, 2009). SSLs are distinct from ADs in that tumorigenesis is not initiated by genetic disruptions of the WNT pathway. Lesions frequently do not have *APC* mutations, and they have epigenetic disruptions including *MLH1* hypermethylation and a 40-75% prevalence of CIMP (Leggett and Whitehall, 2010; Yang et al., 2004). SSLs resemble MSI-H CRCs, which often share these features alongside WNT pathway dysregulation not originating from *APC* mutations (Crockett and Nagtegaal, 2019; Thorstensen et al., 2005).

Substantial effort has been expended on understanding cell-intrinsic tumor-initiating events, but the confluence of extrinsic factors, such as diet, NSAID use, and microbiota alterations, are also critical for neoplastic transformation in the colon (Hullar et al., 2014). Differentiated epithelial cells are vulnerable to microenvironmental injury and can activate repair mechanisms reliant on cellular plasticity, involving a change of cell state into a regenerative state favorable to tumorigenesis (Van Es et al., 2012; Jones et al., 2019a; Tetteh et al., 2016; Vega et al., 2019; Yan et al., 2017b; Yu et al., 2018). Conversely, normal stem cells residing in the crypt base are largely protected from luminal stressors (Kaiko et al., 2016), but they are constantly exposed to replicative stress due to continual proliferation (Barker, 2014). Differing microenvironmental influences are posited to promote malignant transformation in the CRC cell-of-origin, and, in relation to its developmental state, may also require different pathways for tumorigenesis.

This multi-omic Human Tumor Atlas Network study describes the integration of single-cell transcriptomics, genomics, and immunohistopathology to identify the origins and molecular underpinnings of the two major types of pre-cancerous polyps in the colon and the distinct routes they take to CRC. We highlight critical regulatory processes and microenvironmental changes observed in early tumorigenesis that may elucidate new phenotypes and inform cancer prevention strategies.

## Results

### Distinct molecular features of colonic pre-cancer subtypes compared with normal colon

COLON MAP study participants were recruited among individuals undergoing routine screening or surveillance colonoscopy with no known or suspected personal history of colon resection, inflammatory bowel disease, polyposis syndromes, Lynch syndrome, or cancers other than nonmelanoma skin cancers. Polypoid lesions, as well as normal colonic biopsies, were obtained during colonoscopy. We generated concurrent single-cell RNA-sequencing (scRNA-seq), multiplex immunofluorescence (MxIF), and multiplex immunohistochemical (MxIHC) data from 33 individuals for a total of 69 specimens analyzed **(Figure 1A)**. We performed orthogonal bulk RNA-seq and targeted gene sequencing on an additional set of 66 and 274 pre-cancer specimens, respectively **(Table S1)**.

**Figure 1:**
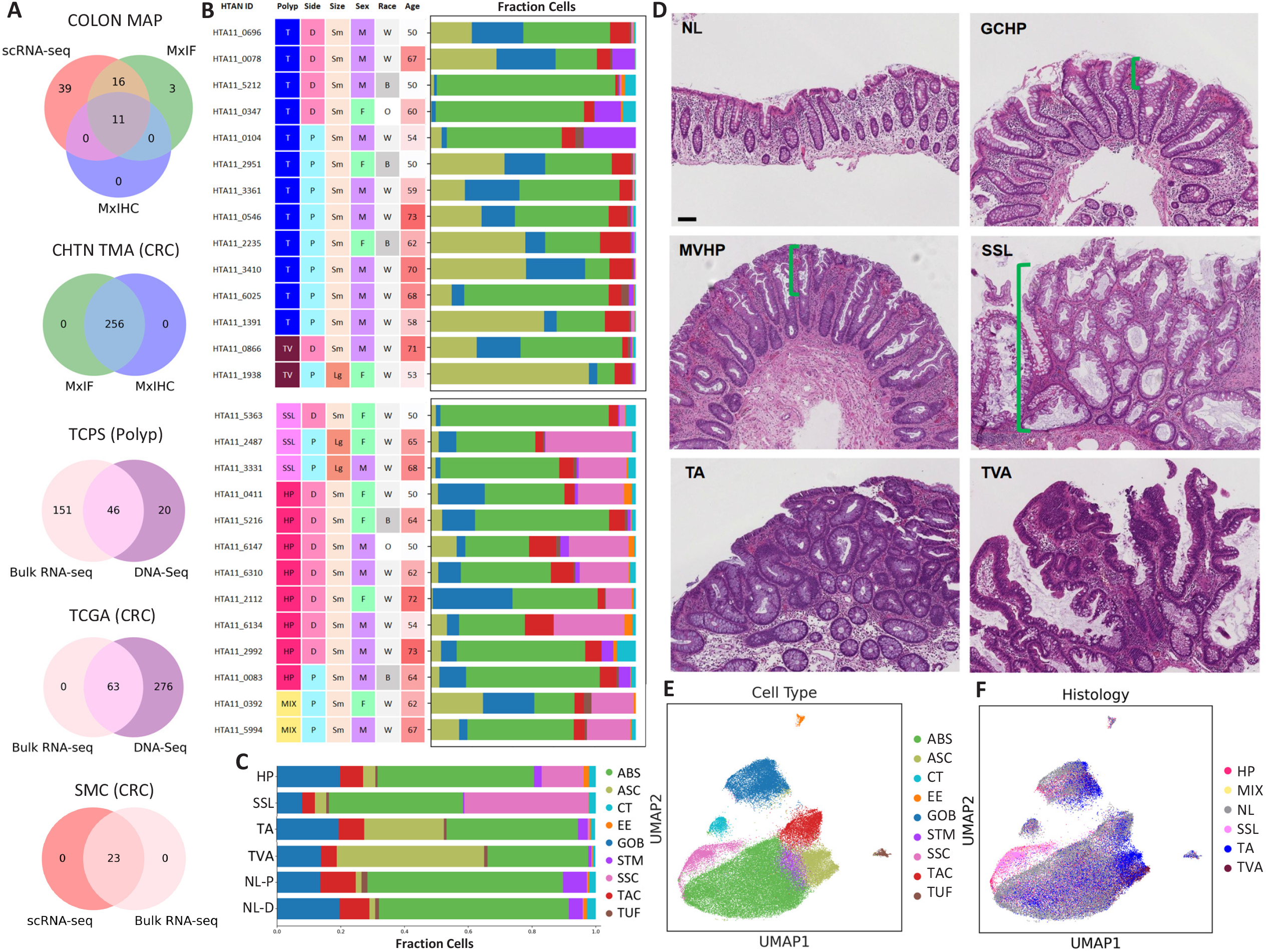
Cellular and histological features of colonic pre-cancer subtypes. **(A)** Venn diagrams describing the multi-omic assays performed on the collections of samples from each analyzed participant set. **(B)** Polyp specimens for single-cell analysis categorized into serrated and conventional pathways, and annotated with patient metadata and histological subtyping by pathologists. Stacked bar plots describe epithelial cellular distribution of each specimen with colors representing cell types as defined by Leiden clustering and marker gene detection. ABS-absorptive cells, ASC-adenoma-specific cells, CT-crypt top colonocytes, EE-enteroendocrine cells, GOB-goblet cells, STM-stem cells, SSC-serrated-specific cells, TAC-transit amplifying cells, TUF-tuft cells. **(C)** Stacked bar plots with the same color scheme depicting combined cellular distribution per histological subtype for hyperplastic polyps (HP), sessile serrated lesions (SSL), tubulovillous adenomas (TVA), tubular adenomas (TA), proximal normal (NL-P), and distal normal (NL-D) colon tissue. **(D)** Representative histology of each polyp subtype: normal (NL) tissue, goblet cell-rich hyperplastic polyp (GCHP), microvesicular hyperplastic polyp (MVHP), sessile serrated lesion (SSL), tubular adenoma (TA), and tubulovillous adenoma (TVA). Green brackets label portions of crypts occupied by serrations in GCHP, MVHP, and SSL. Scale bar: 100 μm. **(E,F)** Combined regulon-based UMAP representation of scRNA-seq data of epithelial cells from normal and neoplastic tissues colored by **(E)** cell type defined by Leiden clustering and **(F)** histological subtype.

For scRNA-seq, we collected a total of 75,996 cells after removing barcodes that failed quality control thresholds using the automatic cell identification pipeline, dropkick (Heiser et al., 2020). Most samples were small specimens (with a median diameter of 5 mm or less). For pre-cancer-specific analysis, we selected a subset of 62 scRNA-seq datasets across normal biopsies and four pre-cancer lesion types defined by histological classification, with 12 tubular ADs, 2 tubulovillous ADs, 3 SSLs, 8 HPs, and 2 specimens with unconfirmed histology but transcriptomically classified as SSL (labeled MIX), as determined by consensus from 2 independent pathologists **(Figure 1B,C; Figure S1A,B)**. Histologically, the crypt architecture of ADs was largely maintained, but there was a loss of differentiation with fewer mature goblet and absorptive cells **(Figure 1D; Figure S1C)**. The nuclei were dark, enlarged, spindle-shaped, and pseudostratified with loss of the stereotypic basal location. ADs were subtyped by the proportion of villous component. Tubular ADs had less than 25% villous component and tubulovillous ADs had between 25-75%. In contrast, SSLs showed epithelial serrations extending to the crypt base and distortion of the crypt architecture with horizontal growth along the muscularis mucosae, crypt base dilation, and asymmetric proliferation. SSLs exhibited normal cytology without atypia, so-called bland cytology. These specimens came from individuals with the following characteristics: gender: 63% male, 37% female; race: 74% white, 19% black, 7% other; age: 41% 50-59 years, 41% 60-69 years, 18% ≥70 years; polyp location: 52% proximal, 48% distal **(Figure 1B)**. Normal colonic biopsies came from participants with the following characteristics: gender: 60% male, 40% female; race: 66% white, 28% black, 6% other; age: 51% 50-59 years, 37% 60-69 years, 12% ≥70 years, tissue location: 49% proximal, 51% distal **(Figure S1A,B)**. Other characteristics of the participants are recorded in **Table S2**. Using raw scRNA-seq data, we observed intermixing of epithelial cells from normal colonic biopsies and immune cells from different participants, indicating the absence of batch effects over multiple runs **(Figure S2A,B)**. However, neoplastic tissues from different participants clustered by sample, demonstrating variability and individualized etiologies in contrast to normal biopsies.

To adjust for polyp-specific effects and reveal generalized principles across polyp subtypes, we applied Single-Cell rEgulatory Network Inference and Clustering (SCENIC), which utilizes regulon information for the batch-robust feature-extraction of scRNA-seq data (Aibar et al., 2017; Van de Sande et al., 2020). Transcription factor (TF)-defined regulon activities are considered to be more determinant of cell identity as opposed to environmentally sensitive genes such as metabolic genes (Aibar et al., 2017). Applying SCENIC to epithelial cell populations, we identified 154 enriched regulons that were used for data embedding, graph generation, and community detection by unsupervised Leiden clustering **(Figure S2C,D)** (Traag et al., 2019). From the resultant regulon nearest-neighbor graph generated by co-embedding 62 scRNA-seq datasets, we identified 9 major populations of epithelial cells **(Figure 1E)**. Normal colonic biopsy tissues served as reference landmarks with cells falling into well-known canonical epithelial cell types, including secretory goblet cells marked by *MUC2* and *ATOH1*, absorptive cells marked by *KRT20* and *GUCA2A*, *BEST4+/MEIS1+* crypt top colonocytes, enteroendocrine cells marked by CHGA and *NEUROD1*, tuft cells marked by *POU2F3* and *SOX9*, transit-amplifying cells marked by *PCNA* and *MKI67*, and stem cells marked by *LGR5* and *OLFM4* **(Figure 1C,E; Figure S3)**. Within this combined cell space, we identified two populations overrepresented by neoplastic lesion-specific cells **(Figure 1C,F; Figure S3)**. One population was enriched with tubular and tubulovillous AD cells (AD-specific cells - ASCs), representing the conventional tumorigenesis subclass. The other population was enriched with cells from SSLs (serrated-specific cells - SSCs), representing the serrated pathway subclass. It should be noted each lesion also contained transcriptionally normal epithelial cells, consistent with the histopathology showing only a portion of each resected polyp was neoplastic in appearance **(Figure S4)**. These analyses showed our data to be high-quality and consistent with known colonic biology, enabling confident identification of neoplastic cells for further analysis.

The malignant potential of hyperplastic polyps (HPs) is controversial. HPs were subdivided into goblet cell-rich (GCHP) and microvesicular (MVHP) based on histopathology **(Figure 1D; Figure S1C)** (Leggett and Whitehall, 2010). In GCHPs, most of the cells were goblet cells and the crypts were taller and wider than in normal mucosa. Epithelial serrations, if present, were confined to the surface. In MVHPs, the epithelium was composed of goblet cells and epithelial cells, some of which contained microvesicular mucin. Epithelial serrations were present in the upper two thirds of the crypts. Importantly, for both types of HPs, the crypt base was morphologically normal with a confined proliferative zone and basally located nuclei that were small and round to oval. We observed concordance between this histology and our scRNA-seq data, as a subset of HP cells contained regulatory signatures similar to SSLs and clustered with the SSC population **(Figure 1B,C,F)**. Specifically, all MVHP lesions contained substantial SSCs, and the HP subset contained SSC cells proportional to the degree of epithelial serrations **(Figure 1F)**. The majority of GCHPs were comprised of transcriptionally normal cells, suggesting a low risk for malignant progression. These results are consistent with MVHPs arising from similar genetic and epigenetic alterations as the serrated pathway, such as activating mutations in *BRAF*/*KRAS* and methylation of *MLH1* (Jass, 2004).

Due to the small polyp size and our prioritization of assays, there was insufficient material for paired genetic analysis. Thus, for mutational analysis, we evaluated a dataset from an independent study of 274 polyps, where we performed targeted sequencing of genes commonly reported in CRC **(Figure 2)**. This included 217 tubular ADs, 50 tubulovillous ADs, and 7 SSLs. As expected, the majority of ADs had *APC* mutations, occurring in 67% (146/217) of tubular ADs and 90% (45/50) of tubulovillous ADs, whereas only a single *APC* mutation was detected in SSLs. Likewise, mutations in *KRAS* increased markedly as ADs progressed histologically, with 6% (13/217) in tubular and 40% (20/50) in tubulovillous ADs; no *KRAS* mutations were detected in SSLs. Few *BRAF* mutations were observed in this study population (3% - 7/274). However, they were markedly enriched in SSLs with 57% (4/7 - 3 V600E and 1 G6S mutation) compared to 0.0046% (1/217) in tubular and 4% (2/50) in tubulovillous ADs. WNT pathway mutations were found in 89% of cases overall owing to the high number of *APC* mutations. Of note, WNT pathway mutations were not equally distributed: 88% in tubular ADs, 96% in more histologically advanced tubulovillous ADs, and only 43% in SSLs, consistent with the paradigm of serrated lesions initiated by a distinct driver pathway and incurring WNT pathway mutations later. Activation of the WNT pathway in SSLs occurs independently of traditional *APC* mutations (Hashimoto et al., 2017; Yan et al., 2017a). Although no *RNF43* or *ZNRF43* mutations were detected in SSLs, mutations did occur in *FAT1* and *NKD1*. Of note, *APC2* mutations were also present in ADs, which in some cases, overlapped with APC mutations, possibly reflecting polyclonality within these pre-malignant polyps. The Hippo pathway was activated in 22% and 14% of tubular and tubulovillous ADs, respectively, but in none of the SSLs, while components of Notch pathway were mutated in 24% and 14% of tubular ADs and tubulovillous ADs, respectively. Notch pathway mutations appeared to be overrepresented in SSLs (57%), with mutations in *JAG2* and *NOTCH3*. TGF-β pathway mutations were found in less than 5% of ADs, and none in SSLs, consistent with these mutations occurring late in the neoplastic process (Grady et al., 1998). While MSI-H cancers are characterized by hypermutation, large numbers of mutations were surprisingly not observed in SSLs; in contrast, one AD had a hypermutator phenotype **(Figure 2)**. Our mutational profiling of pre-cancer polyps paints a picture of WNT-driven tumorigenesis in ADs, accompanied by Hippo pathway alterations, while SSLs are less dependent on this pathway and are not hypermutated.

**Figure 2:**
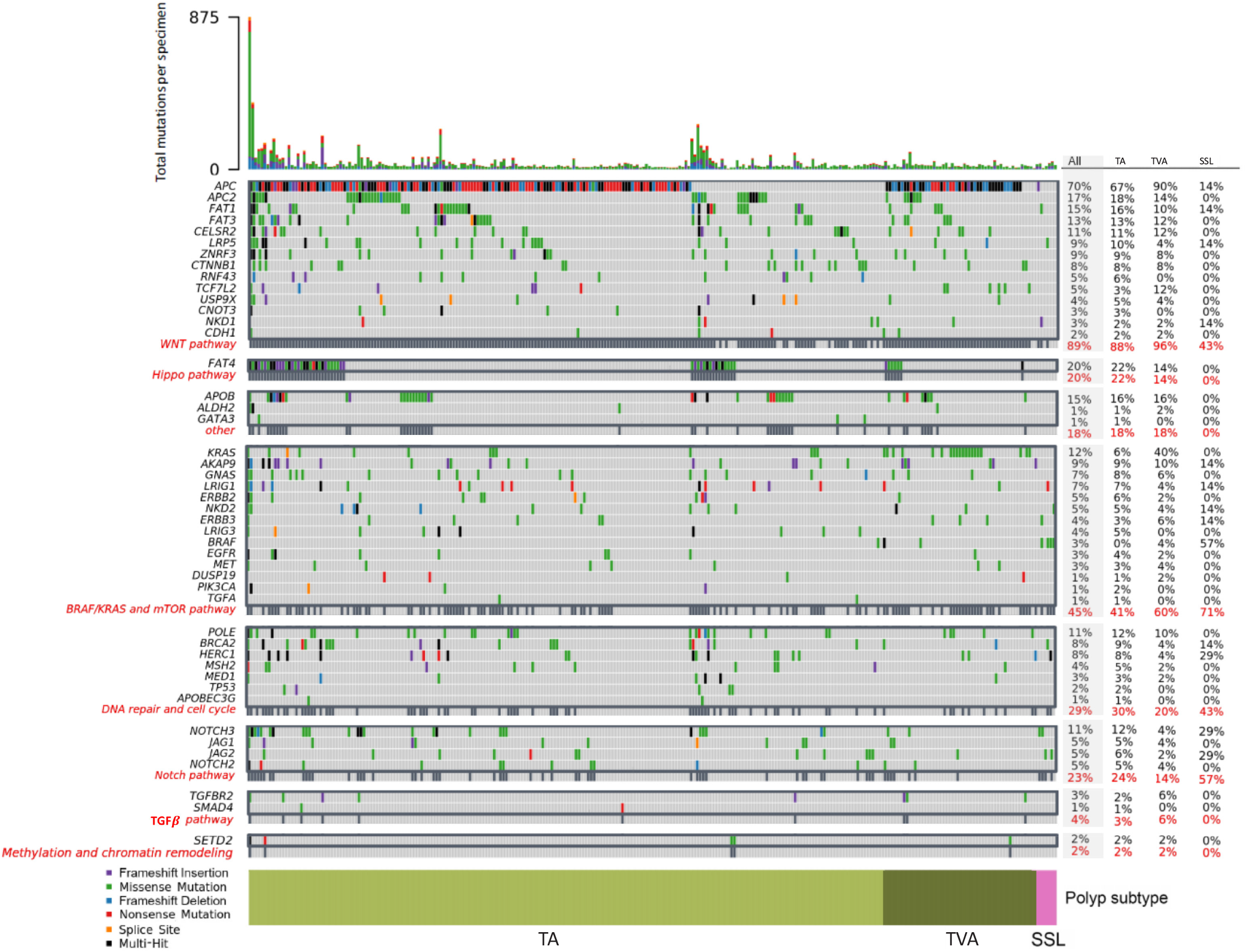
Mutational analysis of colonic pre-cancers. Heatmap representation of the mutational landscape of 274 specimens by targeted sequencing with different types of mutations are color-coded. Total number of mutations detected per sample represented as bar plot (top). Important genes to CRC are presented, grouped into 8 pathways. Proportion of gene and pathway mutation within polyp subtypes summarized as table (right).

### Neoplastic cells in conventional adenomas and serrated polyps present different gene regulatory networks

Next, we focused on identifying gene programs and pathways activated in these two classes of premalignant epithelial cells. Single-cell transcriptomic analysis uncovered distinct clusters for ASCs and SSCs, comprised of both SSLs and HPs with serrated morphologies. Both gene- and regulon-based analysis demonstrated that ASCs resemble stem and transit-amplifying progenitor cells. Molecularly, ASCs from both tubular and tubulovillous ADs highly express genes indicative of WNT pathway activation (*LGR5*, *OLFM4*, *ASCL2*, *AXIN2*, *RNF43*, and *EPHB2*; **Figure 3A**) at a level higher than crypt base columnar stem cells in matching normal colon biopsies from the same individuals. Correspondingly, WNT-related TF activities (TCF7, ASCL2, FOXQ1, and SMARCA4) (Cadigan and Waterman, 2012; Christensen et al., 2013; Park et al., 2009; Schuijers et al., 2015), as defined by sets of predicted target genes organized as regulons, were also enriched **(Figure 3B)**. Additionally, genes known to be expressed at the crypt base (e.g., *CD44* and *CLDN2*) were enriched in this cluster, demonstrating the stem-like nature of this cluster **(Figure 3A)**. TGF-β signaling is also upregulated compared with SSCs, as revealed by *TGFBI*, *TGFBR2*, *SMAD2*, *SMAD3*, and *EP300* expression **(Figure S5A)**. We then applied CytoTRACE to infer the stem potential of single cells through a smoothed regression model of transcriptional diversity (Gulati et al., 2020). Colonic epithelial cells from normal tissue biopsies exhibited stem cells with high CytoTRACE scores transitioning into differentiated cells with lower CytoTRACE scores **(Figure S5B,C)**. Due to a distribution consisting of a mix of stem, transitioning, and differentiated cells in the normal colonic epithelium, we observed a relatively uniform CytoTRACE score distribution, as calculated through kernel density estimation, which differed significantly from the ASC and SSC distributions **(Figure 3C; Figure S5C; Table S3)**. For ASCs, CytoTRACE analysis yielded a distribution skewed towards cells with high predicted stem potential **(Figure 3C)**. Genes most highly correlated with the CytoTRACE score were enriched with a heavy ribosomal signature, which comprised the MYC regulon similarly enriched in ASCs and downstream of canonical WNT signaling **(Figure S5D)**. Among ASCs, we observed a stem-like subpopulation highly expressing *LGR5*, *ASCL2*, and *NKD1* **(Figure S6A-C)**. These cells were associated with GO terms related to cell junctions and exosomal processes, suggesting these cells may maintain the tumor microenvironment through extracellular communication **(Figure S6D-F; Table S4-S6)**. These results support a WNT-dependent stem cell expansion model of tumorigenesis in conventional adenomas, which is consistent with known WNT pathway activating gene mutations prevalent in CRC initiation, most notably loss-of-function mutations in *APC* (Korinek et al., 1997).

**Figure 3:**
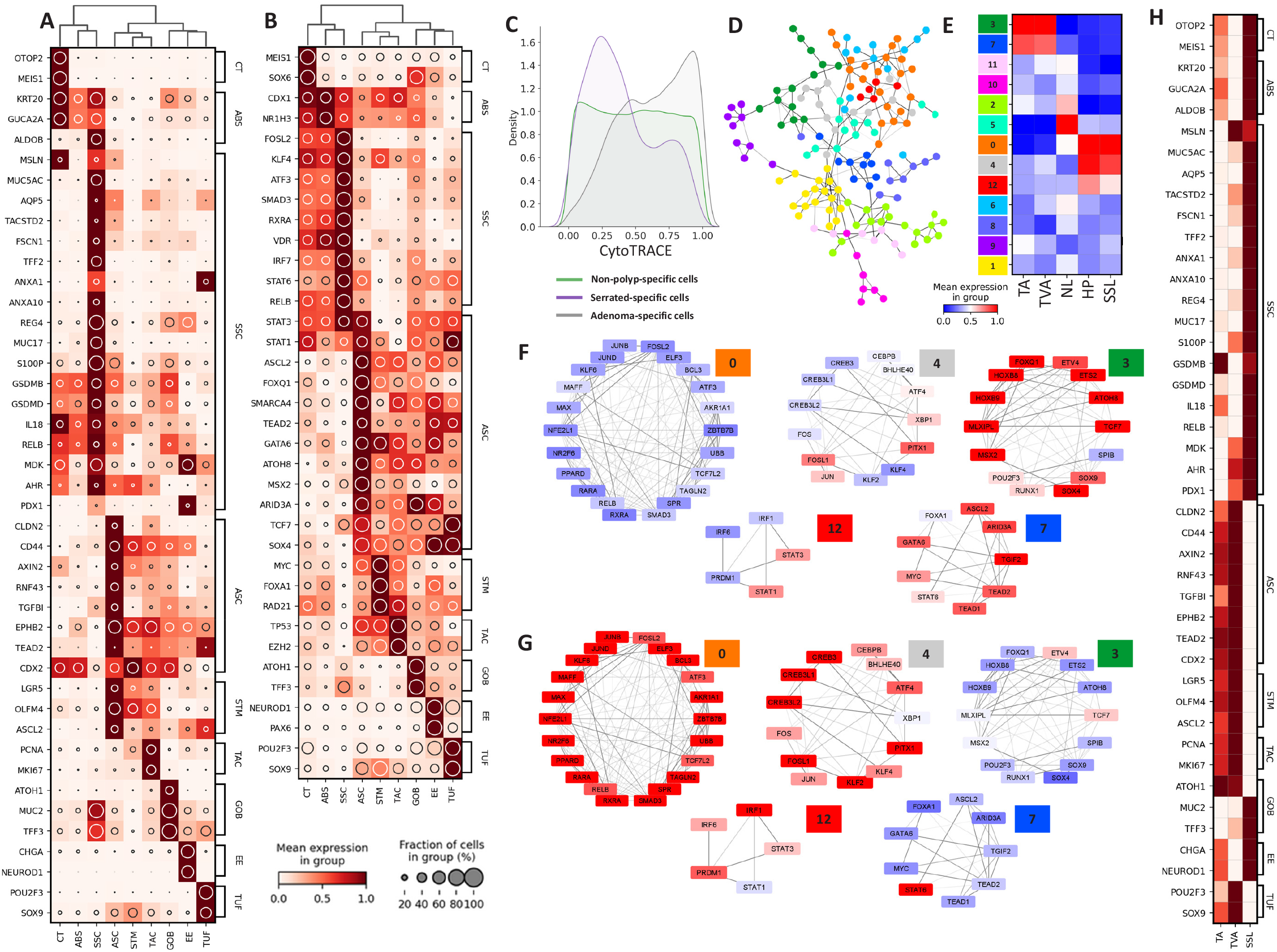
Transcriptomic landscape divergence between conventional and serrated polyps. **(A,B)** Heatmap representation of top biologically relevant and differentially expressed **(A)** genes and **(B)** regulon activities (enrichment scores) from each cell cluster. The mean expression is depicted by heat intensity and prevalence at single-cell level is depicted by the size of the inset circle. **(C)** Density plot of single-cell CytoTRACE scores calculated for cells specific for each lesion type compared with normal cells. **(D)** TF target network created from normal and pre-cancer cells, broken down into subnetworks by Leiden subgraph clustering (color). Nodes represent individual TFs and edge opacity denotes similarities in gene targets. **(E)** Heatmap depicting average regulon enrichment scores per sample histology of each subnetwork. **(F,G)** TF target subnetworks selected for differentially enrichment between polyps from the conventional **(F)** and serrated **(G)** pathways. Color overlays for each TF node are the average regulon enrichment scores per subclassification and edge opacities are the inferred TF-target weightings. **(H)** Heatmap representation of gene signatures identified by scRNA-seq of pre-cancers applied to bulk RNA-seq samples of various pre-cancer subtypes (n=36 tubular, 22 tubulovillous, 8 SSLs). The mean expression is depicted through color intensity.

In marked contrast to ASCs, SSCs did not show WNT pathway activation. Moreover, CytoTRACE scores of SSCs were inversely skewed towards cells with low predicted stem potential, and SSCs shared transcriptomic similarity with differentiated cells **(Figure 3A,C; Figure S5C)**. The overall transcriptomic and regulon profiles of SSCs resembled absorptive-lineage cells, but SSCs also expressed functional goblet cell genes, including *TFF3* and *MUC2*. Unlike normal goblet cells, they did not express the master secretory cell TF *ATOH1* nor were they enriched for ATOH1 TF activity, suggesting SSCs harbor a mixed cellular identity **(Figure 3A,B)**. Of note, SSCs were highly enriched for genes not normally expressed in the colon (*MUC5AC*, *AQP5*, *TACSTD2* (TROP2), *TFF2*, *MUC17, and MSLN*), but instead expressed in other endodermal organs, most notably the stomach **(Figure 3A)**. This led us to hypothesize gastric metaplasia may underlie the etiology of SSLs.

Metaplasia is a process in which differentiated cells transdifferentiate to express non-native cell types. It often occurs in the setting of epithelial damage, which activates a regenerative process inducing normally quiescent differentiated cells to re-enter the cell cycle. Loss of *CDX2* in the colon is associated with an imperfect pyloric-type gastric metaplasia (Balbinot et al., 2018; Tong et al., 2017). *CDX2* is a homeobox gene critical for hindgut specification and with its loss there is a shift towards expression of genes more rostral in the rostral-caudal gradient (Stringer et al., 2012). *CDX2* was expressed in most colonic cell types, including ASCs; however, it was downregulated in SSCs, supporting a loss of regional identity in these cells **(Figure 3A; Figure S3)**. In addition to this metaplastic process, we also observed reversion to an embryonic state in SSCs, as exemplified by re-expression of *MDK* (**Figure 3A; Figure S3**), a heparin-binding growth factor transiently expressed in early colonic development during mouse embryogenesis (Park et al., 2005) with limited expression in adult organs (Kadomatsu et al., 2013; Park et al., 2005) **(Figure 3A)**. Supporting the role of a damage-induced process, we also observed an inflammatory signature in SSCs, characteristic of an exposure of surface epithelial cells to microbes. Upregulated genes, such as *S100P* and NF-κB (*RELB*), and regulons, such as STAT6, STAT3, and IRF7, reflect the immunogenic state of these cells **(Figure 3A,B)**. GSEA further corroborates this finding by enriching for gene ontology (GO) terms and pathways related to the detection of microbial infection, activation of the innate immune system, and epithelial wound healing **(Figure S7; Table S7-10)** (Raudvere et al., 2019). Specifically, coordinated upregulation of inflammasome-related genes such as *IL18* and gasdermins was observed **(Figure 3A)**. SSCs also contain a subpopulation with higher CytoTRACE scores indicative of stem-like cells associated with the GO terms ribosomal processes and innate immune responses **(Figure S8; Table S11-14)**. Similarly, regulons related to FOSL2, KLF4, and ATF3 were enriched **(Figure 3B)**, drawing parallels to work by Ansari *et al*., who observed increased chromatin accessibility and hypomethylation of these TF targets that occurred in a mouse model of colitis in a microbiota-dependent manner (Ansari et al., 2020). These findings are consistent with the observed higher prevalence of biofilms with accompanying invasive bacteria in proximal CRCs, which are also predominantly found in the proximal colon (Drewes et al., 2017).

To assess global gene regulatory differences between ASCs and SSCs, we used predicted TF target similarity as a metric to create a common TF network. We retrieved 13 subnetworks of highly connected TFs using the Leiden community detection algorithm **(Figure 3D)**, and the average enrichment of these regulon communities was calculated, resulting in detection of 5 subnetworks uniquely enriched between ASCs (subnetworks 3 and 7) and SSCs (subnetworks 0, 4, 12) **(Figure 3E)**. These subnetworks represented coordinated regulation amongst multiple TFs and enabled the identification of gene programs and pathways **(Figure S9)**. Following gene set and pathway analysis **(Figure S7)**, we identified two modules from the two subnetworks enriched in ASCs: (1) a WNT-driven stem cell module and (2) a Hippo signaling driven regeneration module **(Figure 3F; Figure S9C-E)**. These results were consistent with the role of Hippo signaling and an ASCL2 transcriptional complex in the damage-regeneration response of intestinal stem cells (Ayyaz et al., 2019; Murata et al., 2020). For SSCs, significantly represented subnetworks form a super module pointing towards interleukin signaling and microbiota interaction through receptors for retinoic acid (RA) (RXRA/RARA), vitamin D (VDR), and aryl hydrocarbon (AHR) **(Figure 3A,G; Figure S9F-H)**. Importantly, a recent paper reported exogenous RA signaling suppressed the YAP-dependent regenerative stem state and promoted absorptive cell differentiation, consistent with the “absorptive” identity of SSCs (Lukonin et al., 2020). We furthered gleaned signature genes from organoids under RA signaling activation and found similar genes, such as *ALDOB*, to be upregulated in SSCs **(Figure 3A)**. There were notable differences between the two datasets, however, including genes marking regional identities dysregulated in SSCs compared to RA - induced absorptive cells in organoids. *ANXA10*, a gastric identity gene, and *ANXA1*, a fetal and esophageal identity gene, were upregulated in SSCs but were suppressed in RA-induced absorptive cell differentiation **(Figure 3A)** (Kim et al., 2015; Paweletz et al., 2000; Tsai et al., 2015; Yui et al., 2018). Furthermore, a subnetwork depicting enhanced metabolic activity was observed in SSCs, consistent with metabolic changes conferred by the RAS/RAF pathway activation **(Figure S9F-H)**. As validation, we performed bulk RNA-seq on an additional 58 ADs (36 tubular, 22 tubulovillous) and 8 SSLs. A signature consistent with gastric metaplasia described by our single-cell data (*MUC5AC*, *ANXA1*, *ANXA10*, *AQP5*, *TACSTD2* (TROP2), *TFF2*, and *MUC17*) was enriched in the SSLs from the validation set (**Figure 3H; Figure S10**). Compared to ADs in this set, we observed a marked depletion in *ATOH1* expression paired with the paradoxical enrichment of goblet cell genes such as *TFF3* and *MUC2*. We observed a consistent WNT-driven stem cell signature (*LGR5*, *OLFM4*, *ASCL2*, *AXIN2*, *RNF43*, and *EPHB2*) in ADs, with tubulovillous ADs exhibiting slightly higher enrichment scores **(Figure 3H; Figure S10)**. These results implicate a WNT-activated regenerative program of stem cells in conventional ADs, and a gastric metaplastic program, likely arising from a committed absorptive lineage in SSLs.

### Lineage analysis implicates the origins of conventional adenomas and serrated lesions

Having shown differences in regulatory programs between ASCs and SSCs, the former driven by WNT-activated stem cell expansion and the latter driven by gastric metaplasia coupled to inflammation, we sought evidence for the cellular origins of these tumor subtypes. Metaplasia arises when damage to differentiated cells induces a transition to a regenerative cell state for repair. Thus, metaplastic lesions may develop from differentiated cells in a “top-down” model of tumorigenesis. In contrast, ADs accrue mutations in proliferative stem cells residing at the crypt base. Loss-of-function mutations in *APC* are often the initiating events, resulting in activation of the WNT pathway. Constitutive activation of WNT prevents differentiation and maintains an aberrant stem-like state.

To determine the origins of neoplastic cells, we first inferred cell-state transition trajectories using p-Creode on the regulatory network landscape, derived through SCENIC, of colonic epithelial cells (Herring et al., 2018). This analysis produced a batch-robust developmental hierarchy where the seven major cell types are organized into lineage branches **(Figure 4A)**. As expected, we observed a secretory lineage branching into goblet, tuft, and enteroendocrine cells, and an absorptive lineage consisting of proliferating transit-amplifying cells, colonocytes, and *BEST4+/MEIS1+* crypt top colonocytes **(Figure S11)**. A CytoTRACE score overlay aligns with the hierarchy, as highly scored cells were enriched in progenitor and stem cells, the latter being enriched for *AXIN2* expression, reflecting active WNT signaling **(Figure 4B; Figure S11)**. Importantly, ASCs shared a branch with stem cells, suggesting the developmental origins of conventional ADs may lie in the aberrant expansion of crypt base stem cells, or “bottom-up” tumorigenesis. In marked contrast, SSCs, which expressed goblet cell genes such as *MUC2*, were paradoxically inferred to develop from absorptive progenitors and colonocytes **(Figure 4C; Figure S11)**.

**Figure 4:**
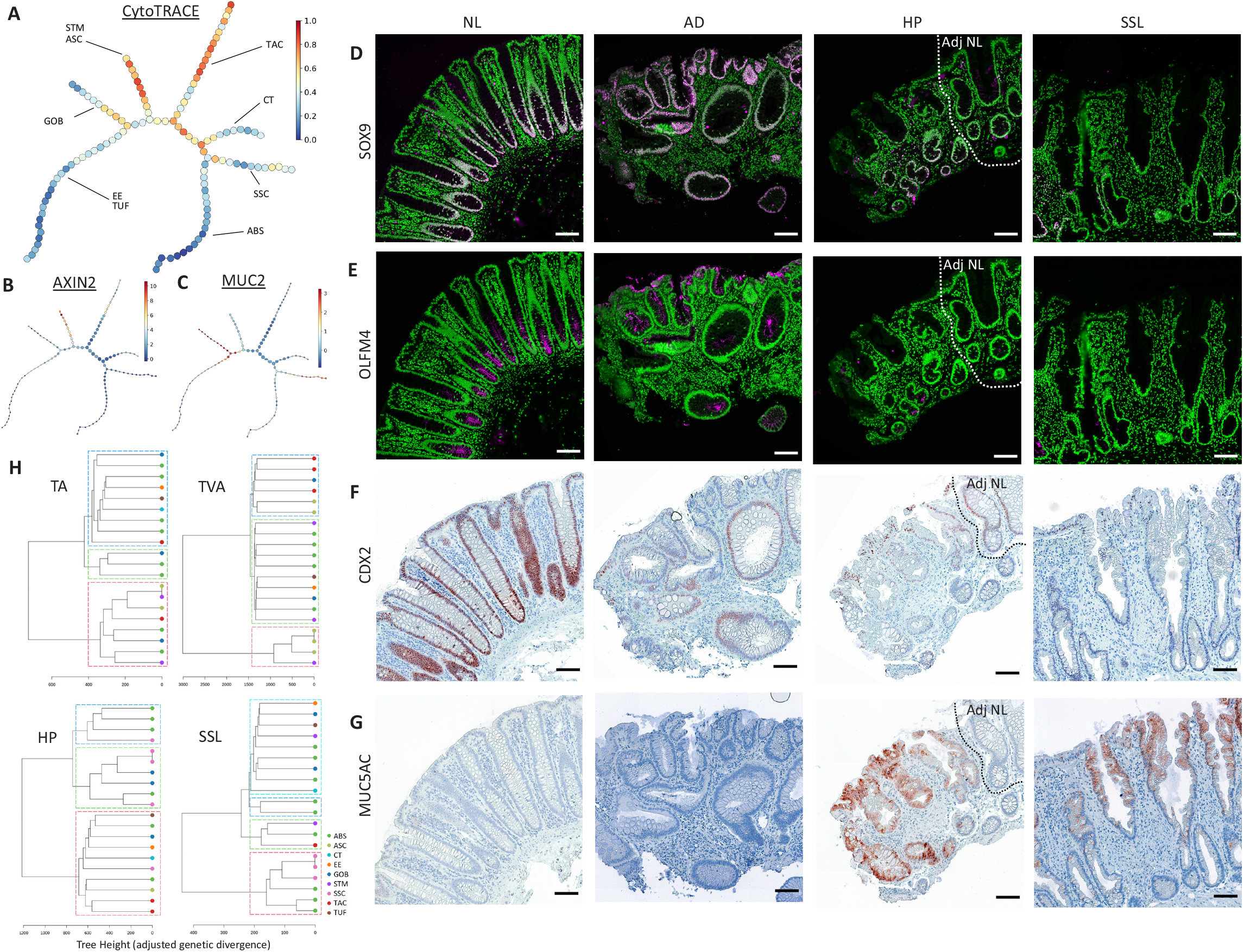
Inferred developmental origins of between conventional and serrated precancers. **(A)** Trajectory inference through p-Creode performed on the regulon landscape of cumulative complement of cells from the study. Overlay represents CytoTRACE predicted stem potential. Individual branches representing developmental lineages labeled by canonical markers. **(B,C)** p-Creode trajectories with **(B)** *AXIN2* overlay to represent WNT pathway activity, and **(C)** *MUC2* overlay to represent goblet cell mucin production. Node size represents the proportion of cells, and overlay heat represents mean expression. **(D-G)** Multiplex imaging of representative tissue subtypes (NL – normal, AD – tubular adenoma, HP – microvesicular hyperplastic polyp, SSL – sessile serrated lesion) for **(D)** SOX9 - magenta, **(E)** OLFM4 – magenta, **(F)** CDX2 - brown, and **(G)** MUC5AC - brown. Nuclei are green in D,E, blue in F,G. Scale bar: 100 μm. **(H)** Representative genetic phylogenies inferred through DENDRO, with each tree representing a specimen. Hierarchical clusters demarcated by dotted boxes and cell clusters denoted by leaf node color. Tree height is derived from a beta-binomial adjusted genetic divergence between single cells and shows the evolutionary time before splits in phylogeny.

To further inform the origin of neoplasia between the two polyp subtypes, we assessed the location of abnormal cells by multiplex imaging, since stem cells and non-stem cells occupy distinct compartments of the colonic crypt axis. To validate the differences observed by transcriptomic analysis, we revealed stem cell markers OLFM4 and SOX9 are enriched in ADs at the protein level, and are also strongly expressed at the bottom of normal crypts **(Figure 4D,E; Figure S12A,B)**. These markers are significantly reduced in HPs and SSLs. Nuclear CDX2 immunoreactivity was detected in the normal colon and was maintained in ADs, but was decreased in HPs and absent in SSLs, supporting gastric metaplasia through loss of regional identity **(Figure 4F; Figure S12C)**. Staining for MUC5AC, a marker of metaplastic serrated cells, was absent in normal colon and ADs but enriched in HPs and SSLs **(Figure 4G; Figure S12D)**. Importantly, MUC5AC-positive cells were often observed at the top of the crypt with MUC5AC-negative, normal appearing cells at the crypt bottom, implying a non-crypt origin of serrated polyps. This is further supported by histological evidence that MVHP, candidate precursors to SSLs, have abnormal serrated cells at the surface of the epithelium and not at the crypt bottom **(Figure 1C; Figure S12D,E)**. Only a subset of GCHPs possessed MUC5AC-positive cells, and only at the extreme luminal side of the crypts **(Figure S12E)**, again supporting that abnormal cells originated from the luminal surface. MUC5AC-positive cells were detected in the majority of abnormal crypts in MVHPs and SSLs **(Figure S12F)**, indicating that metaplasia is a homogeneous feature to pre-cancer initiation within each serrated polyp. While MUC5AC staining was largely absent in the normal colon, occasional MUC5AC immunoreactivity was detected at the luminal surface **(Figure S13)**. Increased MUC5AC immunoreactivity was also observed in the colonic epithelia of individuals with ulcerative colitis **(Figure S13)**, suggesting a response to ongoing damage. Given transient damage may occur even in healthy patients, these surface colonic cells may serve as damage-induced metaplastic cells that form the earliest origins of serrated polyps if damage persists.

The shared mutational profiles between populations of neoplastic cells and normal cells can be used to determine cellular origins. Because we profiled normal cells in each polyp by sequencing, we can leverage mutational information from both neoplastic cells and normal cells on a per polyp basis to determine the genetic distance between cells and evaluate shared origins. To approximate the inherited genomic variations from single-cell data, we used DNA-based EvolutionNary tree preDiction by scRNA-seq technOlogy (DENDRO), a phylogenetic reconstruction algorithm using a beta-binomial model to adjust for the inherent sparsity and stochasticity of single-cell gene transcription (Zhou et al., 2020). We examined the genetic divergence between aggregates of transcriptionally similar cells to account for low sequencing depth on shorter reads. Representative DENDRO trees described the genetic heterogeneity found across histological classifications, showing striking differences between the phylogenetic structures of ASCs and SSCs. ASCs clustered with crypt base stem cells and had a shorter genetic distance to these cells compared to SSCs **(Figure 4H; Figure S14)**. In contrast, SSCs demonstrated divergent genetic profiles with stem cells, and, in fact, often clustered with differentiated colonocytes and absorptive progenitors **(Figure 4H; Figure S14)**. Combined, these data are consistent with the hypothesis that ADs arise from the dysregulation of the stem cell compartment whereas SSLs appear to arise from a committed absorptive cell.

### Concordant molecular signatures between CRC and pre-cancer lesions suggest common tumorigenesis pathway

Having investigated the origins of pre-cancerous polyps, we then addressed their relationships to established CRCs. We analyzed three freshly isolated CRCs and compared the results to pre-cancerous polyp-specific cells **(Table S15)**. Microsatellite testing revealed two of the CRCs were MSS (one proximal and one distal) and the third CRC was MSI-H and proximal. As expected, the proximal MSI-H CRC contained mutant *BRAF* and the most total mutations as determined by whole exome sequencing **(Figure S15A)**. The proximal MSS CRC contained a significant number of mutations compared to the distal MSS CRC, perhaps consistent with the association between increased microbial burden within proximal neoplastic lesions and elevated genotoxic damage (Dejea et al., 2014). The distal MSS CRC appeared to follow the conventional tumorigenesis sequence with mutations in *APC*, *KRAS*, *p53*, while the proximal CRC also contained a *APC* driver mutation coupled to other mutations in the Notch and TGFβ pathways **(Figure S15A)**. Histologically, all three CRCs showed invasive adenocarcinoma with cribriform architecture **(Figure S15B,C)**. The MSI-H case showed mucinous features with cancer glands floating in pools of extracellular mucin, consistent with a portion of MSI-H CRC classified as mucinous adenocarcinomas. To address tumor heterogeneity, we performed scRNA-seq and analyzed tumor-specific epithelial cells **(Figure 5A)**. We first applied the CytoTRACE algorithm to predict stem cell potential on a single-cell basis. While ASCs and MSS cancer cells had similar cell distributions of CytoTRACE scores, SSCs and MSI-H cancer cells had divergent skewed distributions in that the cancer cells had a higher inferred stem potential **(Figure 5B; Table S16)**. These results suggest a transition of pre-cancerous metaplastic cells into more aggressive stemlike cancer cells.

**Figure 5:**
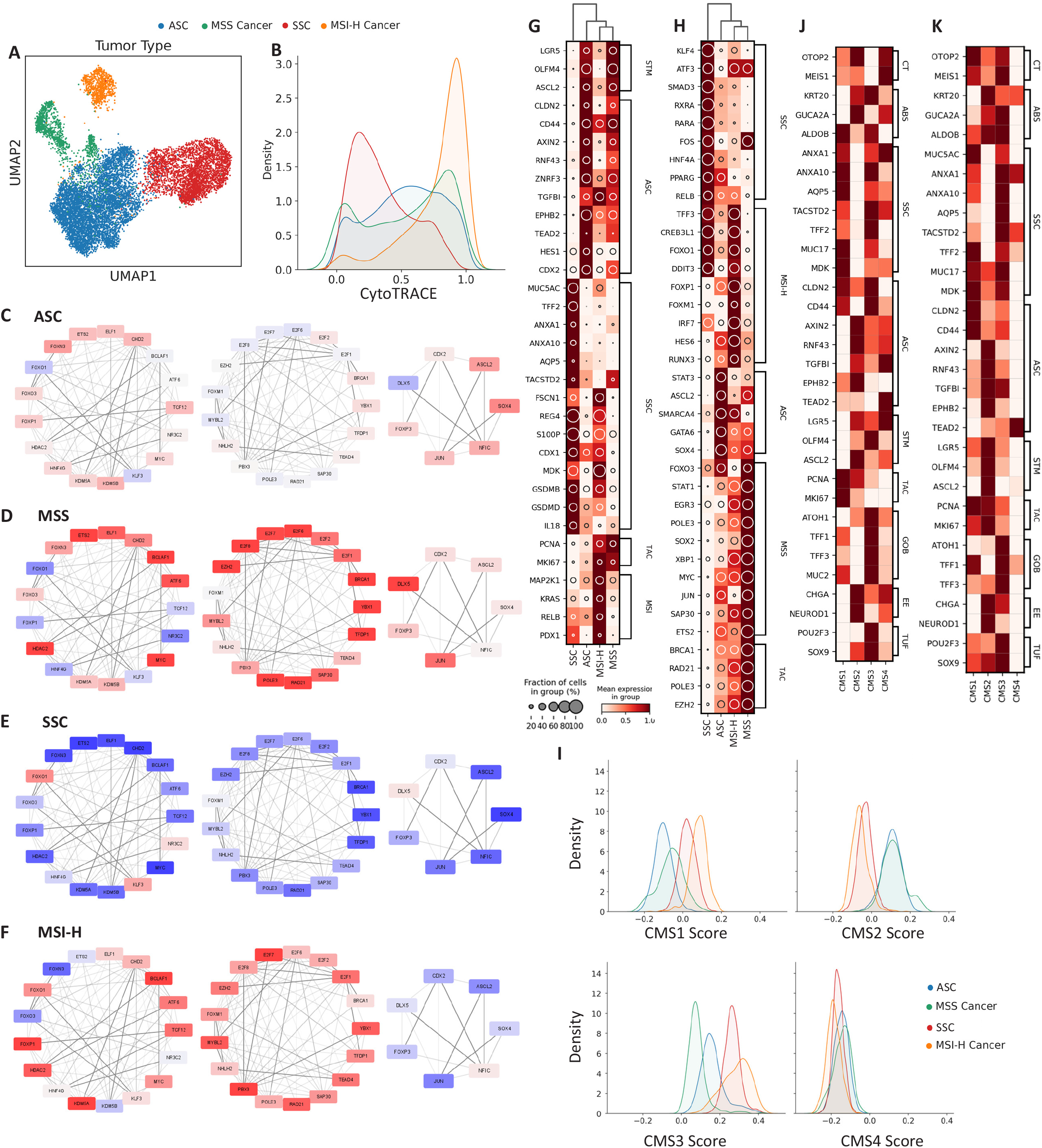
Analysis of CRCs through the lens of pre-cancers. **(A)** Regulon-based UMAP for lesion-specific cells derived from pre-cancerous and cancerous specimens colored by subtypes. **(B)** Density plot of single-cell CytoTRACE scores of different tumor subtypes colored as in A. **(C-F)** TF target subnetworks selected for differential enrichment between molecular pathways specific to tumor subtypes: **(C)** ASC, **(D)**, MSS, **(E)** SSC, and **(F)** MSI-H. Color overlays for each TF node are the average regulon enrichment scores per subclassification and edge opacities are the inferred TF-target weightings. **(G,H)** Heatmap representation of **(G)** gene signatures and **(H)** regulon activities (enrichment scores) on different tumor subtypes, derived from pre-cancer subtyping related to metaplasia or WNT-driven stem cell dysregulation. The mean expression is depicted by heat intensity and prevalence at single-cell level is depicted by the size of the inset circle. **(I)** Density plots of single-cell CMS scoring based on correlation distance to established CMS subtype centroids. Coloring corresponds to coloring depicted in A. **(J,K)** Heatmap representation of gene signatures identified by scRNA-seq of pre-cancers applied to **(J)** bulk RNA-seq of TCGA CRC samples (CMS1, n=8; CMS2, n=26; CMS3, n=10; CMS4, n=19) and **(K)** scRNA-seq of Lee et al. CRC samples (CMS1, n=5; CMS2, n=8; CMS3, n=4, CMS4, n=6). The mean expression is depicted through color intensity.

We then examined gene expression and TF-driven co-regulatory networks of these cancers in comparison to pre-cancerous polyps **(Figure 5C-F; Figure S16)**. Several parallels could be drawn between the profiles of ASCs and MSS cancer cells, primarily supporting the activation of a regenerative program of crypt base stem cells. Genes associated with crypt base stem cells, such as *LGR5* and *OLFM4*, and a subnetwork of regenerative regulons, such as *ASCL2* and *SOX4*, were further increased in MSS cancer cells compared with ASCs **(Figure 5C,D,G,H; Figure S16D,E)**. The regenerative signature was absent in both MSI-H cancer cells and SSCs **(Figure 5E-H)**. Gastric metaplastic genes (*MUC5AC*, *TFF2*, *ANXA1*, *ANXA10*) and environmental communication signatures, such as TGFβ (SMAD3 regulon) and RA signaling (RXRA and RARA regulons), were decreased in MSI-H cancer cells compared with SSCs **(Figure 5G,H; Figure S16F,G)**, suggesting an evolving independence of MSI-H tumor cells from metaplastic cells for maintaining growth at different stages of progression. Strikingly, the major subnetwork describing a microbiota-driven immune response in SSCs was suppressed in MSI-H cancer cells with only the CREB-related and IRF7 regulons remaining active **(Figure S16F,G, S17; Table S17-18)**. However, MSI-H cancer cells contained remnants of gastric metaplasia **(Figure 5G)**, including *MUC5AC* and *REG4* expression and loss of *CDX2* expression. The metaplasia-related program was expressed at levels lower than SSCs but still higher than ASCs and MSS cancer cells, implicating a common origin with SSCs. This program was accompanied by an enrichment of a WNT pathway activation subnetwork **(Figure 5E, F)**. We found activating mutations in components of the WNT pathway other than *APC*, including *APC2*, *FAT2*, and *CELSR2* mutations **(Figure S15A)**. This observation was buttressed by TCGA exome sequencing data showing enrichment of non-APC WNT pathway gene mutations in MSI-H CRC (*APC2*, *RNF43*, *AXIN2*, *LRP1B*, *LRP6*, *TCF7L2*). **(Figure S18)** (Muzny et al., 2012). A major common difference between pre-cancer and cancer cells for both subtypes was the expansion of actively cycling cells with enrichment of genes (*PCNA* and *MKI67*) and regulons (BRCA1, RAD21, POLE3, and EZH2) involved in DNA synthesis and repair **(Figure 5C-F)**. These results demonstrate the consistent role of a stem cell program in the conventional tumorigenesis pathway from ADs to MSS cancer. Interestingly, they also suggest MSI-H cancers no longer require metaplasia-related genes, as seen in serrated lesions, possibly due to acquired mutations in the WNT pathway that fuel tumor progression.

To further support the common origin between pre-cancer and cancer, we applied established bulk transcriptional models of consensus molecular subtypes (CMS) to score ASCs, SSCs, and cancer-specific cells **(Figure S19A)**. The CMS framework classifies four cancer subtypes by transcriptomic signatures, each of which are associated with defined molecular and clinical features (Guinney et al., 2015). CMS scoring, described as a single-sample predictor, measured the median distance between transcriptional centroids of each CMS and each cell’s transcriptome, generating a distribution of scores for each tumor type **(Figure S19B)**. Strikingly, ASCs shared comparable score distributions with the MSS tumors, which are predicted to be CMS2 **(Figure 5I; Figure S19C; Table S19-S22)**. This CMS2 classification features canonical WNT signaling pathway dysregulation through *APC* mutations, resulting in an expanded stem cell population, consistent with the gene network modules we observed in ASCs. In contrast, the MSI-H cancer scored relatively high for CMS1 and CMS3 transcriptional signatures, consistent with immunogenicity (IRF7 and CREB regulons, *S100P* and gasdermin genes) and RAS pathway activation (FOXM1, FOXO1, FOXP1, RUNX3, MAF regulons) (Chi et al., 2009; Feng et al., 2011; Liao et al., 2018), respectively **(Figure 5I; Figure S19C)**. In the case of SSCs, we observed a similar trend with enriched CMS1 and CMS3 signatures and a depleted CMS2 signature **(Figure 5I; Figure S19C)**. None of the examined polyp cells had strong enrichment in CMS4 epithelial-to-mesenchymal transition scores **(Figure 5I; Figure S19C)**, consistent with their identity as early-lesion cells and previously reported absence of CMS4 in ADs (Komor et al., 2018). We provided further support of these distinguishing profiles by examining 63 bulk RNA-seq datasets from the TCGA and 23 scRNA-seq datasets from Lee *et al*. annotated with CMS subtypes **(Figure S20)** (Lee et al., 2020; Muzny et al., 2012). Considering bulk profiles and only tumor cells from scRNA-seq, we found the consistent expression of a gastric metaplastic signature, including genes such as *MUC5AC*, *ANXA1*, *ANXA10*, *AQP5*, *TACSTD2* (TROP2), *TFF2*, and *MUC17* in CMS1 and CMS3 CRCs. These genes were depleted in CMS2. In parallel, genes indicative of WNT pathway upregulation, such as *LGR5*, *OLFM4*, *ASCL2*, *AXIN2*, *RNF43*, and *EPHB2*, were depleted in CMS1 **(Figure 5J,K; Figure S20)**. The consistency in CMS classification between ASCs and MSS cancer cells, and between SSCs and MSI-H cancer cells, supports their molecular similarity and implicates possible progression routes from pre-cancerous polyps to cancer.

To examine whether cellular heterogeneity increases with progression to CRC, we used multiplex immunostaining and whole slide scanning to image large CRCs. The two MSS CRCs exhibited a uniform lack of MUC5AC expression but positive CDX2 expression **(Figure 6A,B; Figure S21A,B)**. In validation of scRNA-seq results, the stem cell markers OLFM4 and SOX9 were highly expressed throughout the tumor **(Figure 6B)**, confirming further expansion of aberrant stem cells in these tumors. The MSI-H CRC displayed considerable heterogeneity in line with our transcriptomic results demonstrating the loss of metaplastic characteristics and simultaneous WNT signaling activation. MUC5AC, a marker of gastric metaplasia, was expressed highly but only in certain regions of the tumor, unlike in SSLs where almost all crypts were MUC5AC-positive **(Figure 6C,D)**. CDX2 was also heterogeneously expressed with positive staining in regions with low MUC5AC expression **(Figure 6D; Figure S21C)**. In these same regions, OLFM4 was highly expressed, demonstrating mutual exclusivity between stem-like cells and metaplastic cells **(Figure 6E,F)**. However, SOX9 was overexpressed in the tumor in both MUC5AC high and low regions, suggesting all cells gained some level of stem-like characteristics upon transformation **(Figure 6F; Figure S21D)**. MDK, a gene expressed transiently in early colonic development, also exhibited heterogenous staining in this specimen **(Figure S21E)**. To validate this observed heterogeneity, we focused our scRNA-seq analysis on the MSI-H cancer cells. Positive *MUC5AC* and *MSLN* expression, coupled to loss of *CDX2* expression, can be used to distinguish metaplastic cells from *LGR5*/β-catenin-expressing, WNT-driven stem cells within the same tumor **(Figure 6G)**. The WNT-driven stem cells were enriched in cell cycle gene expression, consistent with the increased proliferative capacity of CRCs identified in our previous analysis **(Figure 6G)**. We confirmed these findings in additional CRCs. Using MLH1 staining, we inferred the microsatellite status of specimens on a CRC tissue microarray. MLH1-high cancers (presumably MSS) had uniform expression of CDX2 without MUC5AC, whether they were well-differentiated (CRC1) or less differentiated (CRC2) **(Figure 6H)**. CRC3 had uniformly low MLH1 staining (presumably MSI-H) and expressed MUC5AC without CDX2. CRC4 was heterogeneous in MLH1 staining, and had areas of CDX2^low^/MUC5AC^high^ expression and areas with CDX2^high^/MUC5AC^low^ expression. These results suggest ADs progress to MSS cancers through the same stem cell expansion mechanism, while MSI-H cancers actually gain stem cell characteristics in a background of metaplasia leading to significant cellular heterogeneity.

**Figure 6:**
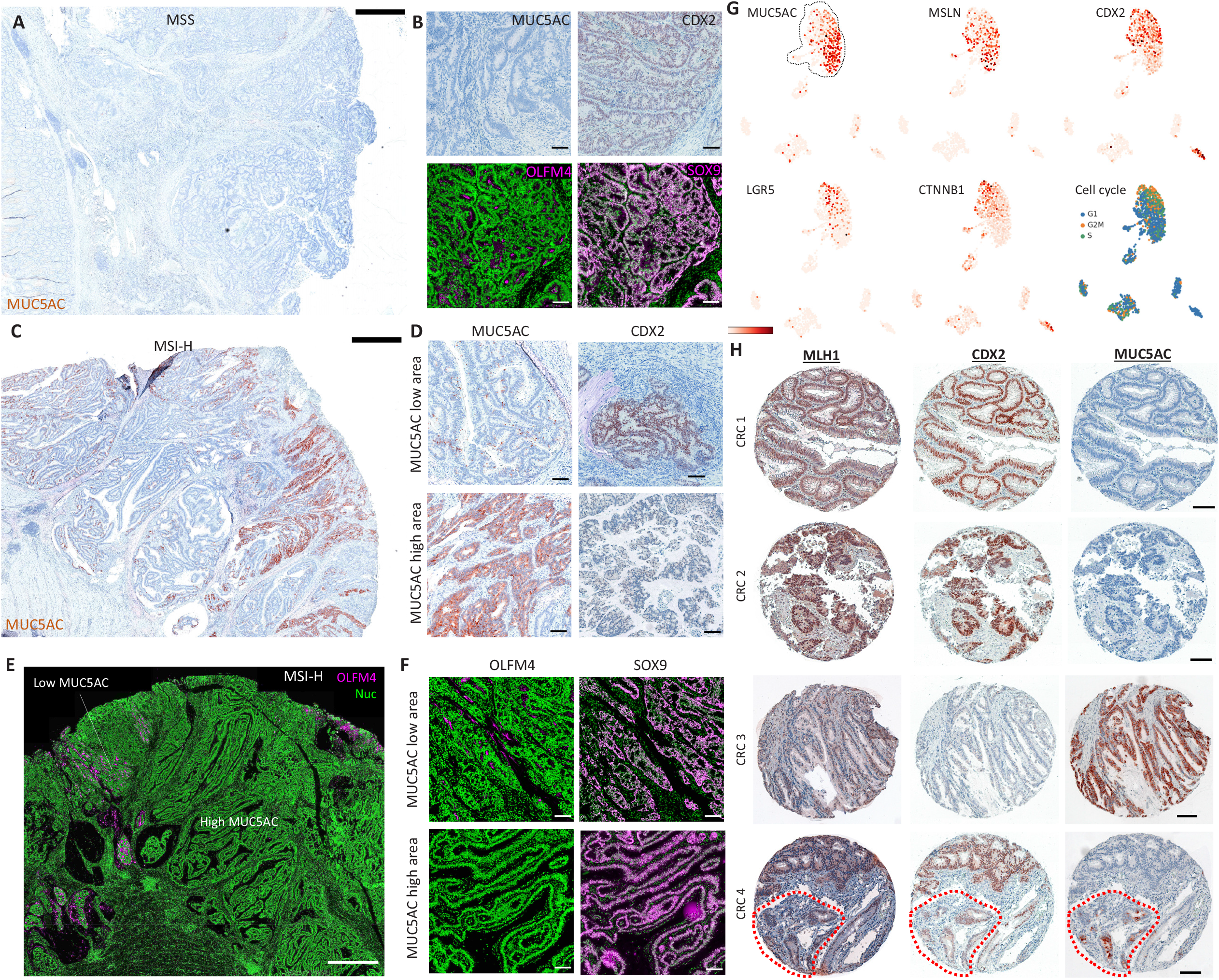
Heterogeneity of CRCs with metaplastic and stem-like features. **(A)** Low magnification view (scale bar: 1mm) of MUC5AC staining, and **(B)** high magnification view (scale bar: 100 μm) of a MSS CRC with various stains for proteins. **(C)** Low magnification view of MUC5AC staining of a MSI-H CRC. **(D)** High magnification view of a MUC5AC high and MUC5AC low area for metaplasia markers of the CRC in C. **(E,F)** Same as in C,D but staining for stem cell markers. **(G)** UMAP of scRNA-seq data of the MSI-H CRC overlaid with various metaplasia and stem cell markers, as well as with cell cycle signatures. **(H)** Staining for various metaplasia markers from 4 CRCs from a tissue microarray. Red outline represents the MLH1 low area in CRC 4. Scale bar: 100 μm. MUC5AC, CDX2, MLH1 = brown; OLFM4, SOX9 = magenta; Nuclei = green for IF and blue for IHC.

### Serrated polyps contain an alternate immune microenvironment from conventional adenomas preceding hypermutation

High microsatellite instability has been hypothesized to induce a cytotoxic microenvironment in CRCs conferring susceptibility to immunotherapy, although unidentified factors likely contribute here (Le et al., 2017; Mlecnik et al., 2016). From mutational analysis, serrated polyps did not exhibit hypermutation **(Figure 2)**, while all analyzed MSI-H cancers were hypermutated **(Figure S15A and S18)**. Our datasets thus provided an opportunity to determine whether cellular origin and pathway differences alone predetermine the tumor microenvironment preceding hypermutation. From scRNA-seq data of the non-epithelial component, we identified T cells, B cells, plasma cells, mast cells, myeloid cells, fibroblasts and endothelial cells based on differential marker expression **(Figure 7A-C)**. Microenvironmental composition was markedly different between tumor subtypes **(Figure 7A,D)**. Both conventional and serrated polyps had abundant T cells, though serrated polyps had a significantly larger ratio of T cells to myeloid cells compared with ADs **(Figure 7A,D)**. B cells were suppressed in both polyp types compared to normal mucosa, although plasma cells were not reduced and were more abundant in ADs than SSLs. Surprisingly, mast cells were also overrepresented in ADs. MSS cancers had fewer total immune cells than the MSI-H cancer. However, the MSI-H cancer had reduced proportions of T cells and increased proportions of myeloid cells compared to pre-cancer. Similar to serrated polyps, plasma cells and mast cells were reduced in MSI-H cancer. There was no significant difference in the small numbers of fibroblasts and endothelial cells among tumor subtypes.

**Figure 7:**
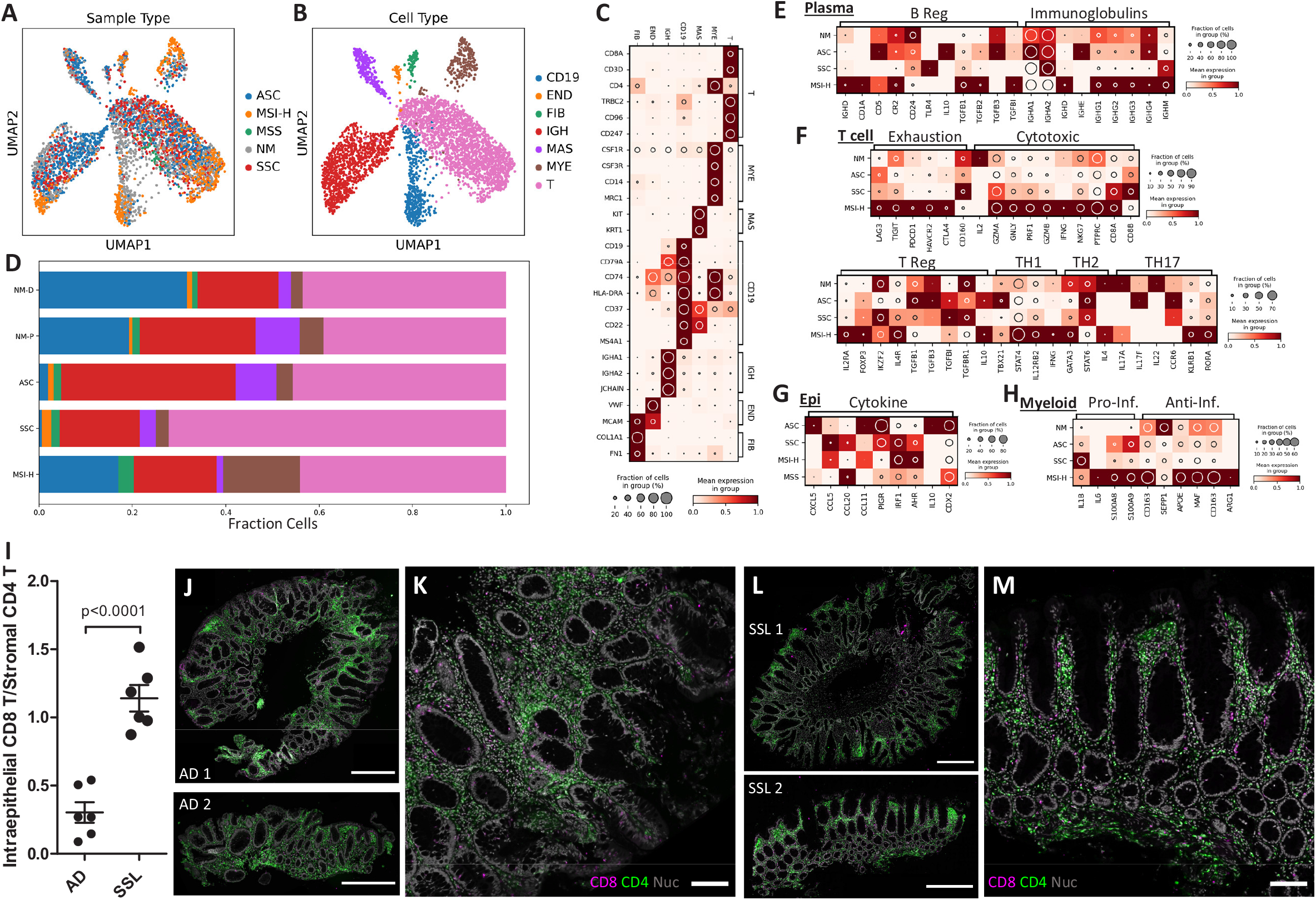
The immune microenvironment of different tumor subtypes. **(A,B) Regulon-based** UMAP representation of non-epithelial cell populations overlaid with **(A)** tumor subtype, and **(B)** cell type defined by Leiden clustering and marker genes. **(C)** Heatmap representation of marker genes defining each cell type per cell cluster. **(D)** Proportion of cell types, following the same color scheme in A, of each tumor subtype. **(E-H)** Heatmap representation of functional genes expressed by each tumor subtype in the **(E)** plasma cell, **(F)** T cell, **(G)** epithelial cell (cytokines), and **(H)** myeloid cell populations. The mean expression is depicted by heat intensity and prevalence at single-cell level is depicted by the size of the inset circle. **(I)** Quantification of the ratio of intraepithelial CD8+/CD4+ T cells in the stroma, normalized by area of each compartment. n=6 fields of view from different polyps. **(J)** Low magnification (Scale bar: 1mm) and **(K)** high magnification (Scale bar: 100μm) views of T cell distribution in different samples of tubular ADs. **(L)** Low magnification (Scale bar: 1mm) and **(M)** high magnification (Scale bar: 100μm) views of T cell distribution in different samples of SSLs.

Pre-cancer resident plasma cells express IgA, which is transported across the epithelium into the lumen by PIGR. *PIGR* was expressed highly in pre-cancer cells compared to both MSS and MSI-H cancer cells, suggesting that the normal gut humoral response is relatively intact in early lesions. IgA2, which has fewer sialic acid N-glycans than IgA1 and triggers a larger immune response (Steffen et al., 2020), was enriched in a subset of HP plasma cells, consistent with higher immunogenicity against microbes **(Figure 7E)**. A subset of tubular AD plasma cells expressed a high level of IgE, an immunoglobulin involved with allergic reactions, consistent with increased mast cells detected in these lesions. In contrast, cancer plasma cells did not express IgA but instead expressed IgG, the major immunoglobulin subtype in blood plasma arising from the spleen and lymph nodes. Furthermore, these cells expressed regulatory B cell markers, suggestive of immunosuppression.

When considering T cells, a dichotomy was observed between ADs and serrated polyps. AD T cells expressed a CD4+ Th1/Th17 inflammatory signature, while serrated polyp T cells expressed a strong CD8+ cytotoxic T cell phenotype with some Th17 markers **(Figure 7F)**. Regarding signaling receptor expression, serrated polyp T cells resembled normal colon mucosa T cells, the majority of which were cytotoxic intraepithelial lymphocytes **(Figure S22A,B)**. In contrast, MSI-H cancer T cells expressed markers of all subtypes, including a strong CD8+ signature **(Figure 7F)**. However, both activation and exhaustion/checkpoint markers were highly expressed, indicating T cell dysfunction. Interestingly, regulatory T cells (Tregs) expressing *FOXP3* were found only in the MSI-H cancer, consistent with active immunosuppression. Immunosuppression of MSI-H cancer, and to some degree ADs, was supported by expression of the regulatory cytokine *IL-10* by T cells from both tumor types but not serrated polyps. Epithelial cells in ASCs expressed chemokines attracting monocytes (*CXCL5*, *CCL11*), while SSCs expressed cytokines attracting lymphocytes (*CCL20*, *CCL5*), further supporting the dichotomy between innate versus adaptive inflammation (Hieshima et al., 1997; Nelson et al., 2001) **(Figure 7G)**. Pre-cancers had fewer myeloid cells than the MSI-H cancer, where they expressed both inflammatory and suppressive genes, which is characteristic of a MDSC-like phenotype **(Figure 7H)**. These data illustrate differences in the tumor microenvironment preceding hypermutation and define the innate versus adaptive inflammatory nature of the two polyp classes.

We used multiplex imaging to evaluate the geographical differences in the T cell compartment between conventional and serrated tumor subtypes. The T cell compartment characteristics are important for predicting immunotherapy responses (Mlecnik et al., 2016). SSLs had a higher ratio of intraepithelial CD8+/CD4+ T cells than ADs, confirming our transcriptomic results **(Figure 7I)**.

Of note, the spatial distributions of T cells between polyp subtypes were different. ADs had CD4+ T cells distributed throughout the tumor stroma with a slight bias towards the pericryptal space **(Figure 7J,K; Figure S22C,D)**. In contrast, CD4+ T cells in SSLs were concentrated at the luminal surface **(Figure 7L,M; Figure S22E,F)**, which coincided with the surface location of these lesions and sporadic MUC5AC+ cells in the normal epithelium. CD8+ T cells were detected in the epithelial compartment of SSLs, supporting the similarity of these cells to IELs. The ratio of T cell subtypes in CRCs were similar to pre-cancerous polyps mirroring the stem-like versus metaplastic properties of epithelial cells. In line with transcriptomic results, MSS cancers had substantially fewer T cells than other tumor types, including lower CD8+ cells throughout the tumor **(Figure S23A-D)**. In contrast, the MSI-H cancer had a heterogeneous distribution of CD8+ cells with enrichment in the epithelial compartments of MUC5AC-positive metaplastic areas and a reduced number in OLFM4-positive stem cell-like areas **(Figure S23E,F)**. These results further strengthen the relationship between the metaplastic program of tumorigenesis and the cytotoxic immune microenvironment.

## Discussion

We present our initial multi-omic analysis of the two major subclasses of pre-malignant polyps of the colon, ADs and serrated polyps including SSLs and HPs. Our findings support that ADs arise from WNT pathway dysregulation in the stem cell compartment. In marked contrast, we have discovered that SSLs originate from a metaplastic process, likely arising from committed absorptive cells.

Both the etiologies and molecular events associated with tumor initiation and progression are fundamentally different between these two subclasses of pre-malignant tumors. The initiating step in sporadic ADs is activation of the WNT pathway, most often due to loss-of-function mutations in *APC*, and tumors progress with cell-autonomous accretion of additional genetic events in the stem cell compartment. Results from genetically engineered mouse models have provided support for stem cells as the cell-of-origin for colorectal adenomas. Genetic elimination of one or both *Apc* alleles in Lrig1 and Lgr5 stem cells, respectively, predictably results in adenomas (Powell et al., 2012; Barker et al., 2007). From our human data, we observed expansion of aberrant cells that resembled crypt base stem cells in ADs with highly activated WNT signaling, self-renewal, and regenerative programs. Inferring the developmental trajectory and genetic phylogeny of these cells supports their derivation from the crypt base stem cell lineage. From exome sequencing analysis, activating mutations in the proto-oncogene *KRAS* were found to increase from tubular ADs to histologically more advanced tubulovillous ADs. We also found mutations in *APC2*, the functional ortholog of *APC*, and *CTNNB1* that overlapped with *APC*, consistent with models in which early tumors are polyclonal but evolve to multiclonality as dominant clones emerge (Thliveris et al., 2013). A small subset of conventional ADs progress to CRC, and from our cancer analysis, we infer that the most likely route of progression of these tumors is into MSS and CIN (CMS2).

The origin of the less frequent and much less studied SSLs is unknown. A central finding of our study is that SSLs and associated serrated polyps originate from a pyloric-type gastric metaplasia. Histopathologically, metaplasia is defined as a process in which normal differentiated lineages are replaced by cell lineages ectopic to the native organ. The metaplastic program we observed arises in response to damage in the colonic epithelium, which activates a regenerative paradigm to marshal the expression of reparative mucous-secreting lineages. This process is characterized by the expression of genes normally reflective of the pyloric glands of the distal stomach (Goldenring, 2018). Metaplastic programs have been documented in the stomach (spasmolytic polypeptide-expressing metaplasia, SPEM) (Nam et al., 2010; Schmidt et al., 1999), pancreas (acinar to ductal metaplasia, ADM) (Means et al., 2005), and small intestine (ulcer-associated lineage, UACL) (Meditskou et al., 2020; Thorsvik et al., 2019; Wright et al., 1990). In these cases, metaplastic reparative mucous cells arise from transdifferentiation of differentiated zymogenic cells (Jones et al., 2019b; Means et al., 2005; Nam et al., 2010). The cell origin of gastric metaplasia and SSCs is less clear in the colon. In SSLs, we observed mis-expression of genes found in other endodermal organs, including *MUC5AC*, *ANXA1*, *MSLN*, *ANXA10*. *AQP5* has recently been noted as a marker of deep antral gland cells and antral gland stem cells (Tan et al., 2020). While cells from serrated polyps express a transcriptomic identity resembling the absorptive lineage, they aberrantly produce mucins such as MUC2 without expressing the intestinal master secretory cell TF ATOH1. Metaplasia in the stomach and pancreas can evolve into more fixed metaplastic variants (intestinal metaplasia and PANIN, respectively). It is therefore relevant that SSCs display upregulation of *TACSTD2* (*TROP2*), which is prominently upregulated in intestinal metaplasia and dysplasia in the human stomach (Riera et al., 2020).

Expression of *CDX2*, a key homeobox gene for assigning hindgut regional identity during embryogenesis, is lost in serrated polyps compared to adjacent normal mucosa and ADs. In fact, there was a striking inverse correlation between MUC5AC and CDX2 staining in SSLs. In the postnatal gut, attenuated *Cdx2* expression in mice results in a rostral shift of tissue identity with expression of markers of gastric identity (Balbinot et al., 2018). We also observed increased expression of *MDK*, a gene expressed in early colonic development. We propose a model in which an unspecified damage in the proximal colon initiates a regenerative process resulting in reduced *CDX2* expression, gastric metaplasia, and reversion to a more primitive developmental state. In this model, a more primitive state may be susceptible to acquiring *BRAF* mutations. We observed SSLs that lack BRAF mutations, MSI or CIMP, in contrast to work by Sakamoto et al. (Sakamoto et al., 2017).

It is important to distinguish metaplastic loss of regional identity from dedifferentiation of cells into crypt base stem cells (Buczacki et al., 2013; Van Es et al., 2012; Schonhoff et al., 2004; Tetteh et al., 2016), as the latter still retains the identity of the original organ. Different types of damage may incite different mechanisms of regeneration. Minor damage to the gland surface is usually repaired through induced hyperplasias. In contrast, the development of metaplasia is associated with more severe full thickness mucosal injuries (Engevik et al., 2016). What incites the metaplastic program is unknown, although it is attractive to consider the effect of microbiota-induced cytoxicity towards surface epithelial cells. We observed shared lineages between SSCs and committed absorptive epithelial cells. MVHPs, reported precursors to SSLs, appeared to be initiated from the surface epithelium and not the crypt base in histopathological analysis. We also observed occasional expression of MUC5AC on the surface of adjacent normal epithelium, and T cells were spatially distributed towards the luminal surface. Inflammation is thought to play a critical role in various metaplastic processes, although it is not absolutely necessary for initiation of transdifferentiation in SPEM (Goldenring et al., 2000; Petersen et al., 2014, 2018). Nevertheless, we observed a strong inflammatory signature in serrated polyps. While cancers of the serrated pathway are associated with CIMP and MSI-H, serrated polyps in our study did not exhibit MSI-H or hypermutation. However, they did exhibit an immune response skewed towards cytotoxicity versus a helper response when compared to ADs. MSI-H cancers have recently received attention due to their susceptibility to checkpoint immunotherapy, thought to be dependent on the neoantigen load conferred by hypermutation. Here, we support the origin of early tumorigenesis as an additional determinant of the tumor microenvironment preceding methylation and hypermutation in these tumors.

## Supporting information

Key Resources Table

## Acknowledgements

The authors wish to thank the study participants who chose to participate in the research and the many individuals who contributed to the work presented in this manuscript. Some of the contributors include Reid Ness, Yu Shyr, Harvey Murff, David Pocalyko, and Joke Reumers, Harrison Kiang, and Eric Eisenberg. We apologize in advance to those we have failed to acknowledge due to space constraints. This work is supported in part through the Human Tumor Atlas Network (HTAN). This study was supported by US National Institutes of Health (NIH) grants U2CCA233291 (to RJS, KL, MJS), R01CA97386 (to WZ), P50CA236733 (to RJC), R01DK103831 (to KL), P50CA950103 (to RJC), K07CA122451 (to MJS), T32LM012412 (in support of BC), and R35CA197570 (to RJC). Funding for the bulk RNA sequencing of polyps was provided by JANSSEN Research & Development (to MJS). Study activities were conducted in part by the Survey and Biospecimen Shared Resource (P30CA68485), the Tissue Pathology Shared Resource (P30CA068485, U24DK059637), the Digital Histology Shared Resource, the NCI Cooperative Human Tissue Network (CHTN) Western Division (UM1CA183727), the CHTN at the University of Virginia, the Vanderbilt Technologies for Advanced Genomics, and REDCap (UL1TR000445). The content of this paper is solely the responsibility of the authors and does not necessarily represent the official views of the National Cancer Institute or the NIH. A portion of the participants were studied as the result of resources and the use of facilities at the Veterans Affairs Tennessee Valley Healthcare System.

## Author Contributions

Conceptualization (MKW, WZ, CLS, RJC, MJS, KL); Data curation (BC,XZ, CNH, LB, MJS, KL; Formal analysis (BC, ETM, MARS, YX, CNH, SV, QL, KL); Investigation (BC, ETM, AJS, XZ, ANSS, NOM, QS, JLD, CNH, YZ, FR, LB, QC, JLF, JTR, TS, WJH, RJC, MJS, KL); Methodology (BC, ETM, AJS, XZ ANSS, JLF, RJC, MJS, KL); Project administration (ETM, AJS, XZ, QL, RJC, MJS, KL); Resources (MKW, WZ, JRG, JTR, WJH, QL, RJC, MJS, KL); Software (BC, CNH, QL, KL); Supervision (RJC, MJS, KL); Validation (BC, CNH); Visualization (BC, MARS, ANSS, QS, CNH, SV, WJH, QL, RJC, MJS, KL); Writing-original draft (BC, WJH, RJC, MJS, KL); Writing-reviewing & editing (BC, ETM, AJS, MARS, XA, ANSS, NOM, QS, JLD, YX, CNH, YZ, MKW, FR, LB, WZ, QC, CLS, JRG, JLF, SV, JTR, TS, WJH, QL, RJC, MJS, KL)

## Supplementary Materials

**Figure S1: Cellular and histological features of pre-cancers and normal colons. (A,B)** Normal colonic specimens for single-cell analysis categorized into **(A)** proximal and **(B)** distal locations corresponding to their collection site in the colon alongside patient metadata. Stacked bar plots describe epithelial cellular distribution of each specimen with colors representing cell types as defined by Leiden clustering and marker gene detection. ABS-absorptive cells, ASC-adenoma-specific cells, CT-crypt top colonocytes, EE-enteroendocrine cells, GOB-goblet cells, STM-stem cells, SSC-serrated-specific cells, TAC-transit amplifying cells, TUF-tuft cells. **(C)** More examples of histology of each polyp subtype, HP-hyperplastic polyps, SSL-sessile serrated lesions, TVA-tubulovillous adenomas, TA-tubular adenomas. Scale bar: 100 μm.

**Figure S2: Single-cell data as a function of sample heterogeneity characteristics. (A,B)** UMAP co-embedding of uncorrected epithelial scRNA-seq count data generated from patient tissue, and colored by **(A)** HTAN patient ID, and **(B)** histological subtype, clusters, location of polyp, predicted cell cycle phase, patient gender, and patient race. **(C,D)** UMAP co-embedding of SCENIC-corrected epithelial scRNA-seq data overlaid with the same information as in A and B.

**Figure S3: Cell type identification from scRNA-seq**. UMAP co-embedding of SCENIC-corrected epithelial cell scRNA-seq data overlaid with subpopulation-specific genes.

**Figure S4: Low magnification of view of specimen histologies.** Scale bar: 100 μm.

**Figure S5: Stem characteristics of pre-cancer cells. (A)** Violin plot representation of log-normalized expression of genes involved in the TGF-β pathway in ASC or SSC populations. **(B)** Density plot of single-cell CytoTRACE scores calculated for cells in each cell cluster. **(C)** UMAP embedding of epithelial cell scRNA-seq data overlaid with CytoTRACE scores. Red outlines demarcate cell populations specific to pre-cancer specimens. **(D)** Bar plot depicting genes with the 20 highest and lowest correlations to CytoTRACE scores.

**Figure S6: Heterogeneity of adenoma-specific cells. (A-C)** UMAP embedding of heterogeneous ASCs overlaid with **(A)** Leiden subclusters, **(B)** CytoTRACE scores, and **(C)** normalized expression values of key genes. **(D-F)** GSEA performed on ASC subpopulations depicted by Manhattan plots of significant gene sets enriched in ASC subcluster **(D)** 0, **(E)** 1, and **(F)** 2. Datapoints represent log(p-values) across six GSEA databases: GO:MF (Gene Ontology: Molecular Function), GO:BP (Gene Ontology: Biological Process), GO:CC (Gene Ontology: Cellular Process), KEGG (Kyoto Encyclopedia of Genes and Genomes), REAC (Reactome), and WP (WikiPathway). Size of datapoints represent the number of enriched genes in a gene set.

**Figure S7: Pathway differences between lesion-specific cell populations and normal cells. (A,B)** GSEA Manhattan plots of significant gene sets enriched in **(A)** ASCs, and **(B)** SSCs. **(C,D)** Manhattan plots of enrichment analysis performed using regulon activities in **(C)** ASCs and **(D)** SSCs. Datapoints represent log(p-values) across six GSEA databases: GO:MF, GO:BP, GO:CC, KEGG, REAC, and WP. Size of datapoints represent the number of enriched genes in a gene set.

**Figure S8: Heterogeneity of serrated-specific cells. (A-C)** UMAP embedding of heterogeneous SSCs overlaid with **(A)** Leiden subclusters, **(B)** CytoTRACE scores, and **(C)** normalized expression values of key genes. **(D-G)** GSEA performed on SSC subpopulations depicted by Manhattan plots of significant gene sets enriched in SSC subcluster **(D)** 0, **(E)** 1, **(F)** 2, and **(G)** 3. Datapoints represent log(p-values) across six GSEA databases: GO:MF, GO:BP, GO:CC, KEGG, REAC, and WP. Size of datapoints represent the number of enriched genes in a gene set.

**Figure S9: TF network analysis of pre-cancers. (A)** Subnetworks of the larger TF target network in Figure 3D and annotated by the same coloring scheme. **(B-D)** Average TF-regulon enrichment score, with red being enrichment and blue being depletion for each subnetwork in **(B)** normal cells, **(C)** ASCs, and **(D)** SSCs. **(E-H)** Further breakdown of these networks by histological subtypes for **(E)** tubular adenomas, **(F)** tubulovillous adenomas, **(G)** sessile serrated lesions, **(H)** and hyperplastic polyps.

**Figure S10: Transcriptomic analysis of pre-cancers.** Heatmap representation of gene signatures identified by scRNA-seq of pre-cancers applied to bulk RNA-seq samples of various pre-cancer subtypes (n=36 tubular, 22 tubulovillous, 8 SSLs). Gene expression of each sample is depicted through heat intensity.

**Figure S11: Trajectory inference from pre-cancer single-cell data. (A)** p-Creode trajectories inferred from epithelial scRNA-seq data from the study. Overlay of genes relevant to developmental lineages. Node size represents the proportion of cells, and overlay color represents mean expression. **(B)** Stacked bar plot describes cellular distribution of histological subtype across p-Creode branches with colors representing cell lineages.

**Figure S12: Stem cell and metaplastic programs in conventional and serrated polyps.** Multiplex imaging of representative tissue subtypes (TA – tubular adenoma, TVA - tubulovillous adenoma, HP – microvesicular hyperplastic polyp, SSL – sessile serrated lesion) for **(A)** SOX9 - magenta, **(B)** OLFM4 – magenta, **(C)** CDX2 - brown, and **(D)** MUC5AC - brown. White outline demarcates HP location. **(E)** MUC5AC staining for a progressive series of serrated polyps (GCHP – goblet cell rich hyperplastic polyp, MVHP – microvesicular hyperplastic polyp, SSL – sessile serrated lesion). **(F)** Low magnitude view of MVHPs and SSLs for MUC5AC (brown) staining. Nuclei are green in A,B, blue in C-F. Scale bar: 100 μm.

**Figure S13: Abnormal cells in the normal colonic epithelium.** MUC5AC staining of nonneoplastic colonic epithelium: normal, tumor adjacent, and ulcerative colitis. Brown = MUC5AC, Blue = nuclei. Scale bar: 100 μm.

**Figure S14: Phylogeny inference from pre-cancer single-cell data. (A-D)** More example hierarchical phylogeny created using DENDRO-inferred genetic distance created on a per specimen basis, for **(A)** tubular, **(B)** tubulovillous, **(C)** sessile serrated, and **(D)** hyperplastic polyp subtypes. Hierarchical clusters demarcated by dotted boxes and cell clusters denoted by leaf node color. Tree height is derived from a beta-binomial adjusted genetic divergence between single cells and shows the evolutionary time before splits in phylogeny.

**Figure S15: Features of CRCs by subtype. (A)** Heatmap representation of the mutational landscape of 3 CRCs by exome sequencing, formatted as in Figure 2. **(B,C)** Histology of each cancer at **(B)** low and **(C)** high magnification views of the corresponding colored insets. Scale bar: 1mm.

**Figure S16: Combined TF network analysis of pre-cancers and CRCs. (A)** TF target network created from pre-cancer and cancer cells, broken down into subnetworks by Leiden subgraph clustering (color). Nodes represent individual TFs and edge opacity denotes similarities in transcriptional factor targets. **(B)** Heatmap depicting average regulon enrichment scores per subtype of each subnetwork. **(C)** Subnetworks of the larger TF target network in A and annotated by the same coloring scheme. Average TF-regulon enrichment score, with red being enrichment and blue being depletion for each subnetwork in **(D)** ASCs, **(E)** MSS cancer cells, **(F)** SSCs, and **(G)** MSI-H cancer cells.

**Figure S17: Pathway differences between cancer cell populations and pre-cancer cell populations. (A-B)** GSEA performed on cancer cell populations versus other pre-cancer-specific cell populations (ASCs and SSCs) using regulon activities. Manhattan plots of significant gene sets enriched in **(A)** MSI, and **(B)** MSS cancer cells. Datapoints represent log(p-values) across six GSEA databases: GO:MF, GO:BP, GO:CC, KEGG, REAC, and WP. Size of datapoints represent the number of enriched genes in a gene set.

**Figure S18: Genomic landscape of TCGA CRCs.** OncoPrinter plot visualizing genomic tracks for 276 assayed TCGA CRC samples. Each row represents unsorted mutational status of key pathway genes. Each column represents an individual sample with hypermutation status noted.

**Figure S19: Pre-cancer relationship with CRCs. (A)** Regulon-based UMAP overlaid with histologically-derived cancer and pre-cancer subtypes. **(B)** UMAP constructed from bulk RNA-seq of TCGA CRC samples colored by CMS subtype, showing distinct molecular profiles. **(C)** UMAP from overlaid with the median correlation distance to single sample predictor centroids derived from CMS1, CMS2, CMS3, and CMS4 subtypes.

**Figure S20: Transcriptomic analysis of CRCs. (A-B)** Heatmap representation of gene signatures identified by scRNA-seq of pre-cancers applied to **(A)** bulk RNA-seq of TCGA CRC samples (CMS1, n=8; CMS2, n=26; CMS3, n=10; CMS4, n=19) and **(B)** scRNA-seq of Lee et al. CRC samples (CMS1, n=5; CMS2, n=8; CMS3, n=4, CMS4, n=6). The mean expression is depicted through heat intensity.

**Figure S21: Spatial heterogeneity of CRCs.** Low magnification views of **(A)** MUC5AC staining of a MSS CRC, CDX2 staining of **(B)** MSS and **(C)** MSI-H CRCs, and **(D)** SOX9 staining of a MSI-H CRC. Nuclei are blue in A-C, green in D. Scale bar: 1mm. **(E)** MDK staining of a MSI-H-CRC. Nuclei are purple. Scale bar: 100μm.

**Figure S22: Signaling from tumor infiltrating lymphocytes.** Heatmap representation of signaling receptor gene expression of **(A)** T cells and **(B)** plasma cells from different tumor subtypes. **(C)** Low magnification (Scale bar: 1mm) view of a tubulovillous AD and **(D)** high magnification (Scale bar: 100μm) views of different samples of tubular ADs depicting T cell distribution. **(E)** Low magnification (Scale bar: 1mm) view of a microvesicular HP and **(F)** high magnification (Scale bar: 100μm) views of different samples of SSLs depicting T cell distribution.

**Figure S23: T cell distribution in CRCs. (A)** Low magnification (Scale bar: 1mm) view of a MSS CRC, and **(B)** high magnification (Scale bar: 100μm) view of 2 different areas of the cancer depicting T cell distribution. **(C,D)** Same as A,B but with another MSS specimen. **(E)** Low magnification (Scale bar: 1mm) view of a MSI-H CRC, and **(F)** high magnification (Scale bar: 100μm) view of a MUC5AC-low area and a MUC5AC-high area of the cancer depicting T cell distribution.

**Table S1: Characteristics of Tennessee Colorectal Polyp Study participants.**

**Table S2: Characteristics of COLON MAP study participants and polyp samples.**

**Table S3: Pairwise CytoTRACE Mann-Whitney U test results for ASC, SSC, and NL cell populations.**

**Table S4: Count-based GSEA of Cluster 0 in ASC heterogeneity analysis.**

**Table S5: Count-based GSEA of Cluster 1 in ASC heterogeneity analysis.**

**Table S6: Count-based GSEA of Cluster 2 in ASC heterogeneity analysis.**

**Table S7: Count-based GSEA of ASC.**

**Table S8: Count-based GSEA of SSC.**

**Table S9: Regulon-based GSEA of ASC.**

**Table S10: Regulon-based GSEA of SSC.**

**Table S11: Count-based GSEA of Cluster 0 in SSC heterogeneity analysis.**

**Table S12: Count-based GSEA of Cluster 1 in SSC heterogeneity analysis.**

**Table S13: Count-based GSEA of Cluster 2 in SSC heterogeneity analysis.**

**Table S14: Count-based GSEA of Cluster 3 in SSC heterogeneity analysis.**

**Table S15: Characteristics of Cooperative Human Tumor Network (CHTN) participants and samples.**

**Table S16: Pairwise CytoTRACE Mann-Whitney U test results for ASC, SSC, MSI-H, and MSS cell populations.**

**Table S17: Regulon-based GSEA of MSI-H cancer.**

**Table S18: Regulon-based GSEA of MSS cancer.**

**Table S19: Pairwise CMS1 score T-test results for ASC, SSC, MSI-H, and MSS cell populations.**

**Table S20: Pairwise CMS2 score T-test results for ASC, SSC, MSI-H, and MSS cell populations.**

**Table S21: Pairwise CMS3 score T-test results for ASC, SSC, MSI-H, and MSS cell populations.**

**Table S22: Pairwise CMS4 score T-test results for ASC, SSC, MSI-H, and MSS cell populations.**

## STAR METHODS

### RESOURCE AVAILABILITY

#### Lead Contact

Further information and requests for resources and reagents should be directed to and will be fulfilled by Lead Contact: Ken Lau, PhD at.

#### Materials Availability

This study did not generate any unique reagents.

#### Data and Code Availability

The raw scRNA-seq sequencing and final QC filtered data generated for the COLON MAP cohort from this study are available at through Synapse in Coordination with HTAN DCC and is available with the Synapse DOI: 10.7303/syn21050481. The code used to process these scRNA-seq data are available at: https://github.com/KenLauLab/STAR_Protocol. Additionally, the full GSEA tables and their respective statistics generated through g:Profiler are available in **Supplemental Data 1.**

### EXPERIMENTAL MODEL AND SUBJECTS DETAILS

#### Colorectal Molecular Atlas Project (COLON MAP)

COLON MAP participants were recruited from among adults undergoing routine screening or surveillance colonoscopy at Vanderbilt University Medical Center in Nashville, TN, USA that began in March 2019 and is still on-going, participant characteristics are shown in Table 1. The participants included in this study are the first 30 participants from COLON MAP with polyps collected for analysis by scRNA-seq. All participants provided written informed consent approved by the Vanderbilt University Medical Center Institutional Review Board.

#### Cooperative Human Tumor Network (CHTN)

Tissue was collected for COLON MAP from three colorectal cancer (CRC) patients via the CHTN Western Division. The samples and participants consist of: a stage IIIB MSI cancer from a 69 year old white male, a stage IIIB MSS cancer from a 54 year old white female, and a stage I MSS cancer from a 40 year old black male. Participant characteristics of the three CRC patients obtained from the CHTN Western Division are shown in Table 14.

A colorectal carcinoma progression tissue microarray (TMA) was provided by the CHTN MidAtlantic Division which included cores from 54 individuals. The mean (standard deviation) age of the individuals included on the TMA was 56.9 (14.7), 56.9% were men, and 43.1% were women. Race and ethnicity were not provided. Information on the TMA is available at https://chtn.sites.virginia.edu/chtn-crc2

#### Tennessee Colorectal Polyp Study (TCPS)

The TCPS was a large colonoscopy-based case-control study among individuals undergoing colonoscopy in Nashville, Tennessee, USA between February 2003 and October 2010. Institutional approval for human subjects research was provided by the VUMC and VA Institutional Review Boards and the VA Research and Development Committee.

Detailed methods have been previously published (Davenport et al., 2018). In this analysis, a subset of TCPS formalin-fixed paraffin-embedded polyps which were previously analyzed by bulk RNA-seq were included to validate findings from the COLON MAP scRNA-seq analysis. In addition, a subset of fresh frozen polyps which were selected for targeted gene sequencing were also included. Characteristics of the participants included in this analysis are provided in Table 2.

### METHOD DETAILS

#### COLON MAP Eligibility and Recruitment

Eligibility criteria for COLON MAP include ability to provide informed consent, free-living (not a resident of an institution), ability to speak and understand English, aged 40 to 75 years, permanent residence or telephone, and no personal confirmed or suspected histories of hereditary polyposis syndromes, familial or genetic colorectal cancer syndromes, inflammatory bowel disease, primary sclerosing cholangitis, colon resection or colectomy, cancer, neoadjuvant therapy, or cystic fibrosis. Eligible individuals were first identified from the schedule within the electronic health record (EHR) and assigned a random number. Potential participants were further selected using a stratified weighted random sampling design to increase the inclusion of non-White or Latinx participants in the study. Within strata of colonoscopy appointment day and time, random sampling was weighted by EHR-derived racial/ethnic category (White non-Latinx vs all other races and ethnicities) such that non-White or Latinx patients were first selected at random within colonoscopy day and time. White non-Latinx patients were then selected at random within remaining time slots.

Following selection, study staff conducted a manual review of the EHR to confirm study eligibility. The majority of eligible individuals were mailed a letter to introduce the study and a few days later were attempted to be reached by telephone to discuss their willingness to participate in the study. Individuals who were willing to participate completed an additional screening form to confirm eligibility and eligible and willing individuals completed an interviewer-administered computer-assisted telephone interview to solicit information on personal health history, family history of cancer and polyps, lifestyle factors, and other risk factors for colorectal polyps and cancer. When the schedule of the study staff would allow, individuals who were not reached by telephone were approached in the waiting room to determine eligibility and willingness as well as some individuals who did not receive a mailing.

#### COLON MAP Biological Specimen Collection and Processing, blood and oral rinse

Prior to the procedure, an oral mouthwash rinse sample was collected from participants. Blood was also collected through the IV line prior to colonoscopy in EDTA and serum tubes. The EDTA and serum samples were spun at 1,500g for 10 minutes, using a refrigerated centrifuge (at 4°C). The plasma was pipetted into four sterile 2ml cryovials, white blood cells were aliquoted into two 2ml vials, and red blood cells were stored in two 2ml vials after being washed two times with cold saline solution. Serum was pipetted into four 2ml vials and the blood clot into two 2ml vial. The mouth rinse samples were centrifuged, and the pellets were suspended using TE buffer, then aliquoted into a 2ml vial. All samples were placed into −80°C freezers for storage until use.

#### COLON MAP Biological Specimen Collection and Processing, colorectal tissue

During the colonoscopy, the gastroenterologist used biopsy forceps to collect normal appearing mucosa samples from the ascending and descending colon for all participants. One of the biopsies from each colon segment was placed into RPMI. Any polyps were removed during the colonoscopy per standard clinical practice. In this analysis, the first polyp which was removed from a participant that was larger than 0.5 cm was selected for scRNA-seq analysis (index lesion). Polyps which were removed intact were bisected along the vertical access using a sterile razor blade and half was placed in RPMI. For polyps which were removed piecemeal, the second largest piece was placed in RPMI. The other portions of the polyps were placed into formalin for diagnosis and fixed and processed using standard clinical practice in the Vanderbilt Pathology Laboratory. All polyps which were placed in RPMI were immediately transported to the research lab for use in scRNA-seq analysis.

#### COLON MAP Histopathology and Diagnosis

Information on the colonoscopy and diagnosis was initially abstracted from the EHR colonoscopy and pathology reports by study staff including *in vivo* size and polyp location. Two study pathologists additionally reviewed each case to standardize diagnoses and identify HP subtypes which are not part of routine clinical practice. For polyps which were partial due to the sampling for this study, the portion which had been reserved for clinical diagnosis was reviewed. SSLs were defined using the World Health Organization criteria of at least one distorted, dilated, or horizontally branched crypt within the polyp (Rex et al., 2012). Subtypes of ADs were identified using standard diagnostic criteria based on the villous component (tubular (< 25% villous component), tubulovillous (25%-74% villous component), and villous (≥ 75%)). HPs were classified as microvesicular HP or goblet cell HP (Leggett and Whitehall, 2010). In this analysis, participants were classified based upon the diagnosis of their index lesion but may have had synchronous polyps with the same or different histopathologies as shown in Table 1.

#### COLON MAP DNA/RNA extraction

DNA was isolated from thawed buffy coat or mouth rinse samples using a QIAmp DNA kit (Qiagen). DNA and total RNA were isolated from FFPE sections using AllPrep DNA/RNA FFPE kit (Qiagen, Cat# 80234) following manufactory’s instructions. In brief, tumor tissues were scraped from 1-5 of 10 μm FFPE sections, deparaffinized using xylene, and lysed in an optimized lysis buffer that contains proteinase K. The sample lysate was then centrifuged to give an RNA-containing supernatant and a DNA-containing pellet, which then undergo separate purification procedures. Following the proteinase K digestion to release RNA or DNA from the tissue, a higher temperature incubation to reverse formalin crosslinking, and the DNase or RNase treatment to eliminate genomic DNA or RNA contamination, the RNA and DNA samples were then purified by the RNeasy and QIAamp MinElute spin column, respectively, and eluted for downstream applications.

#### COLON MAP Single cell encapsulation and library generation for scRNA-seq

Colonic biopsy samples were first placed into RPMI solution, minced to approximately 4mm^2^, and washed with 1x DPBS. These samples were then incubated in chelation buffer (4mM EDTA, 0.5 mM DTT) at 4 °C for 1 h 15 min. Then, the resulting suspension was dissociated with cold protease and DNAse I for 25 minutes. This suspension was titurated throughout the process, every 10 minutes, then washed three times with 1x DPBS before encapsulation. The encapsulation and library preparation procedure was based on the inDrop platform. Accordingly, the protocol used is as published in Southard-Smith et al. (Southard-Smith et al., 2020), which is closely based on the 1Cell-Bio library preparation protocol version 2.3. Following sequencing library preparation, they were sequenced in a S4 flow cell using a PE150 kit on an Illumina NovaSeq 6000 to a target of 150 million reads.

#### COLON MAP scRNA-seq Read Alignment and Droplet Matrix Generation

We demultiplexed, aligned, and corrected the detected read counts of these libraries with the DropEst pipeline (Petukhov et al., 2018), using the STAR aligner with the Ensembl reference genome (Dobin et al., 2013), GRCh38 release 25. This was paired with the corresponding GTF annotations. Our use of the inDrops scRNA-seq platform yielded sequencing libraries (Klein et al., 2015; Zilionis et al., 2017), for each individual sample, following a super-poissonian bead distribution.

#### COLON MAP scRNA-seq Droplet Matrix Quality Control

We identified high-quality, cell-containing droplets and their respective barcodes through the joint application of cumulative sum inflection point thresholding, our dropkick QC algorithm (Heiser et al., 2020), and prior-knowledge gene expression profiling. This droplet matrix was processed as an AnnData object using our preprocessing pipeline which utilizes the Scanpy toolkit (Wolf et al., 2018). First, we ran dropkick with 5-fold cross validation on the unprocessed droplet matrix, which assigned each barcode a probability of being a high-quality cell. Second, the droplet matrix was preprocessed for low dimensional analysis through finding the inflection point of the cumulative sum curve, and droplets with low information content were removed. Third, the remaining cells were normalized to the median number of counts per single-cell library per dataset, inverse hyperbolic sine transformed, and then scaled as a z-score. Fourth, normalized matrices were projected into 2 dimensions by using its 50 principal component decomposition to initialize a UMAP (McInnes et al., 2018). Fifth, gene expression and dropkick probability scores were overlaid and checked for consistency. The genes overlaid were based on prior knowledge of the colonic epithelial markers, deferring to dropkick scores when no markers were found. Sixth, the selection of the final set of high-quality cell-containing droplets were determined by setting a binarization threshold on the dropkick probability scores, given concordance to marker gene expression and other general quality metrics such as total counts, mitochondrial count percentage, and transcriptional diversity.

#### CHTN Eligibility and Recruitment

See the **EXPERIMENTAL MODEL AND SUBJECTS DETAILS** subsection for **COLON MAP Eligibility and Recruitment.**

#### COLON MAP and CHTN TMA MxIHC

MxIHC was performed by iterative antibody staining and chromogen removal based on the protocol in Tsujikawa et al. (Tsujikawa et al., 2017). Chromogen was removed between sequential rounds through sequential alcohol baths, and antibody was stripped by high temperature (95°C for 15 minutes). Incubation and detection conditions are listed in the table below.

| **Marker** | **Primary Antibody Concentration** | **Primary Antibody Incubation Conditions** | **Detection Conditions** | **Chromogen Conditions** |
| --- | --- | --- | --- | --- |
| Phospho-p44/42 MAPK (Erk1/2) | 1:800 | Room Temperature Overnight | 30 minutes | 12 minutes |
| MUC5AC | 1:400 | Room Temperature Overnight | 30 minutes | 12 minutes |
| CDX2 | 1:800 | Room Temperature Overnight | 30 minutes | 12 minutes |
| Midkine (MDK) | 1:75 | Room Temperature Overnight | 30 minutes | 12 minutes |
| YAP | 1:400 | Room Temperature Overnight | 30 minutes | 12 minutes |
| MLH1 | 1:100 | Room Temperature 60 minutes | 30 minutes | 12 minutes |

#### COLON MAP and CHTN TMA MxIF

Cyclical antibody staining, detection, and dye inactivation was performed as described previously by Gerdes et al. (Gerdes et al., 2013) Briefly, fluorescence imaging was performed on a GE IN Cell Analyzer 2500 using the Cell DIVe platform. Images were acquired at x200 magnification with exposure times determined for each antibody. Antibody reagents are listed in the **Key Resources Table**. Staining sequence, conditions, and exposure times are listed in the table below. For each round of staining, DAPI images were aligned using rigid transformations to the first imaging round. The registered images were corrected for uneven illumination and autofluorescence was removed for each channel.

| Round | Channel | Marker | Concentration | Staining Conditions | Image Exposure (ms) |
| --- | --- | --- | --- | --- | --- |
| 1 | GFP | Muc2 | 1:150 | 4° C Overnight | 50 |
| 1 | Cy3 | Collagen Peptide | 1:300 | 4° C Overnight | 10 |
| 1 | Cy5 | SNA | 1:300 | 4° C Overnight | 100 |
| 2 | GFP | Background | NA | NA | 100 |
| 2 | Cy3 | Background | NA | NA | 200 |
| 2 | Cy5 | Background | NA | NA | 500 |
| 3 | GFP | CD11B | 1:100 | 4° C Overnight | 100 |
| 3 | Cy3 | CD45 | 1:100 | 4° C Overnight | 100 |
| 3 | Cy5 | CD20 | 1:150 | 4° C Overnight | 50 |
| 4 | GFP | Background | NA | NA | 100 |
| 4 | Cy3 | Background | NA | NA | 200 |
| 4 | Cy5 | Background | NA | NA | 500 |
| 5 | GFP | PCNA | 1:150 | 4° C Overnight | 50 |
| 5 | Cy3 | B-catenin | 1:200 | 4° C Overnight | 50 |
| 5 | Cy5 | p-STAT3 | 1:75 | 4° C Overnight | 300 |
| 6 | GFP | Background | NA | NA | 100 |
| 6 | Cy3 | Background | NA | NA | 200 |
| 6 | Cy5 | Background | NA | NA | 500 |
| 7 | GFP | pEGFR | 1:200 | 4° C Overnight | 100 |
| 7 | Cy3 | CgA | 1:75 | 4° C Overnight | 50 |
| 7 | Cy5 | CD4 | 1:200 | 4° C Overnight | 200 |
| 8 | GFP | Background | NA | NA | 100 |
| 8 | Cy3 | Background | NA | NA | 200 |
| 8 | Cy5 | Background | NA | NA | 500 |
| 9 | GFP | Cox2 | 1:100 | 4° C Overnight | 100 |
| 9 | Cy3 | CD3d | 1:100 | 4° C Overnight | 100 |
| 9 | Cy5 | HLA-A | 1:200 | 4° C Overnight | 50 |
| 10 | GFP | Background | NA | NA | 100 |
| 10 | Cy3 | Background | NA | NA | 200 |
| 10 | Cy5 | Background | NA | NA | 500 |
| 11 | GFP | PanCK | 1:200 | 4° C Overnight | 50 |
| 11 | Cy3 | OLFM4 | 1:300 | 4° C Overnight | 20 |
| 11 | Cy5 | CD8 | 1:200 | 4° C Overnight | 150 |
| 12 | GFP | Background | NA | NA | 100 |
| 12 | Cy3 | Background | NA | NA | 200 |
| 12 | Cy5 | Background | NA | NA | 500 |
| 13 | GFP | Alpha-actinin | 1:100 | 4° C Overnight | 100 |
| 13 | Cy3 | CD68 | 1:200 | 4° C Overnight | 50 |
| 13 | Cy5 | NaKATPase | 1:200 | 4° C Overnight | 100 |
| 14 | GFP | Background | NA | NA | 100 |
| 14 | Cy3 | Background | NA | NA | 200 |
| 14 | Cy5 | Background | NA | NA | 500 |
| 15 | GFP | Vimentin | 1:300 | 4° C Overnight | 20 |
| 15 | Cy3 | Sox9 | 1:200 | 4° C Overnight | 50 |
| 15 | Cy5 | FOXP3 | 1:100 | 4° C Overnight | 400 |
| 16 | GFP | Background | NA | NA | 100 |
| 16 | Cy3 | Background | NA | NA | 200 |
| 16 | Cy5 | Background | NA | NA | 500 |
| 17 | GFP | Lysozyme | 1:150 | 4° C Overnight | 100 |
| 17 | Cy3 | SMA | 1:500 | 4° C Overnight | 3 |
| 17 | Cy5 | ERBB2 | 1:150 | 4° C Overnight | 500 |

#### TCPS Eligibility and Recruitment

TCPS participants were aged between 40 to 75 years of age and had no personal history of colon resection, cancer, polyposis syndrome, inflammatory bowel disease, hereditary colorectal cancer syndromes, or previous adenoma. In TCPS, the diagnostic criteria for polyps were identical to the criteria used for COLON MAP. Additionally, all polyps were reviewed by one of the COLON MAP pathologists.

#### TCPS Eligibility and Recruitment

Fresh frozen polyps were collected using the same protocol as described for COLON MAP and polyps were stored at −80°C until use. For the original projects from which the data for this analysis were obtained, polyp samples were selected among the TCPS participants with a history of multiple (>1) adenoma and advanced adenoma (≥ 1 cm, tubulovillous, villous, and/or high-grade dysplasia) who had completed a surveillance colonoscopy subsequent to sample collection. For the bulk RNA-seq analysis, 66 fresh frozen polyp samples were included. For the targeted sequencing, 274 polyps from 197 participants were included. Features of the samples included in this study are shown in **Supplemental Table 2.**

#### TCPS DNA/RNA extraction

DNA was extracted from FFPE tissue sections using QIAamp DNA FFPE Tissue Kit (Qiagen, Cat# 56404), following the manufacturer’s instructions. Briefly, tumor tissues were scraped from 1-5 of 10 μm FFPE sections, deparaffinized using xylene, and lysed under denaturing conditions with proteinase K. The sample lysate was incubated at 90°C to reverses formalin crosslinking and then applied to a QIAamp MinElute spin column, where DNA was captured on a silica membrane. The genomic DNA was then washed and eluted from the membrane.

DNA and total RNA were extracted from fresh frozen polyps and purified using Qiagen’s AllPrep DNA/RNA/miRNA Universal Kit (Qiagen, Cat# 80224), following the manufacturer’s instructions. Briefly, the frozen tissue samples were first disrupted and homogenized using Lysing Matrix E (MP Bio, Cat# 116914100) by shaking the tubes on a bead-beater at 5.5 m/sec for 30 second. The lysate was then passed through an AllPrep DNA Mini spin column. This column allows selective and efficient binding of genomic DNA. Following on-column Proteinase K digestion, the column was then washed and pure, ready-to-use DNA was eluted. Flow-through from the DNA Mini spin column was then digested by Proteinase K in the presence of ethanol and applied to the RNeasy Mini spin column, where the total RNA binds to the membrane. Following DNase I digestion, contaminants were efficiently washed away and high-quality RNA was eluted in RNase-free water. The quantity and quality of the DNA/RNA samples were checked by Nanodrop (E260/E280 and E260/E230 ratio) and by separation on an Agilent BioAnalyzer.

Total RNA, including microRNA, were extracted and purified from the fresh frozen TCPS polyps using Qiagen’s AllPrep DNA/RNA/miRNA Universal Kit (Qiagen, Cat# 80224), following the manufacturer’s instructions. Briefly, the frozen tissue samples were first disrupted and homogenized using Lysing Matrix E (MP Bio, Cat# 116914100) by shaking the tubes on a beadbeater at 5.5 m/sec for 30 second. The lysate were then digested by Proteinase K in the presence of ethanol and applied to the RNeasy Mini spin column, where the total RNA binds to the membrane. Following the on-column DNase digestion, contaminants were efficiently washed away and high-quality RNA was eluted in RNase-free water. The quantity and quality of the RNA samples were checked by Nanodrop (E260/E280 and E260/E230 ratio) and by separation on an Agilent BioAnalyzer.

#### TCPS Targeted DNA sequencing

The list of candidate genes included in the targeted sequencing was developed from a literature review of candidate mutations which showed 1) evidence that mutation is common in adenoma (>5% of adenomas), 2) evidence that the mutation is associated with or predictive of adenoma recurrence in previous studies, 3) evidence that mutation is associated with clinically more significant adenoma (i.e. advanced adenoma or multiplicity), 4) evidence that mutation is associated with colorectal field carcinogenesis, and 5) evidence that mutation is associated with colorectal cancer aggressiveness and survival. In addition, additional candidate mutations were identified from potential mutations observed in Lrig1-Cre:Apc adenomas. All primer development and next-generation sequencing were conducted by Covance. Sequencing depth was 500X. Targeted sequencing reads were aligned to the human reference genome hg19 using BWA (Li and Durbin, 2009),and then were sorted and indexed by Sambamba (Tarasov et al., 2015). Alignments were further refined, and variants were called using GATK Best Practices tools (Van der Auwera et al., 2013), including mark duplicates with Picard (Broad Institute, 2019), base quality-score recalibration, and variant calling with HaplotypeCaller and GenotypeGVCFs (Poplin et al., 2017). SNPs were filtered using GATK VariantFiltration function with the parameters “QD < 2.0 || Qual < 30.0 || FS > 60.0 || SOR>3.0 || MQ < 40.0 || MQRankSum < −12.5 || ReadPosRankSum < −8.0”, while indels were filtered with the parameters “QD < 2.0 || Qual < 30.0 || FS > 200.0 || ReadPosRankSum < −20.0”. The variants with a minor allele frequency >0.1% in ExAC, gnomAD, TOPMed or 1,000 Genomes were also removed. The functional effects of variants were annotated by ANNOVAR (Wang et al., 2010; Yang and Wang, 2015).

#### TCPS Bulk RNA Sequencing

Bulk RNA-sequencing was performed by Aros Applied Biotechnology A/S. This process involves the initial QC on an Agilent Bioanalyzer, with a minimum quality threshold of the DV200 at 30%. Total RNA-seq libraries which pass this QC threshold are prepared alongside a high-quality human reference RNA control. 100ng of RNA per sample is input to an Illumina TruSeq RNA Access Library Prep Kit, with protocol version 0.2. The yielded libraries undergo another round of QC through qPCR and quantified with a Qubit 2.0 Fluorometer, using its corresponding DNA BR Assay kit, and size profiled on an Agilent Bioanalyzer. Pools of 4 libraries in equimolar amounts are created and undergo a final round of QC. These pools are loaded onto paired-end flow cells of a HiSeq2500 equipped with a cBot for sequencing at: 101 read cycles, 7 index cycles, and 101. The samples will be sequenced on a HiSeq2500 using 101 cycles for read 1, 7 index reads, and another 101 cycles for read 2. Following sequencing data generation, the reads are demultiplexed through Illumina’s CASAVA software, which detected an average of 120 million reads per 4 sample pool.

### QUANTIFICATION AND STATISTICAL ANALYSIS

#### Regulon Network Prediction and Activity Inference

The Single-Cell rEgulatory Network Inference and Clustering or SCENIC pipeline was used to integrate multiple cancerous, pre-cancerous, and their corresponding normal tissue datasets (Aibar et al., 2017; Van de Sande et al., 2020). These sets included 2,129 cells, 34,714 cells, and 30,374 cells for integration, respectively. The normal counterparts paired with our COLON MAP cancer samples were not used for this regulon analysis. Similarly, this pipeline was also used to integrate 4,083 corresponding non-epithelial cells. For each group of datasets for integrated analysis, we concatenated the individual target datasets with an outer join and generated a combined AnnData object. This AnnData object underwent further gene filtering, selecting only those that were expressed in at least 1% of all cells, primarily for the sake of speedup in running the module inference step of SCENIC. The resulting cumulative count matrix was input, without normalization, into the first step of SCENIC with default parameters, as suggested by the published protocol. We used a Dask client to parallelize the grnboost2 version of this step. Subsequently, cisTarget was performed using default parameters and three hg19 .feather ranking databases, comparing 10 species: tss-centered-5kb, tss-centered-10kb, and 500bp-upstream. Further, this cisTarget step produced a list of detected regulons, their driving TFs, and their corresponding weights for the prediction of individual gene expression. We generated a feature matrix using these weights to build a regulon-regulon target network, per SCENIC run. This target network was based on a k-nearest neighbors graph (with k equal to the square root of the number of total regulons) of the feature matrix, with each regulon being an observation and its target genes being the features. For each of these target networks, the Leiden community detection algorithm was run at a resolution of 2 (Traag et al., 2019). This regulon-regulon target network (along with its cluster labels and average enrichment per comparison group) was exported as a weighted adjacency matrix for visualization in Cytoscape (Shannon et al., 2003). Finally, we performed AUCell with default recommended parameters across 64 threads. The resultant regulon enrichment matrix was jointly analyzed with the count-based matrix.

#### Unsupervised Clustering and Differential Gene Signature Analysis

The labeling of single-cell subpopulations was done through the Leiden algorithm, as part of the Scanpy toolkit. We performed Leiden clustering based on the KNN derived from the distances calculated in the principal component space of z-score transformed regulon enrichment scores, as these represented cell-cell transcriptional states in a more batch-robust manner. The resolution of this clustering was based on the detection of rarer populations such as enteroendocrine cells, at 2. Since this algorithm detected discrete clusters in a continuum of cell states, we aggregated multiple discrete clusters by the observation of marker gene expression. The differential testing of gene expression was performed based on these cluster labels, both in the context of raw gene counts and regulon enrichment values. For both cases, we used Mann-Whitney U tests with Benjamini-Hochberg corrections, on the raw values, implemented through scanpy’s rank_genes_groups function, identifying the top 200 genes and top 100 regulons. Further, biological insight was gathered through Scanpy’s integration of g:profiler gene set enrichment framework (Raudvere et al., 2019). The full GSEA tables and their respective statistics generated through g:Profiler are available in **Supplemental Data 1.**

#### Predicting Differentiation Potential with CytoTRACE

CytoTRACE was performed based on the default recommended settings after concatenating the batches of interest using an outer join (Gulati et al., 2020). The resultant CytoTRACE scores were stored as an observation vector for each respective AnnData object being analyzed. Pairwise comparisons between each cell group were done through two-sided, non-parametric Mann-Whitney U tests. This test was chosen after observing non-gaussian score distributions. Given the sets of three (consisting of 7,594 ASC, 3,670 SSC, and 53,824 non-ASC/SSC) or four cell groups (consisting of 6,273 ASC, 3,287 SSC, 1,272 MSS cancer, and 857 MSI-H cancer cells) compared, a total of three or six pairwise tests were performed per set of CytoTRACE scores, thus significance levels were adjusted through the Bonferroni correction method to 0.0169 and 0.0085 respectively. The results of these statistical analyses are in **Supplemental Tables S3 and S16.**

#### Single-cell CMS Scoring

The single-cell distributions of CMS scores were calculated using the CMSclassifier R package, as described by Guinney et al., 2015 (Guinney et al., 2015). To accommodate the heterogeneity of the single-cell landscape, the single sample predictor or SSP mode of the software was used after converting gene symbols to Entrez IDs. This SSP mode calculated the median correlation distance between each single cell to established, standard centroids derived from CMS1, CMS2, CMS3, and CMS4 CRC subtypes. Further, these score distributions were visualized through a normalized kernel density estimation implemented in the Seaborn python package. In total, these calculations were performed on 6,273 ASC, 3,287 SSC, 1,272 MSS cancer, and 857 MSI-H cancer cells. Pairwise comparisons between each cell group were done through two-sided, parametric T-tests of independent samples with unequal population variances (Welch’s T-test). This test was chosen after observing gaussian score distributions. Given the four cell groups compared, a total of six pairwise tests were performed per set of CMS scores, thus significance levels were adjusted through the Bonferroni correction method to 0.0085. The results of these statistical analyses are in **Supplemental Tables S19-S22.**

#### Trajectory Inference

pCreode was used to map the developmental state transitions of the cumulative transcriptional landscape of our pre-cancer and normal COLON MAP samples, consisting of 65,088 cells (Herring et al., 2018). This algorithm was generalized to process regulon-based principal components, inheriting its batch-robust properties. By examining the variation captured by the principal components, we selected the first 4 components based on their capture of rare cell populations, such as Tuft and enteroendocrine cells. We developed this algorithm to traverse a density weighted KNN generated from the pairwise distances between each single cell; subsequently, we used a histogram thresholding method to estimate the neighborhood distance cutoff for calculating local densities. These densities were used as input to a supervised variant of pCreode, which established developmental endstates through K-means clustering and marker-defined labels. The downsampling and noise parameters were both set to 4, resulting in samples of around 6,000 cells per run, and repeated 50 times. Each of these runs was scored by the minimization of the Gromov–Hausdorff distance, resulting in a single, most representative graph layout. Overlays were generated based on pre-computed single-cell observation vectors, such as a CytoTRACE score, or the normalized, transformed, and z-scored gene expression values.

#### Subclone Phylogeny Estimation

We used a DENDRO, an algorithm designed to reconstruct subclonal phylogenies within scRNA-seq datasets (Zhou et al., 2020). Since our sequencing libraries are generated through the tag-based inDrop method, the short, 3’-biased reads necessitated the aggregation of clusters of single-cells. For each of the 8 libraries we performed this analysis on (2 TA, 2 TVA, 2 SSL, 2 HP), we defined 20 clusters through regulon neighborhood-based K-means clustering (Hartigan and Wong, 1979). Thus, we predicted the average genotypic representation of multiple single-cells and created a phylogenetic tree between the defined clusters. DropEst produced a filtered and sorted .bam file, which we derived read information from, and split into 20 distinct .bam files using the sinto python package (Tim Stuart, 2018). These bam files were then processed with GATK4 (Van der Auwera et al., 2013), according to guidelines detailed by Zhou et al. The GATK4 steps of this pipeline involved the following: adding read groups, marking duplicates, splitting N-cigar reads, applying base quality recalibration with known SNPS, haplotype calling to generate VCF files, consolidating these VCFs into genomicsdb databases, and then genotyping these data. DENDRO operated on the called genotypes to produce a dendrogram based on a beta-binomial distribution to account for single-cell transcriptional stochasticity.

#### Image Analysis

To understand the spatial distribution of cell types within epithelial and stromal tissue compartments, we performed an analysis of the distribution of T cells (CD4+ and CD8+) cells in AD and SSL samples imaged with MxIF. For each field of view, a tissue mask was created using a relaxed nuclear staining threshold, which was refined manually to demarcate tumor regions for quantification. The tumor region was divided into epithelial an epithelial region (masked by beta-Catenin, pan-Cytokeratin, and NaKATPase expression), and a stromal region (tumor mask minus the epithelial mask). CD4+ T cells were defined by intersecting CD4 and CD3 masks, while CD8+ cells were defined using the CD8 marker. We calculated the pixels occupied by CD4+ T cells in the stromal region, normalized to the number of pixels occupied by the stromal area. We calculated the pixels occupied by CD8+ T cells in the epithelial region, normalized to the epithelial nuclei pixels. A ratio of intraepithelial CD8+ cells to stromal CD4+ T cells was calculated from the two values. A t-test was performed between the AD and SSL fields of view.

