## Supplemental Figures 1-23 for "Human colorectal pre-cancer atlas identifies distinct molecular programs underlying two major subclasses of pre-malignant tumors"

### Figure S1

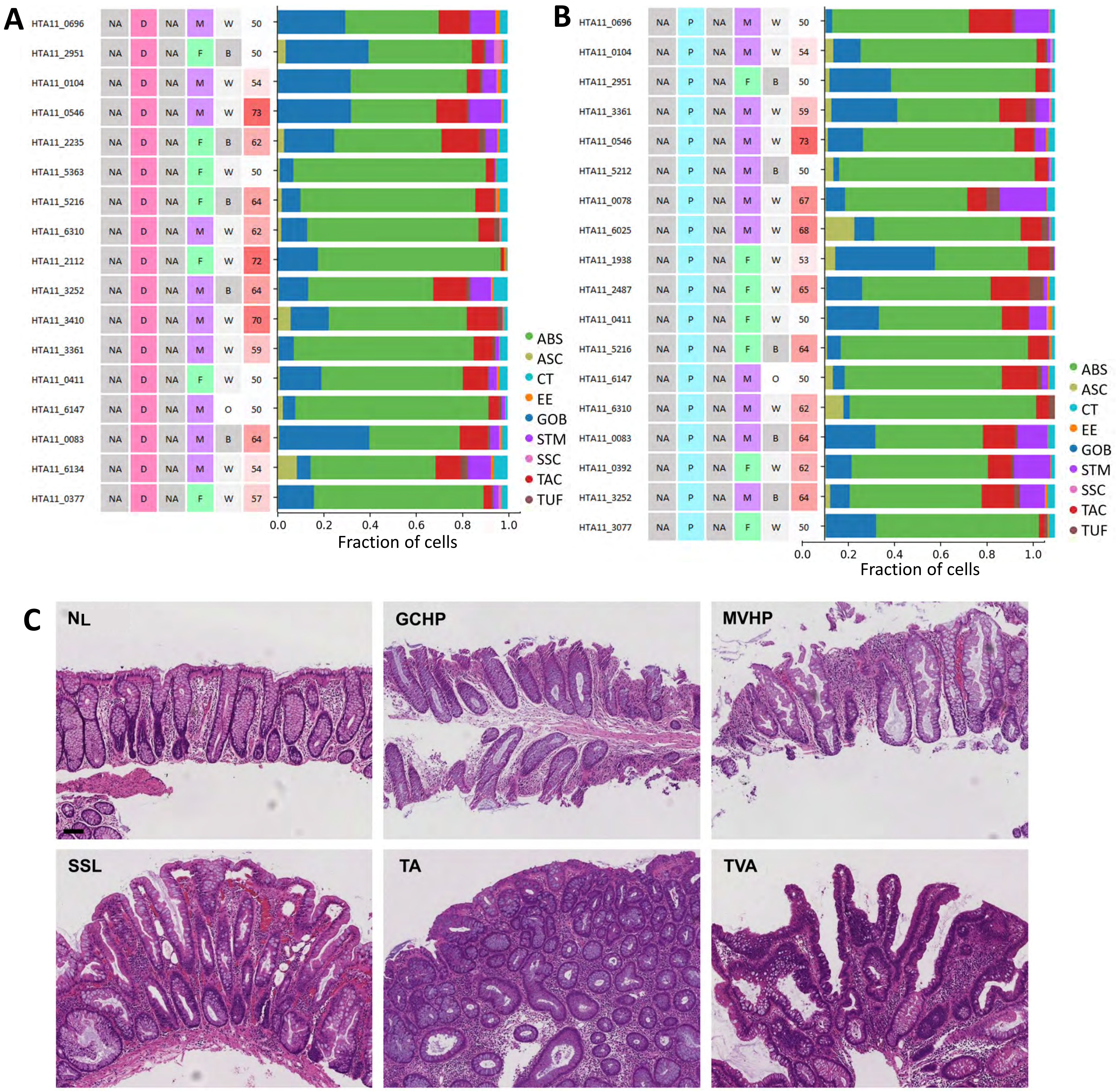
A
B
C
ABS
ASC
CT
EE
GOB
STM
SSC
TAC
TUF
Fraction of cells
0.0
0.2
0.4
0.6
0.8
1.0
NL
GCHP
MVHP
SSL
TA
TVA

### Figure S2

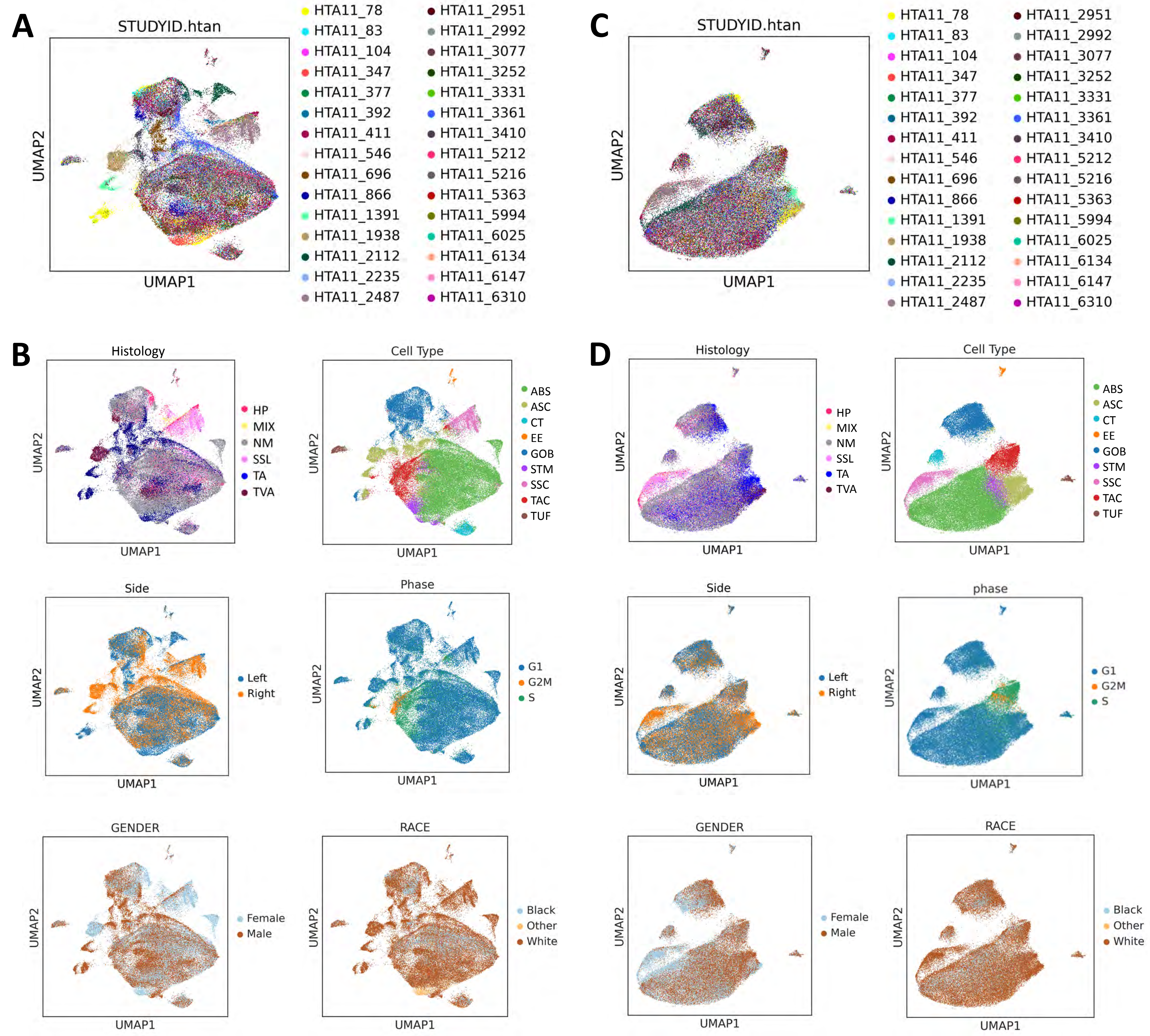
A
STUDYID.htan
UMAP2
UMAP1
HTA11_78
HTA11_83
HTA11_104
HTA11_347
HTA11_377
HTA11_392
HTA11_411
HTA11_546
HTA11_696
HTA11_866
HTA11_1391
HTA11_1938
HTA11_2112
HTA11_2235
HTA11_2487
HTA11_2951
HTA11_2992
HTA11_3077
HTA11_3252
HTA11_3331
HTA11_3361
HTA11_3410
HTA11_5212
HTA11_5216
HTA11_5363
HTA11_5994
HTA11_6025
HTA11_6134
HTA11_6147
HTA11_6310
C
STUDYID.htan
UMAP2
UMAP1
B
Histology
HP
MIX
NM
SSL
TA
TVA
Cell Type
ABS
ASC
CT
EE
GOB
STM
SSC
TAC
TUF
Side
Left
Right
Phase
G1
G2M
S
GENDER
Female
Male
RACE
Black
Other
White
D
Histology
Cell Type
Side
phase
GENDER
RACE

### Figure S3

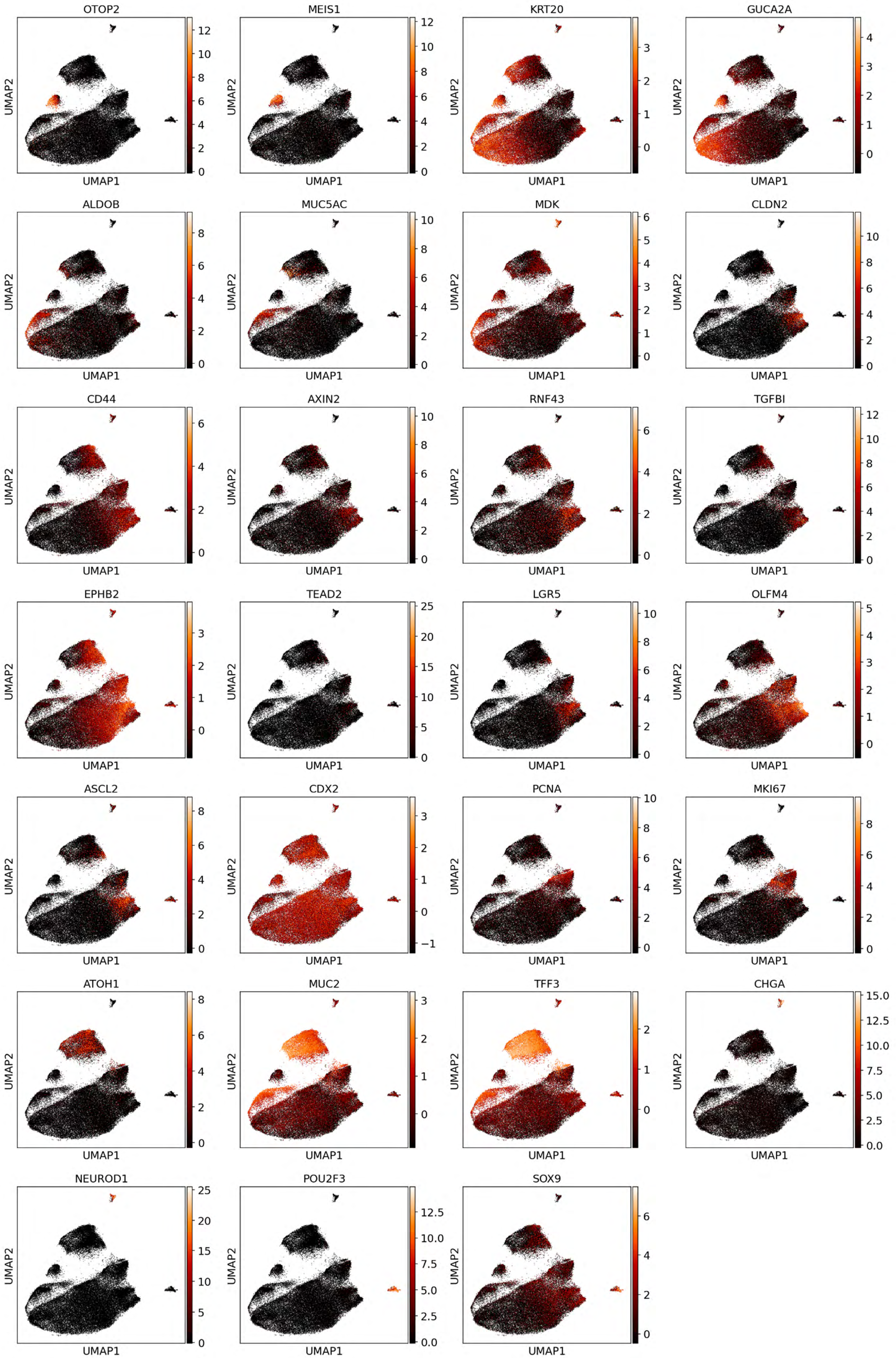
OTOP2
MEIS1
KRT20
GUCA2A
ALDOB
MUC5AC
MDK
CLDN2
CD44
AXIN2
RNF43
TGFBI
EPHB2
TEAD2
LGR5
OLFM4
ASCL2
CDX2
PCNA
MKI67
ATOH1
MUC2
TFF3
CHGA
NEUROD1
POU2F3
SOX9
UMAP1
UMAP2

Figure S4

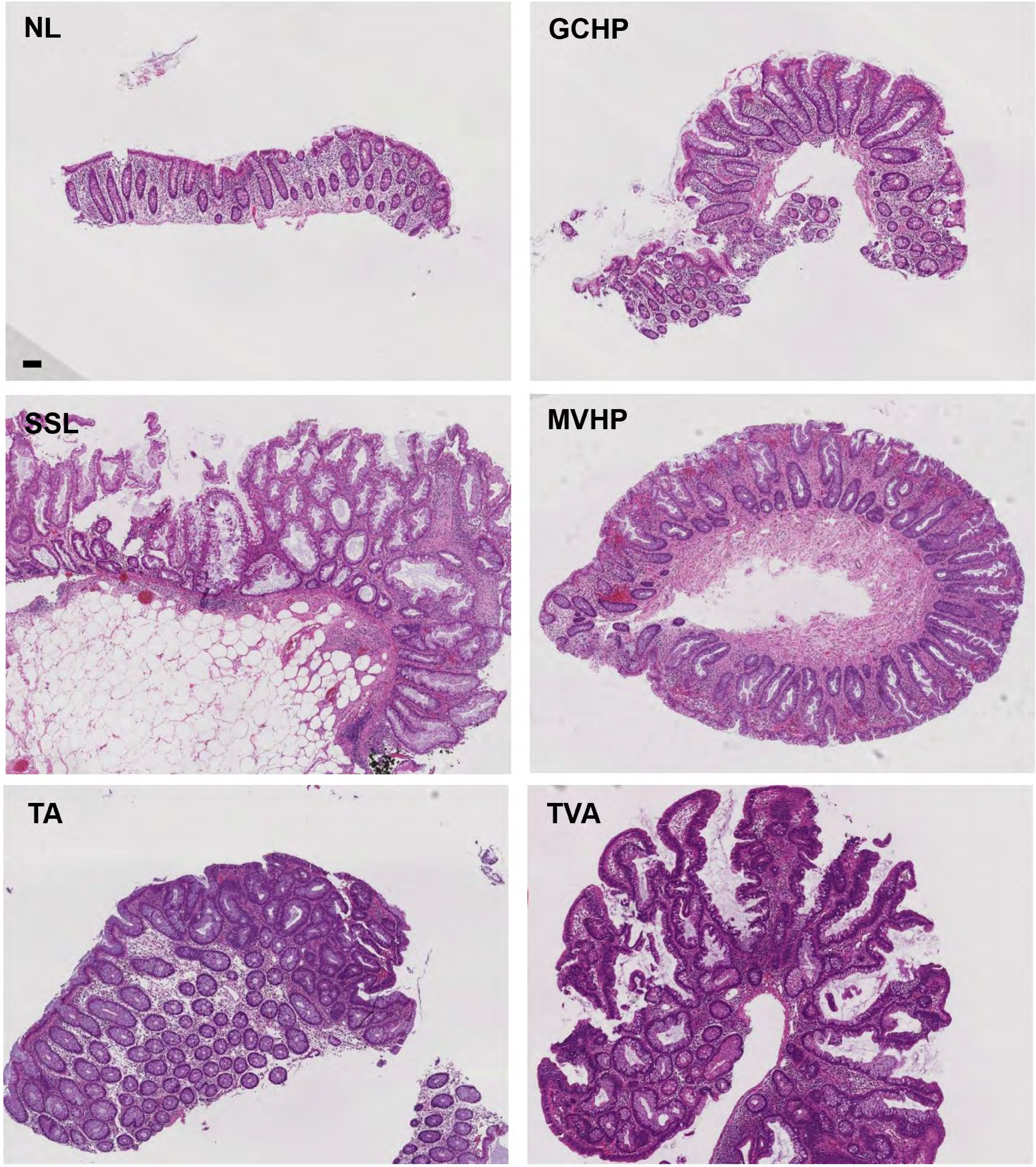

**Figure S5**

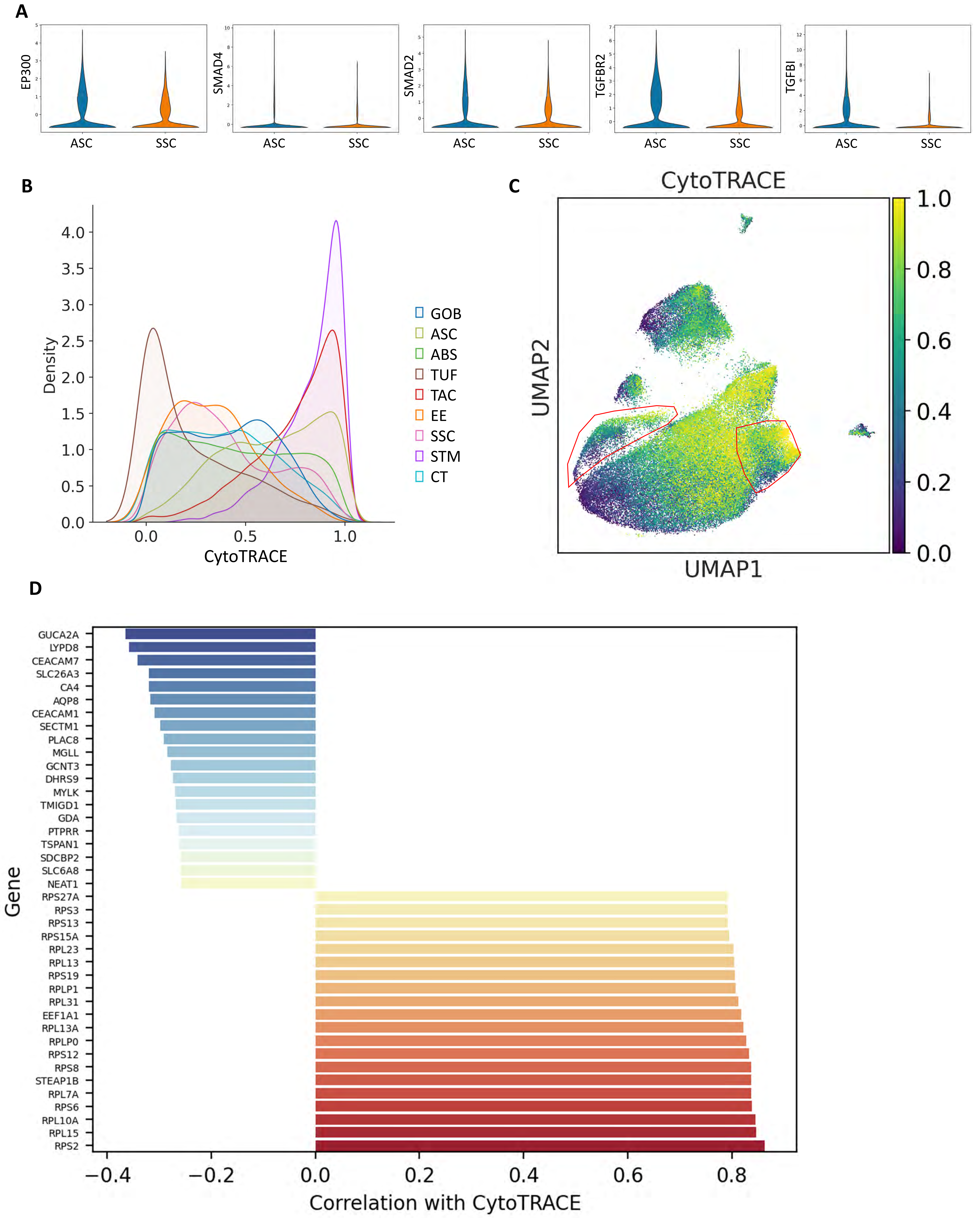
A
EP300
SMAD4
SMAD2
TGFBR2
TGFBI
ASC
SSC
B
Density
CytoTRACE
4.0
3.5
3.0
2.5
2.0
1.5
1.0
0.5
0.0
0.0
0.5
1.0
GOB
ASC
ABS
TUF
TAC
EE
SSC
STM
CT
C
CytoTRACE
UMAP2
UMAP1
1.0
0.8
0.6
0.4
0.2
0.0
D
Gene
GUCA2A
LYPD8
CEACAM7
SLC26A3
CA4
AQP8
CEACAM1
SECTM1
PLAC8
MGLL
GCNT3
DHRS9
MYLK
TMIGD1
GDA
PTPRR
TSPAN1
SDCBP2
SLC6A8
NEAT1
RPS27A
RPS3
RPS13
RPS15A
RPL23
RPL13
RPS19
RPLP1
RPL31
EEF1A1
RPL13A
RPLP0
RPS12
RPS8
STEAP1B
RPL7A
RPS6
RPL10A
RPL15
RPS2
−0.4
−0.2
0.0
0.2
0.4
0.6
0.8
Correlation with CytoTRACE

### Figure S6

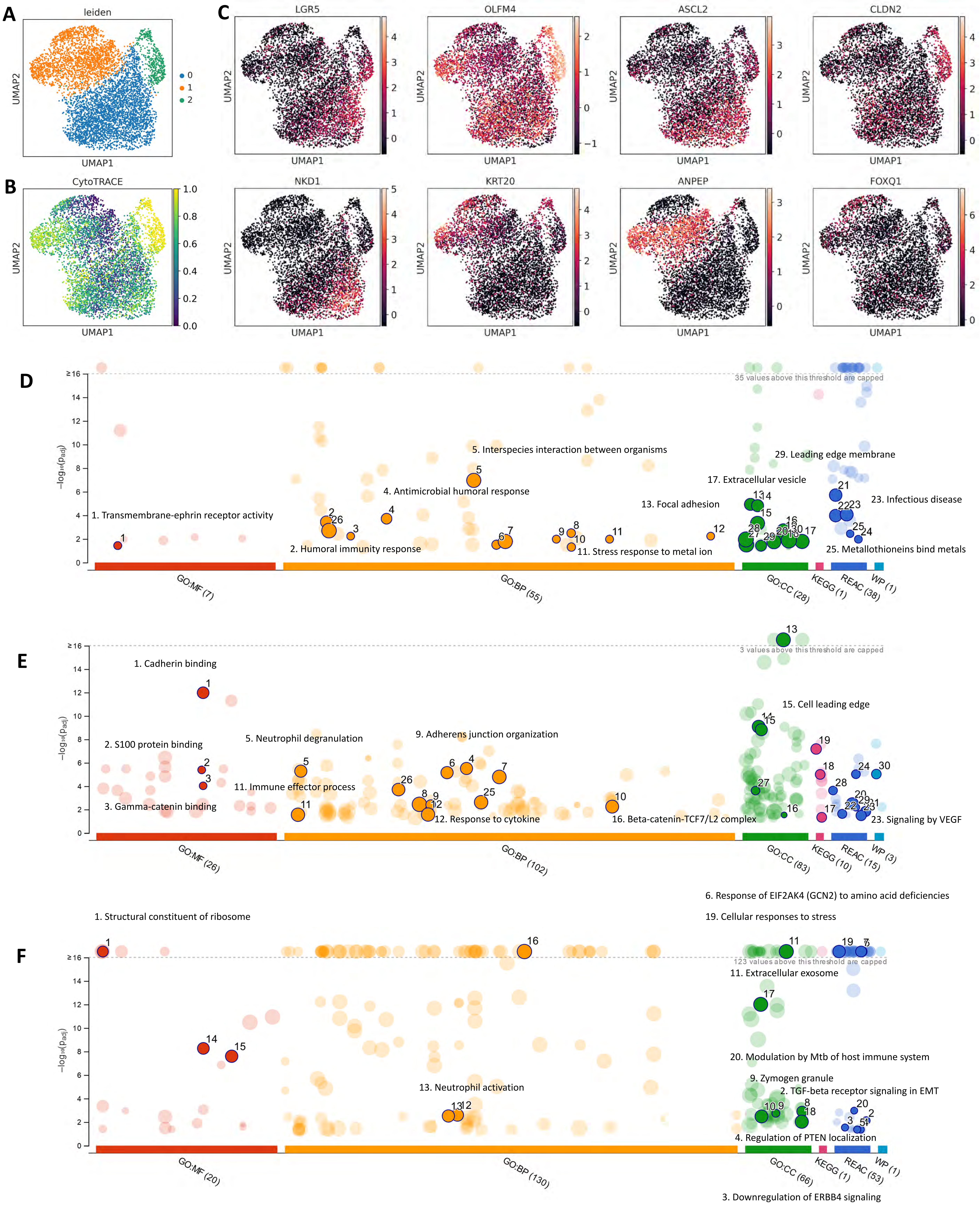

A
leiden
UMAP2
UMAP1
0
1
2
B
CytoTRACE
1.0
0.8
0.6
0.4
0.2
0.0
C
LGR5
OLFM4
ASCL2
CLDN2
NKD1
KRT20
ANPEP
FOXQ1
D
−log₁₀(p_adj)
35 values above this threshold are capped
1. Transmembrane-ephrin receptor activity
2. Humoral immunity response
4. Antimicrobial humoral response
5. Interspecies interaction between organisms
11. Stress response to metal ion
13. Focal adhesion
17. Extracellular vesicle
29. Leading edge membrane
23. Infectious disease
25. Metallothioneins bind metals
GO:MF (7)
GO:BP (55)
GO:CC (28)
KEGG (1)
REAC (38)
WP (1)
E
3 values above this threshold are capped
1. Cadherin binding
2. S100 protein binding
3. Gamma-catenin binding
5. Neutrophil degranulation
11. Immune effector process
9. Adherens junction organization
12. Response to cytokine
16. Beta-catenin-TCF7/L2 complex
15. Cell leading edge
23. Signaling by VEGF
GO:MF (26)
GO:BP (102)
GO:CC (83)
KEGG (10)
REAC (15)
WP (3)
6. Response of EIF2AK4 (GCN2) to amino acid deficiencies
1. Structural constituent of ribosome
19. Cellular responses to stress
F
123 values above this threshold are capped
11. Extracellular exosome
13. Neutrophil activation
20. Modulation by Mtb of host immune system
9. Zymogen granule
2. TGF-beta receptor signaling in EMT
4. Regulation of PTEN localization
GO:MF (20)
GO:BP (130)
GO:CC (66)
KEGG (1)
REAC (53)
WP (1)
3. Downregulation of ERBB4 signaling

### Figure S7

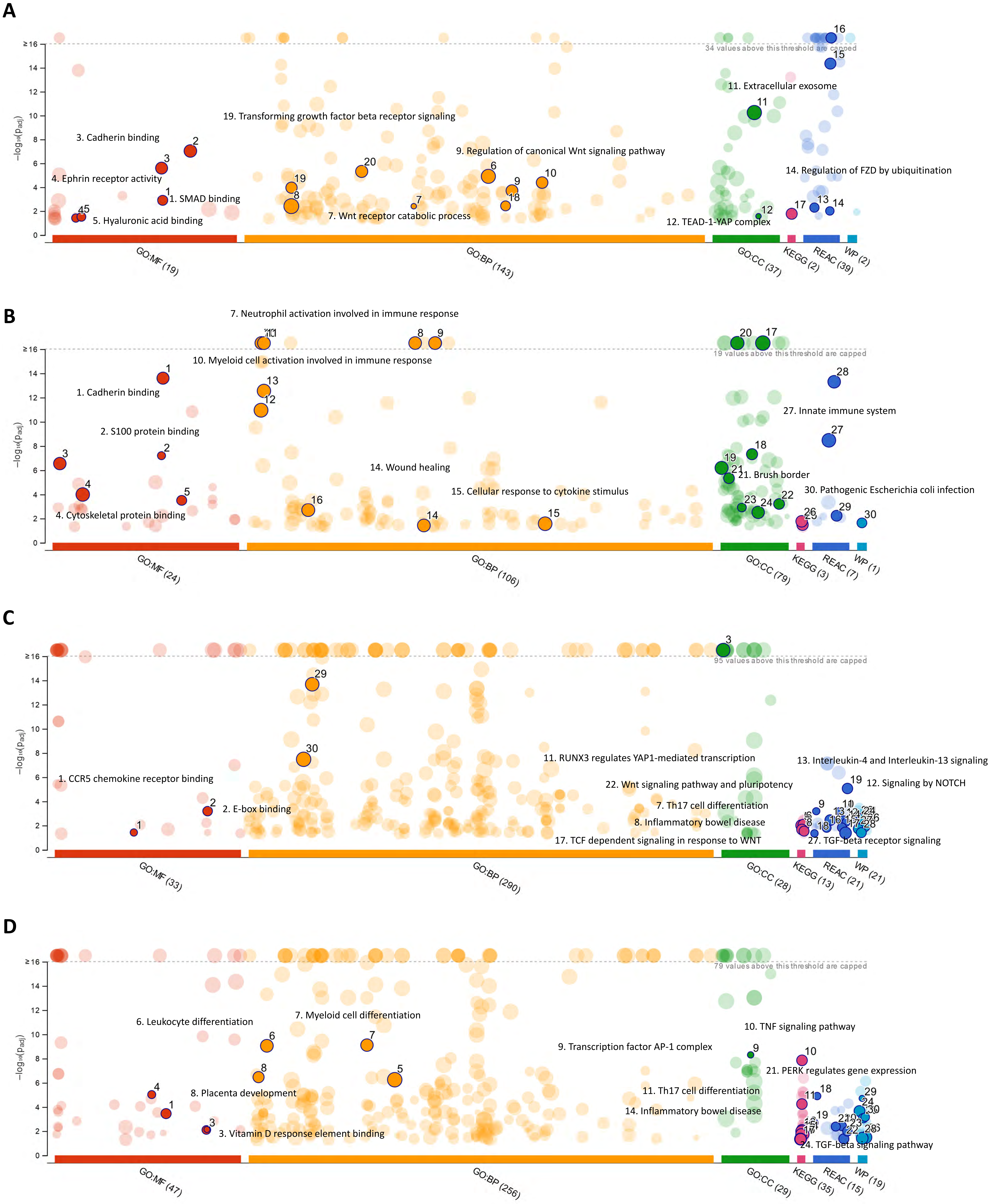
A
34 values above this threshold are capped
−log₁₀(p_adj)
3. Cadherin binding
4. Ephrin receptor activity
1. SMAD binding
5. Hyaluronic acid binding
19. Transforming growth factor beta receptor signaling
7. Wnt receptor catabolic process
9. Regulation of canonical Wnt signaling pathway
11. Extracellular exosome
14. Regulation of FZD by ubiquitination
12. TEAD-1-YAP complex
GO:MF (19)
GO:BP (143)
GO:CC (37)
KEGG (2)
REAC (39)
WP (2)
B
19 values above this threshold are capped
7. Neutrophil activation involved in immune response
10. Myeloid cell activation involved in immune response
1. Cadherin binding
2. S100 protein binding
4. Cytoskeletal protein binding
14. Wound healing
15. Cellular response to cytokine stimulus
27. Innate immune system
21. Brush border
30. Pathogenic Escherichia coli infection
GO:MF (24)
GO:BP (106)
GO:CC (79)
KEGG (3)
REAC (7)
WP (1)
C
95 values above this threshold are capped
1. CCR5 chemokine receptor binding
2. E-box binding
11. RUNX3 regulates YAP1-mediated transcription
22. Wnt signaling pathway and pluripotency
7. Th17 cell differentiation
8. Inflammatory bowel disease
17. TCF dependent signaling in response to WNT
13. Interleukin-4 and Interleukin-13 signaling
12. Signaling by NOTCH
27. TGF-beta receptor signaling
GO:MF (33)
GO:BP (290)
GO:CC (28)
KEGG (13)
REAC (21)
WP (21)
D
79 values above this threshold are capped
6. Leukocyte differentiation
7. Myeloid cell differentiation
8. Placenta development
3. Vitamin D response element binding
9. Transcription factor AP-1 complex
10. TNF signaling pathway
21. PERK regulates gene expression
11. Th17 cell differentiation
14. Inflammatory bowel disease
24. TGF-beta signaling pathway
GO:MF (47)
GO:BP (256)
GO:CC (29)
KEGG (35)
REAC (15)
WP (19)

### Figure S8

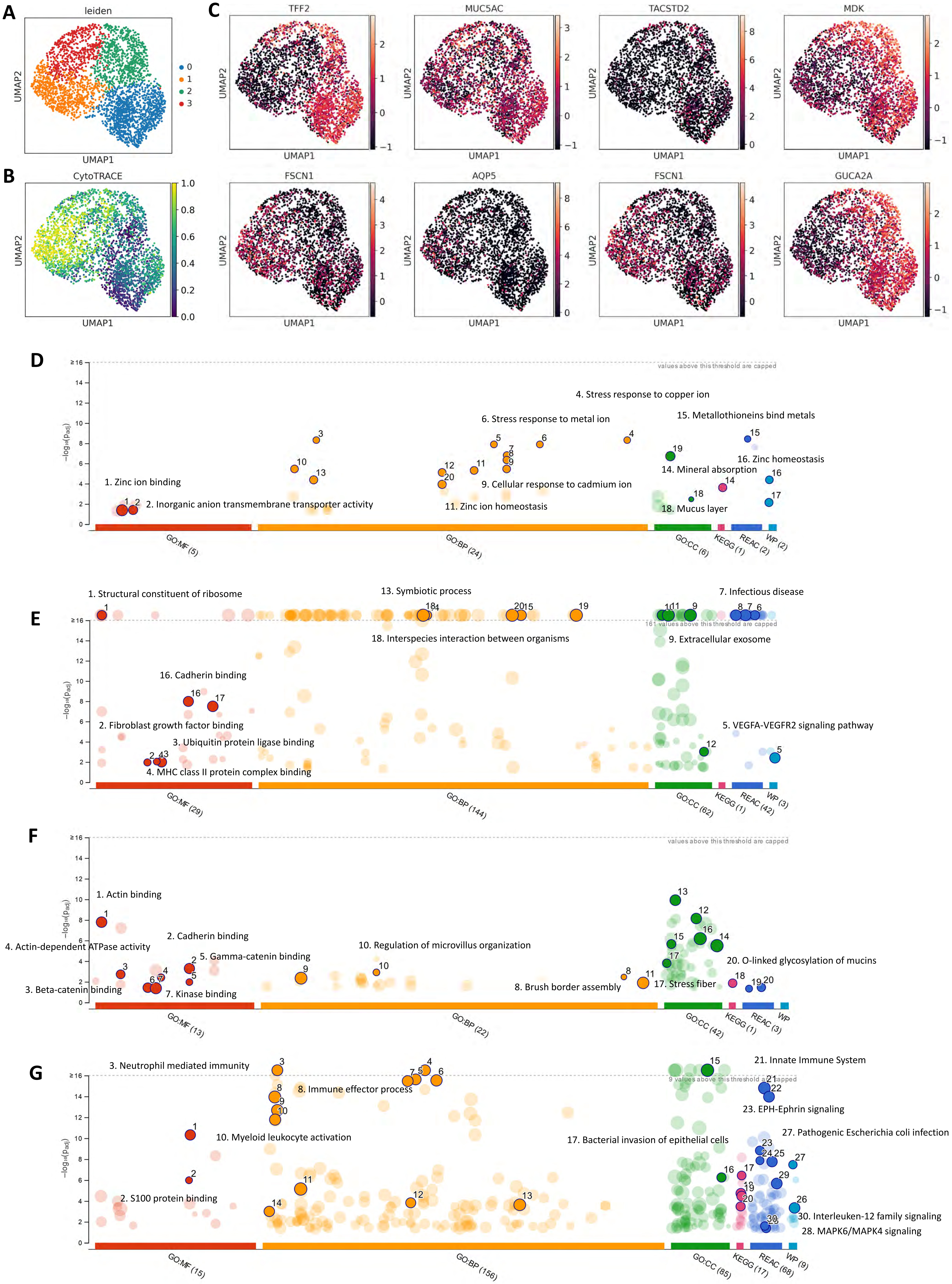

A
leiden
UMAP1
UMAP2
0
1
2
3
B
CytoTRACE
C
TFF2
MUC5AC
TACSTD2
MDK
FSCN1
AQP5
FSCN1
GUCA2A
D
values above this threshold are capped
4. Stress response to copper ion
6. Stress response to metal ion
15. Metallothioneins bind metals
16. Zinc homeostasis
14. Mineral absorption
1. Zinc ion binding
9. Cellular response to cadmium ion
2. Inorganic anion transmembrane transporter activity
11. Zinc ion homeostasis
18. Mucus layer
GO:MF (5)
GO:BP (24)
GO:CC (6)
KEGG (1)
REAC (2)
WP (2)
E
1. Structural constituent of ribosome
13. Symbiotic process
7. Infectious disease
161 values above this threshold are capped
18. Interspecies interaction between organisms
9. Extracellular exosome
16. Cadherin binding
2. Fibroblast growth factor binding
5. VEGFA-VEGFR2 signaling pathway
3. Ubiquitin protein ligase binding
4. MHC class II protein complex binding
GO:MF (29)
GO:BP (144)
GO:CC (62)
KEGG (1)
REAC (42)
WP (3)
F
values above this threshold are capped
1. Actin binding
2. Cadherin binding
4. Actin-dependent ATPase activity
10. Regulation of microvillus organization
5. Gamma-catenin binding
20. O-linked glycosylation of mucins
3. Beta-catenin binding
7. Kinase binding
8. Brush border assembly
17. Stress fiber
GO:MF (13)
GO:BP (22)
GO:CC (42)
KEGG (1)
REAC (3)
WP
G
3. Neutrophil mediated immunity
21. Innate Immune System
8. Immune effector process
23. EPH-Ephrin signaling
10. Myeloid leukocyte activation
27. Pathogenic Escherichia coli infection
17. Bacterial invasion of epithelial cells
2. S100 protein binding
30. Interleukin-12 family signaling
28. MAPK6/MAPK4 signaling
GO:MF (15)
GO:BP (156)
GO:CC (85)
KEGG (17)
REAC (68)
WP (9)

#### Figure S9

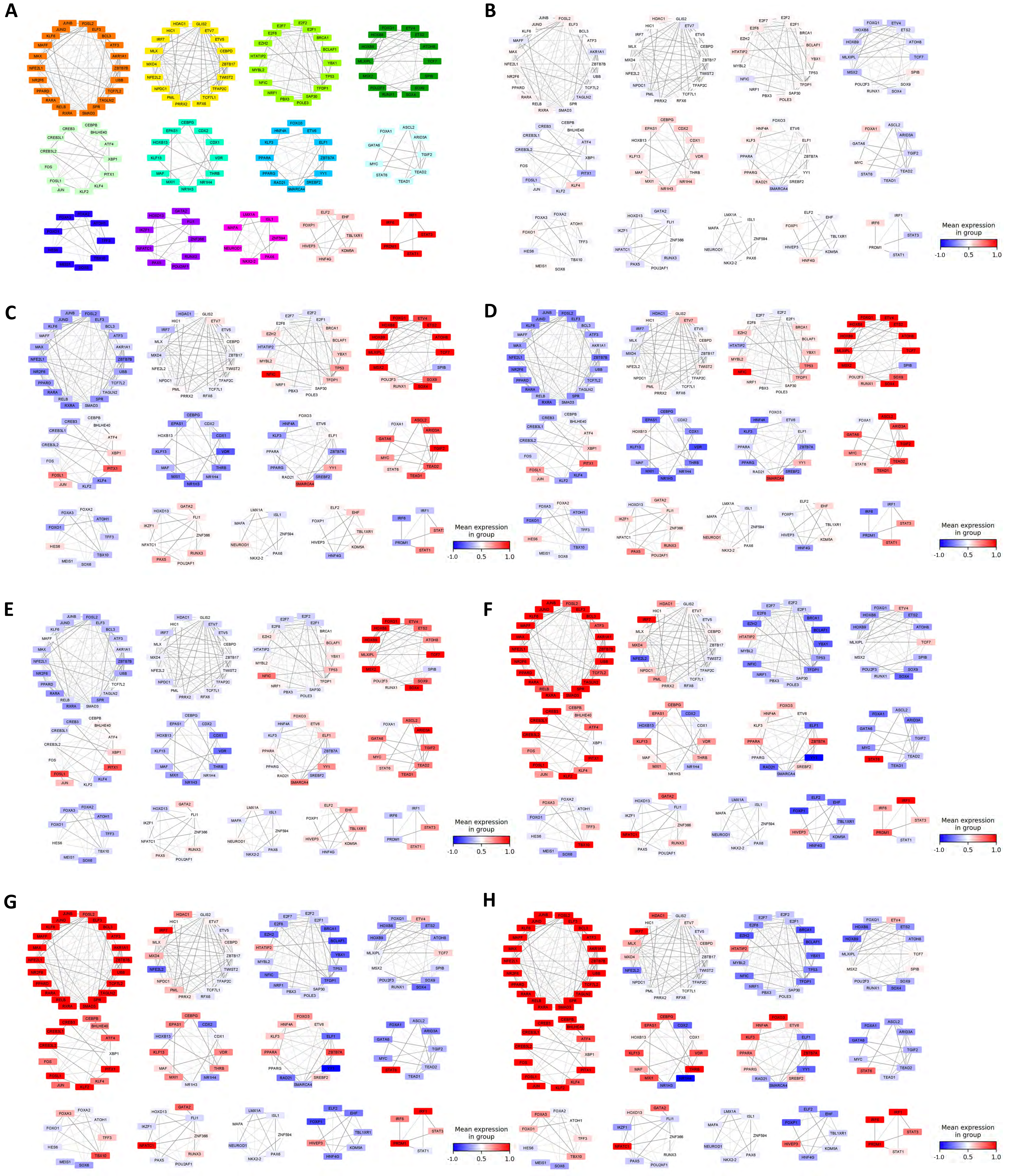

### Figure S10

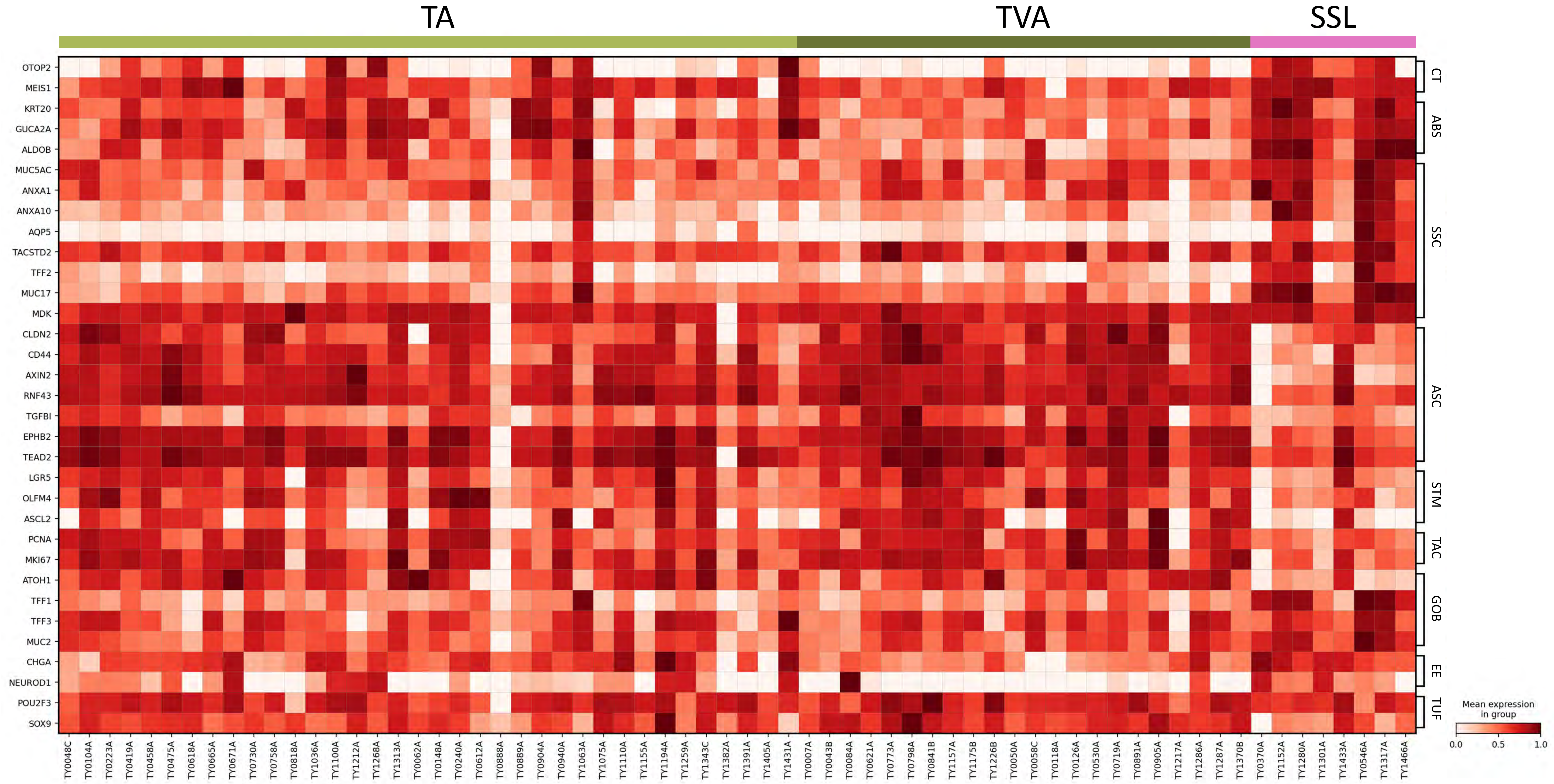
TA
TVA
SSL
Mean expression in group
CT
ABS
SSC
ASC
STM
TAC
GOB
EE
TUF

### Figure S11

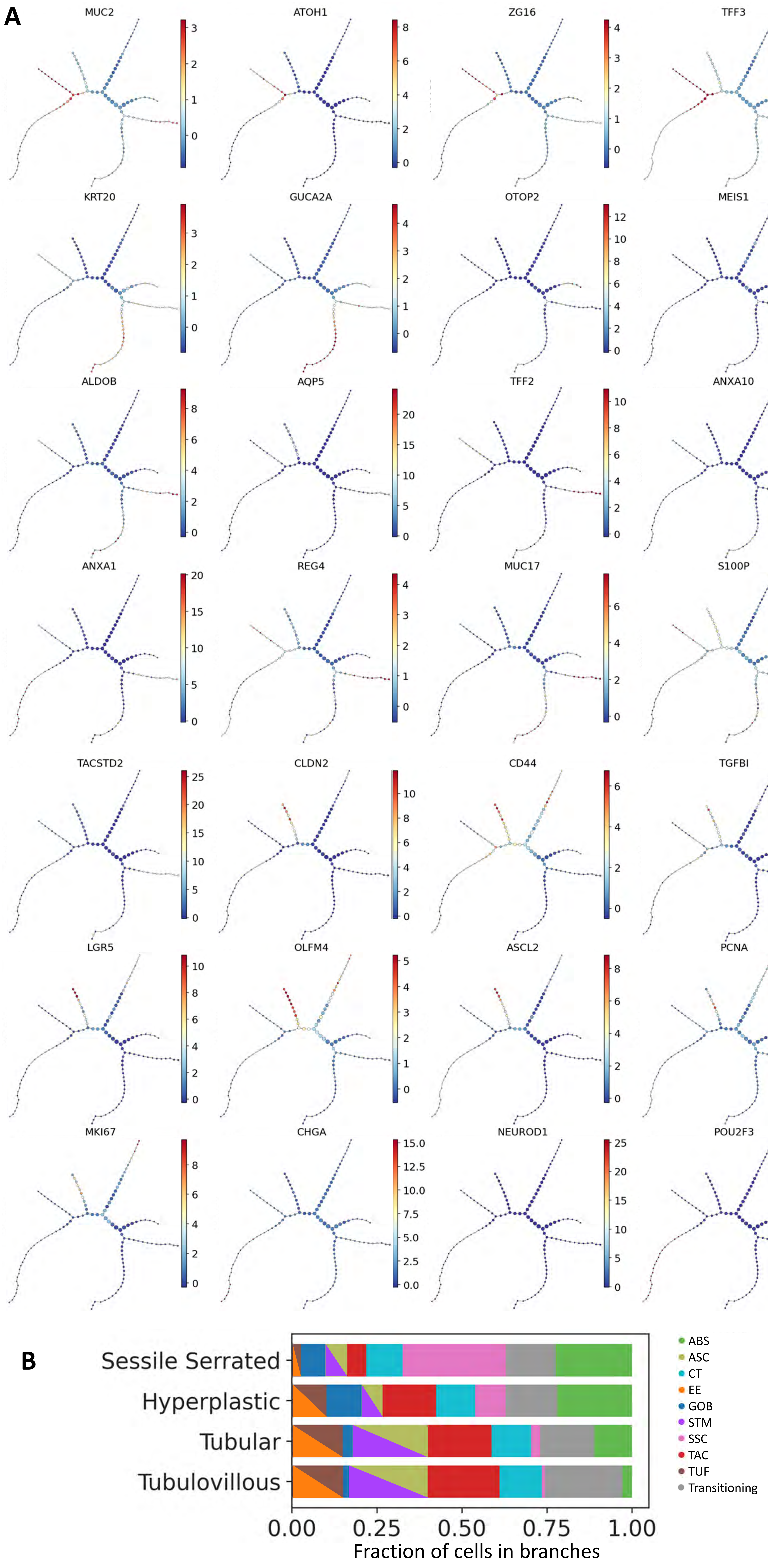
A
MUC2
ATOH1
ZG16
TFF3
KRT20
GUCA2A
OTOP2
MEIS1
ALDOB
AQP5
TFF2
ANXA10
ANXA1
REG4
MUC17
S100P
TACSTD2
CLDN2
CD44
TGFBI
LGR5
OLFM4
ASCL2
PCNA
MKI67
CHGA
NEUROD1
POU2F3
B
Sessile Serrated
Hyperplastic
Tubular
Tubulovillous
0.00
0.25
0.50
0.75
1.00
Fraction of cells in branches
ABS
ASC
CT
EE
GOB
STM
SSC
TAC
TUF
Transitioning

**Figure S12**

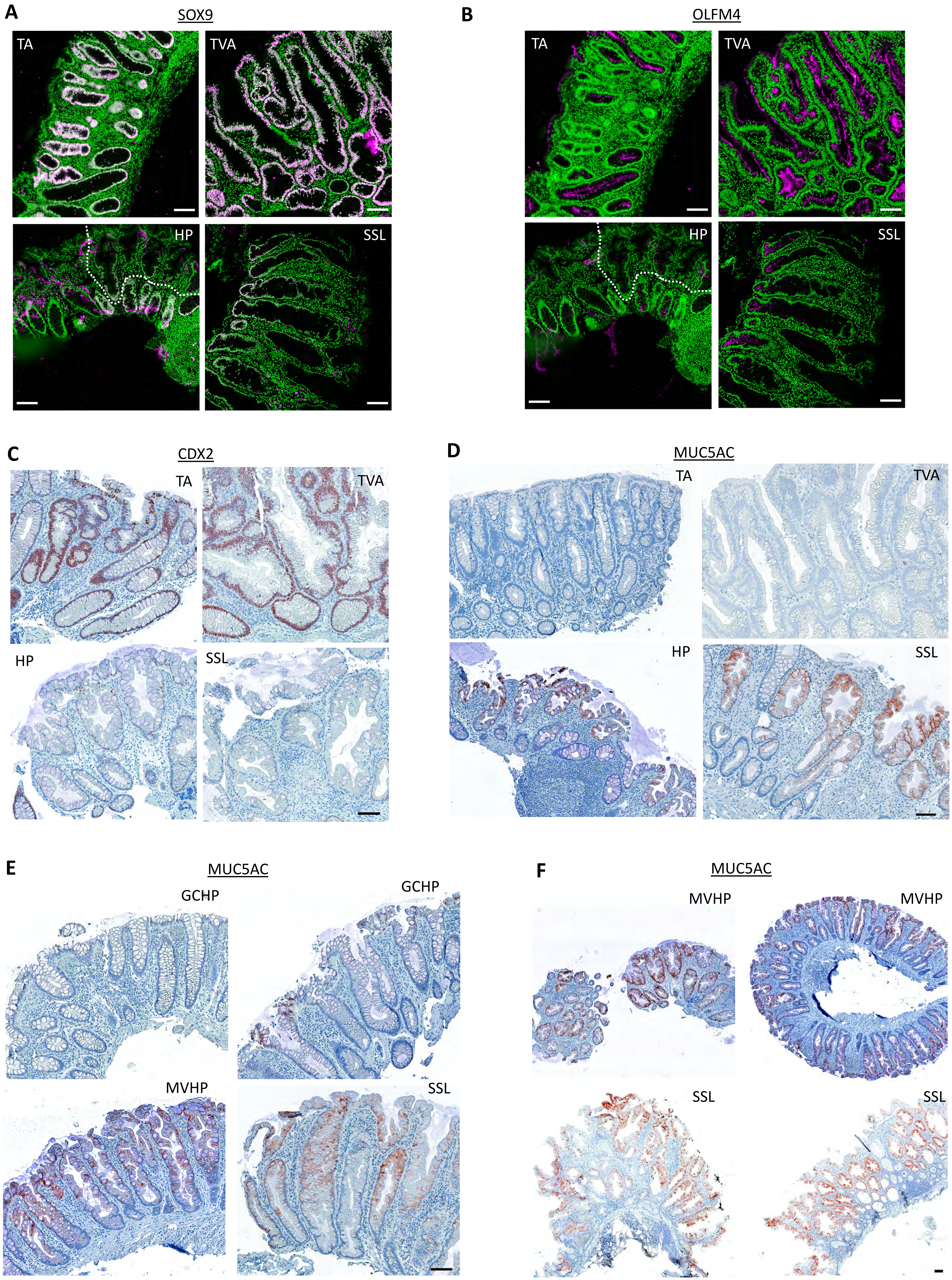
A
SOX9
TA
TVA
HP
SSL
B
OLFM4
TA
TVA
HP
SSL
C
CDX2
TA
TVA
HP
SSL
D
MUC5AC
TA
TVA
HP
SSL
E
MUC5AC
GCHP
GCHP
MVHP
SSL
F
MUC5AC
MVHP
MVHP
SSL
SSL

**Figure S13**

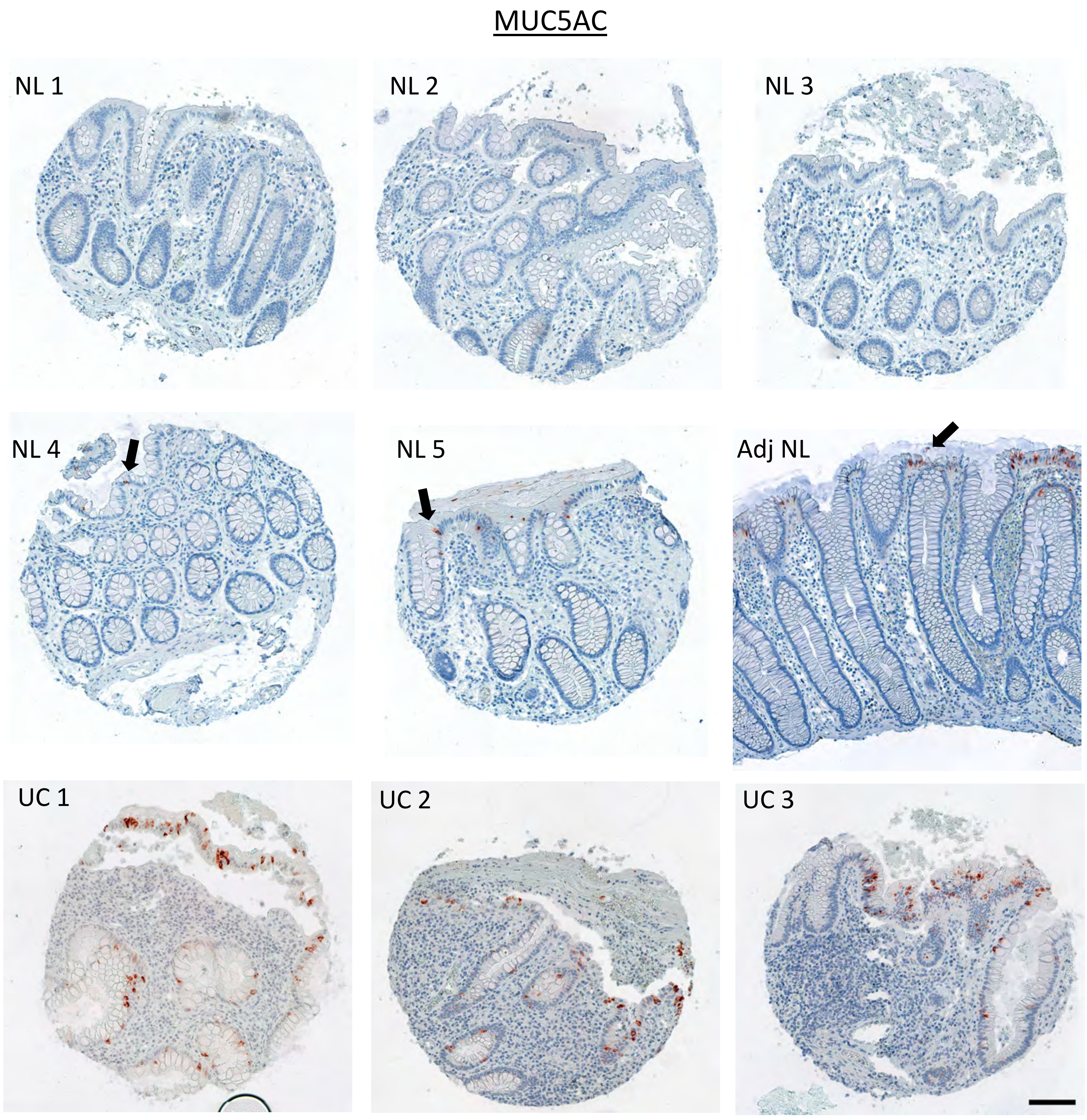

**Figure S14**

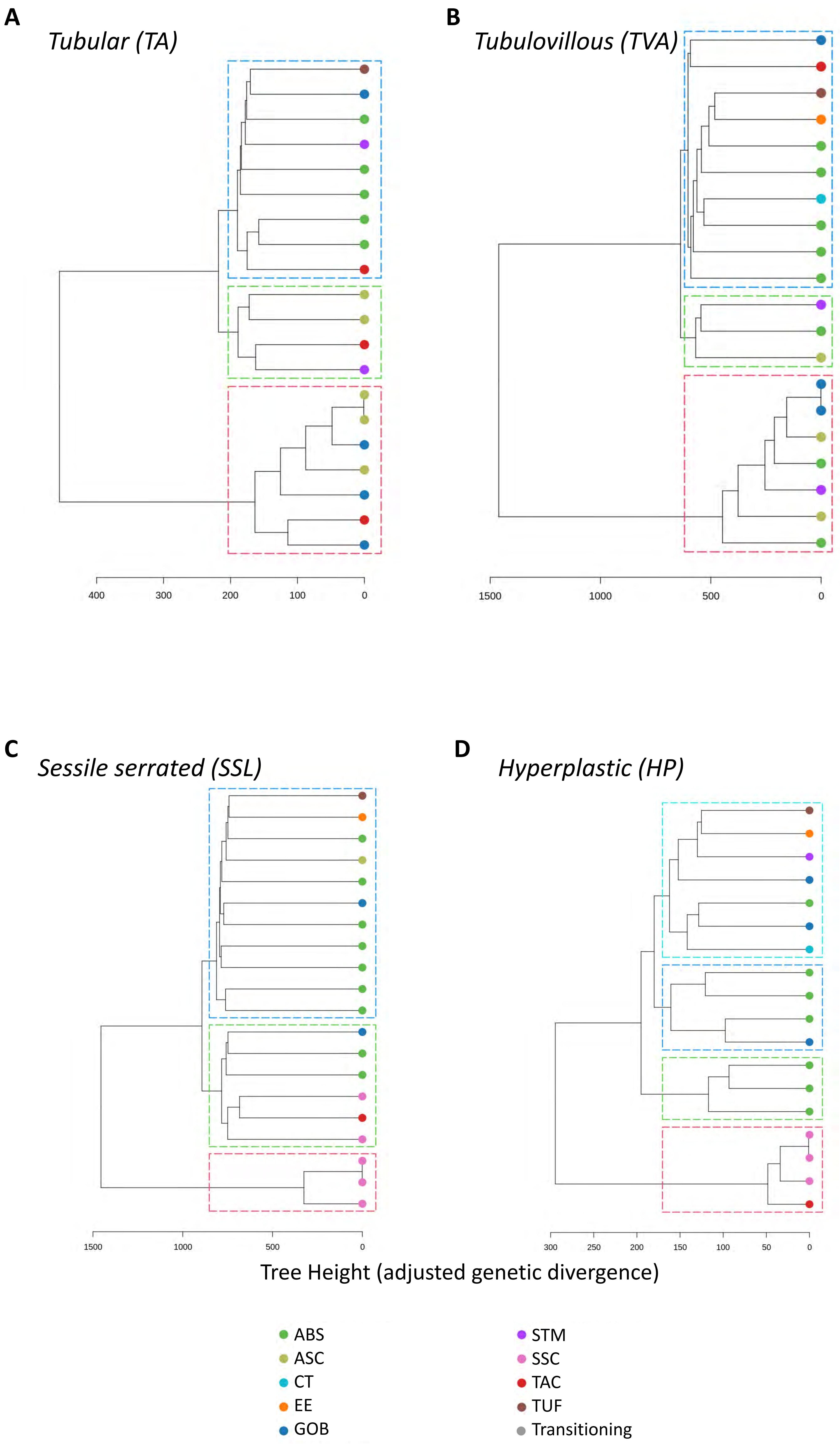
A
Tubular (TA)
400
300
200
100
0
B
Tubulovillous (TVA)
1500
1000
500
0
C
Sessile serrated (SSL)
1500
1000
500
0
D
Hyperplastic (HP)
300
250
200
150
100
50
0
Tree Height (adjusted genetic divergence)
ABS
ASC
CT
EE
GOB
STM
SSC
TAC
TUF
Transitioning

### Figure S15

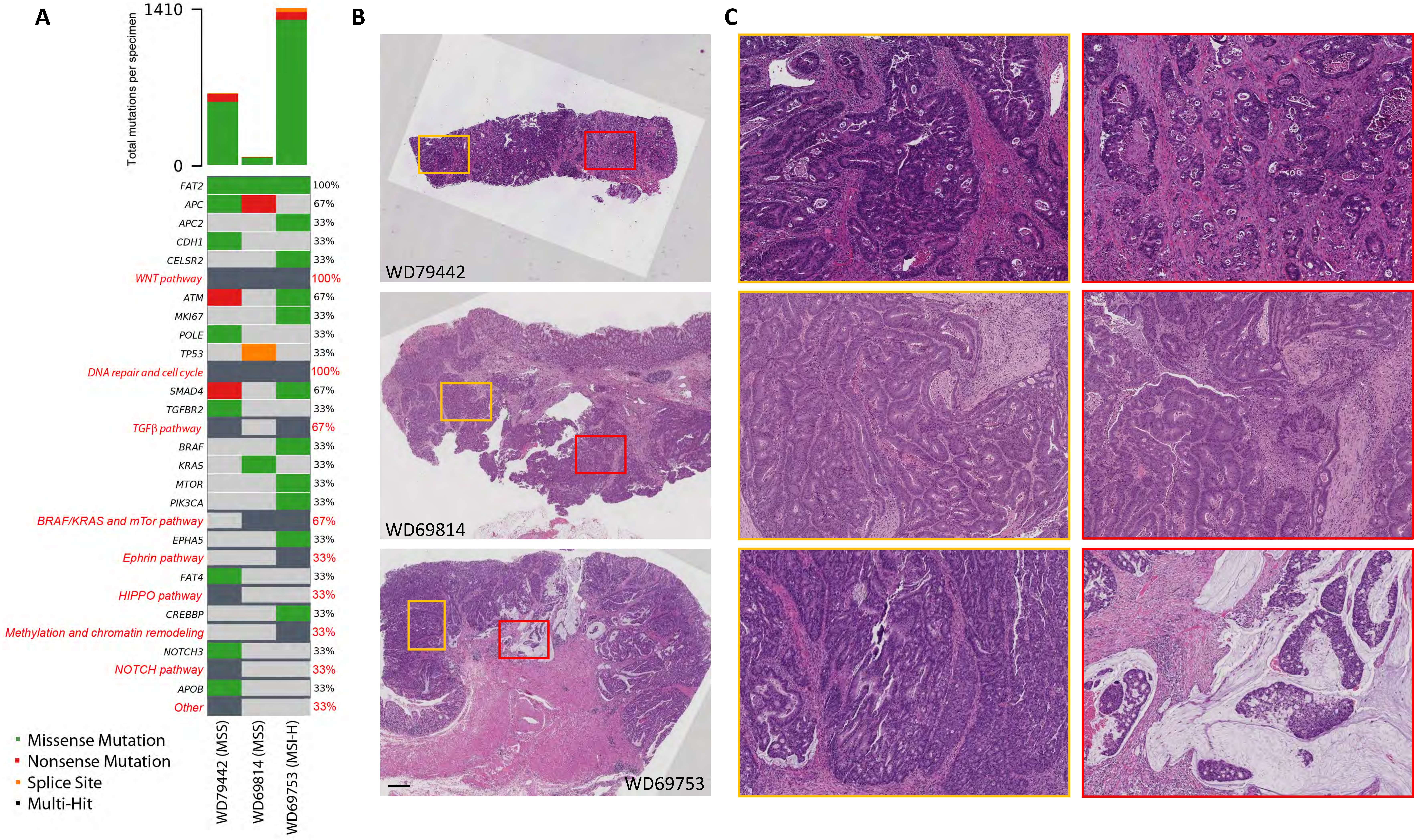
A
B
C
1410
0
Total mutations per specimen
FAT2
APC
APC2
CDH1
CELSR2
WNT pathway
ATM
MKI67
POLE
TP53
DNA repair and cell cycle
SMAD4
TGFBR2
TGFβ pathway
BRAF
KRAS
MTOR
PIK3CA
BRAF/KRAS and mTor pathway
EPHA5
Ephrin pathway
FAT4
HIPPO pathway
CREBBP
Methylation and chromatin remodeling
NOTCH3
NOTCH pathway
APOB
Other
100%
67%
33%
33%
33%
100%
67%
33%
33%
33%
100%
67%
33%
67%
33%
33%
33%
33%
67%
33%
33%
33%
33%
33%
33%
33%
33%
33%
33%
Missense Mutation
Nonsense Mutation
Splice Site
Multi-Hit
WD79442 (MSS)
WD69814 (MSS)
WD69753 (MSI-H)
WD79442
WD69814
WD69753

Figure S16

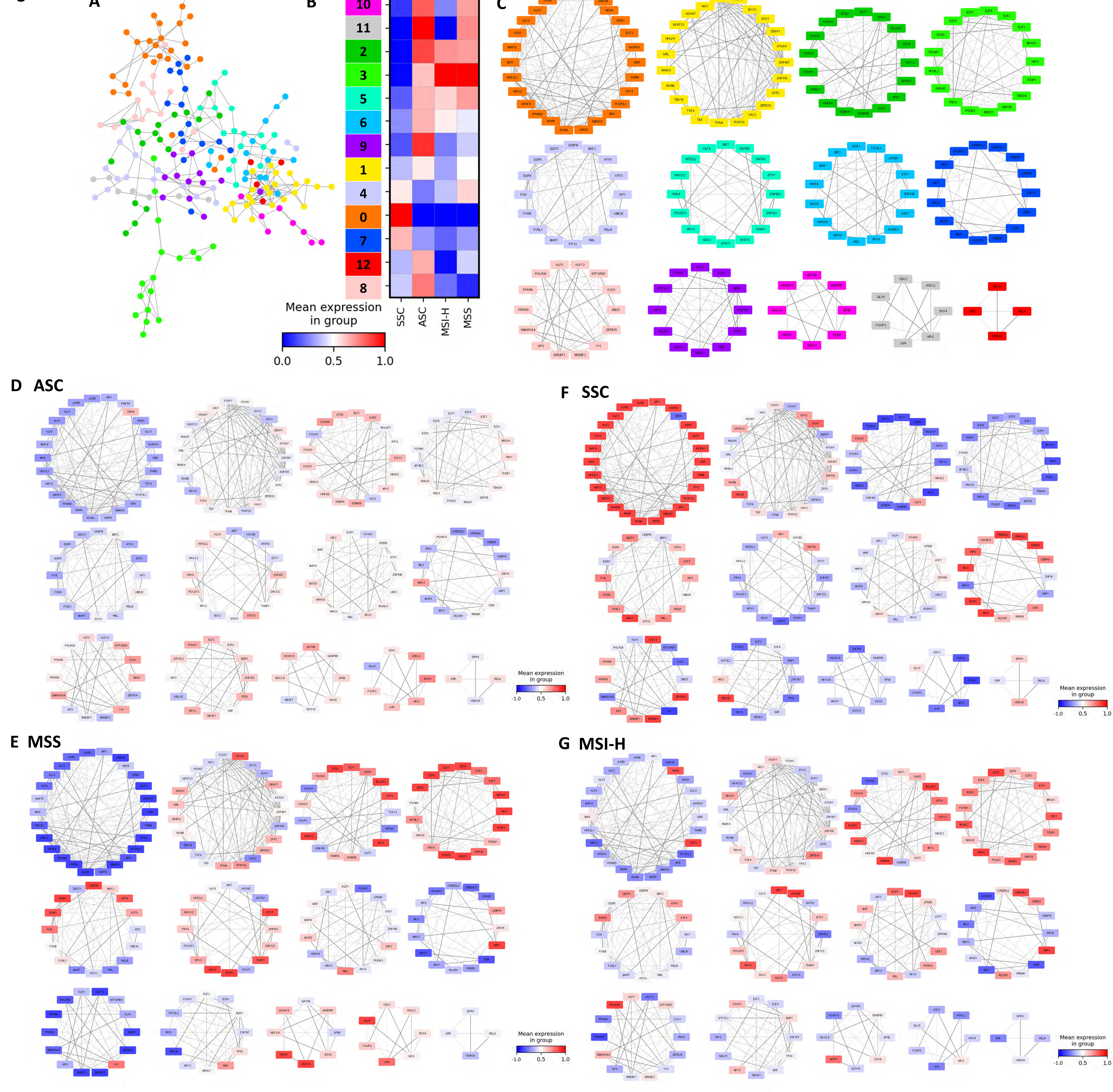
A
B
10
11
2
3
5
6
9
1
4
0
7
12
8
SSC
ASC
MSI-H
MSS
Mean expression in group
0.0
0.5
1.0
C
D ASC
F SSC
E MSS
G MSI-H
Mean expression in group
-1.0
0.5
1.0

### Figure S17

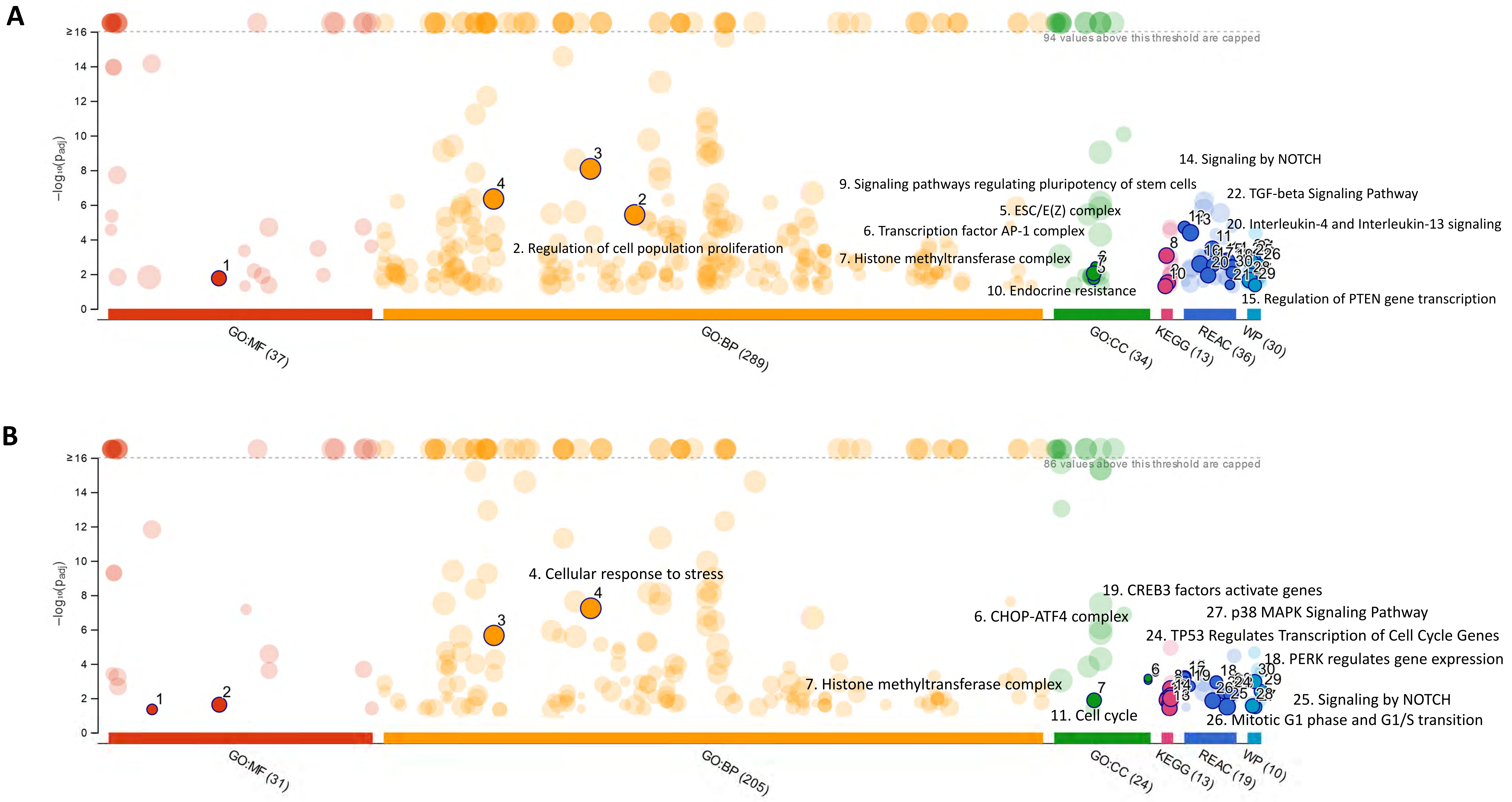
A
94 values above this threshold are capped
−log10(p_adj)
14. Signaling by NOTCH
9. Signaling pathways regulating pluripotency of stem cells
22. TGF-beta Signaling Pathway
5. ESC/E(Z) complex
20. Interleukin-4 and Interleukin-13 signaling
6. Transcription factor AP-1 complex
2. Regulation of cell population proliferation
7. Histone methyltransferase complex
10. Endocrine resistance
15. Regulation of PTEN gene transcription
GO:MF (37)
GO:BP (289)
GO:CC (34)
KEGG (13)
REAC (36)
WP (30)
B
86 values above this threshold are capped
−log10(p_adj)
4. Cellular response to stress
19. CREB3 factors activate genes
27. p38 MAPK Signaling Pathway
6. CHOP-ATF4 complex
24. TP53 Regulates Transcription of Cell Cycle Genes
18. PERK regulates gene expression
7. Histone methyltransferase complex
25. Signaling by NOTCH
11. Cell cycle
26. Mitotic G1 phase and G1/S transition
GO:MF (31)
GO:BP (205)
GO:CC (24)
KEGG (13)
REAC (19)
WP (10)

#### Figure S18

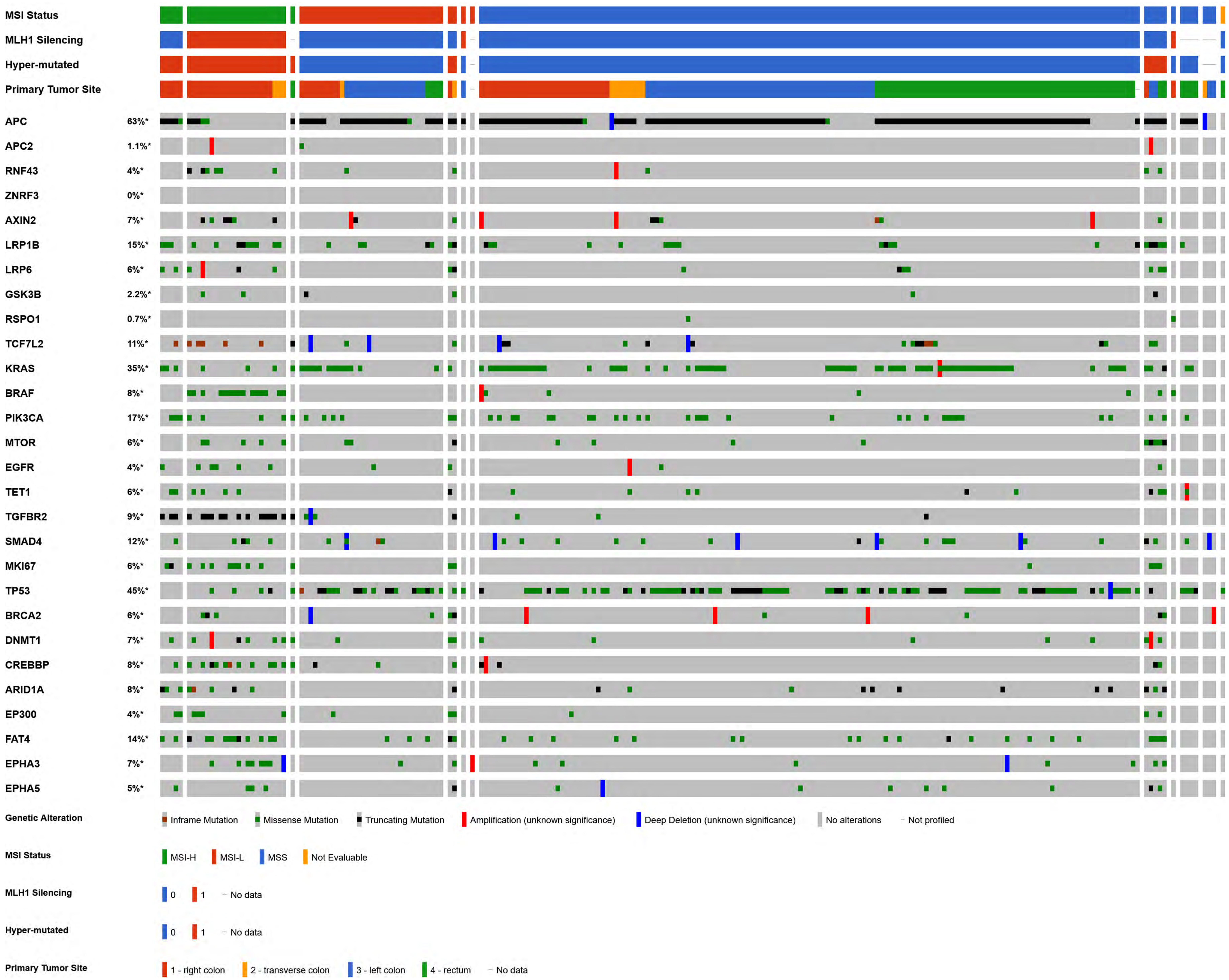
MSI Status
MLH1 Silencing
Hyper-mutated
Primary Tumor Site
APC 63%*
APC2 1.1%*
RNF43 4%*
ZNRF3 0%*
AXIN2 7%*
LRP1B 15%*
LRP6 6%*
GSK3B 2.2%*
RSPO1 0.7%*
TCF7L2 11%*
KRAS 35%*
BRAF 8%*
PIK3CA 17%*
MTOR 6%*
EGFR 4%*
TET1 6%*
TGFBR2 9%*
SMAD4 12%*
MKI67 6%*
TP53 45%*
BRCA2 6%*
DNMT1 7%*
CREBBP 8%*
ARID1A 8%*
EP300 4%*
FAT4 14%*
EPHA3 7%*
EPHA5 5%*
Genetic Alteration
Inframe Mutation
Missense Mutation
Truncating Mutation
Amplification (unknown significance)
Deep Deletion (unknown significance)
No alterations
Not profiled
MSI Status
MSI-H
MSI-L
MSS
Not Evaluable
MLH1 Silencing
0
1
No data
Hyper-mutated
0
1
No data
Primary Tumor Site
1 - right colon
2 - transverse colon
3 - left colon
4 - rectum
No data

**Figure S19**

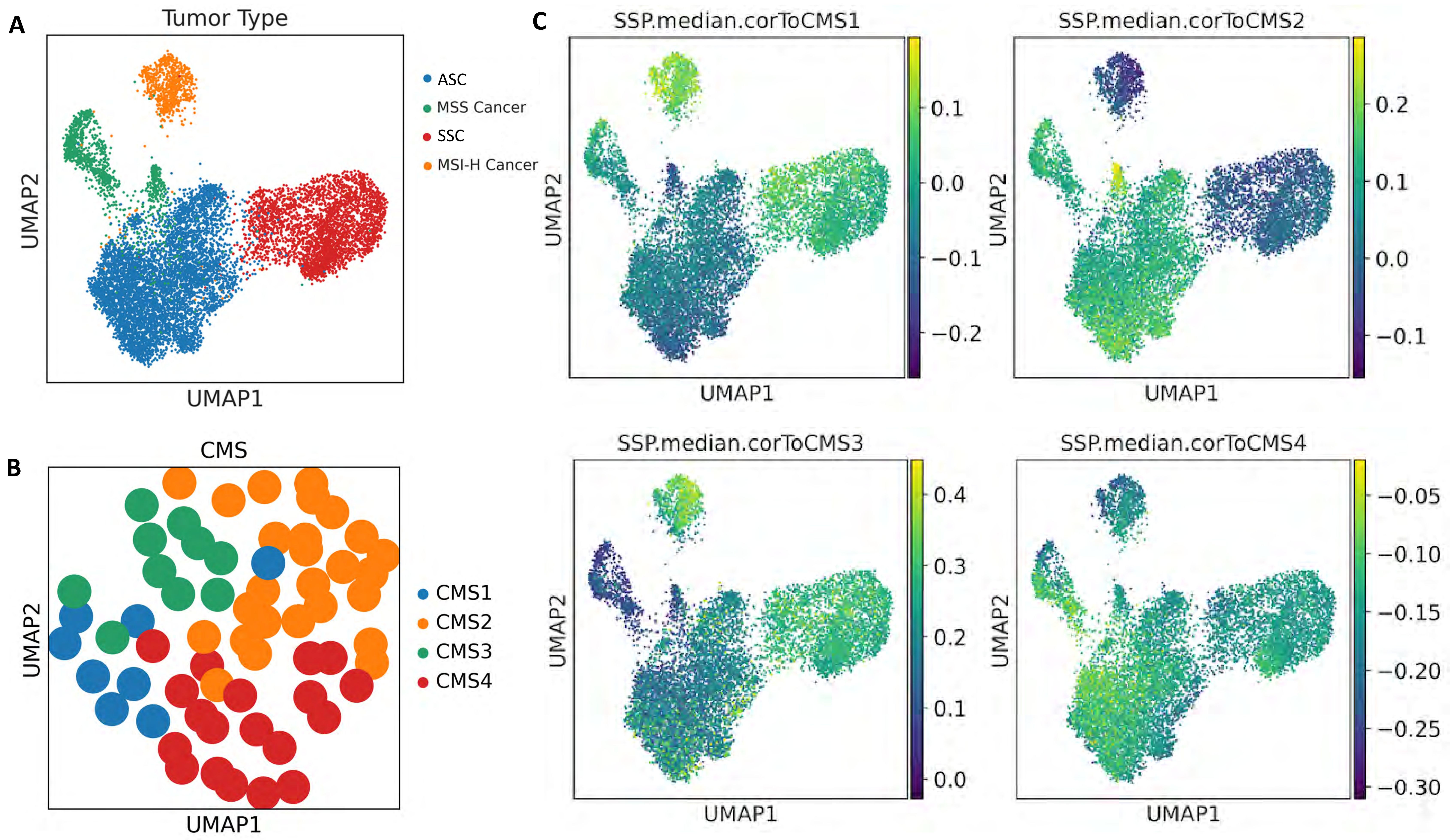

A
Tumor Type
ASC
MSS Cancer
SSC
MSI-H Cancer
UMAP2
UMAP1
B
CMS
CMS1
CMS2
CMS3
CMS4
UMAP2
UMAP1
C
SSP.median.corToCMS1
0.1
0.0
−0.1
−0.2
SSP.median.corToCMS2
0.2
0.1
0.0
−0.1
SSP.median.corToCMS3
0.4
0.3
0.2
0.1
0.0
SSP.median.corToCMS4
−0.05
−0.10
−0.15
−0.20
−0.25
−0.30
UMAP2
UMAP1

**Figure S20**

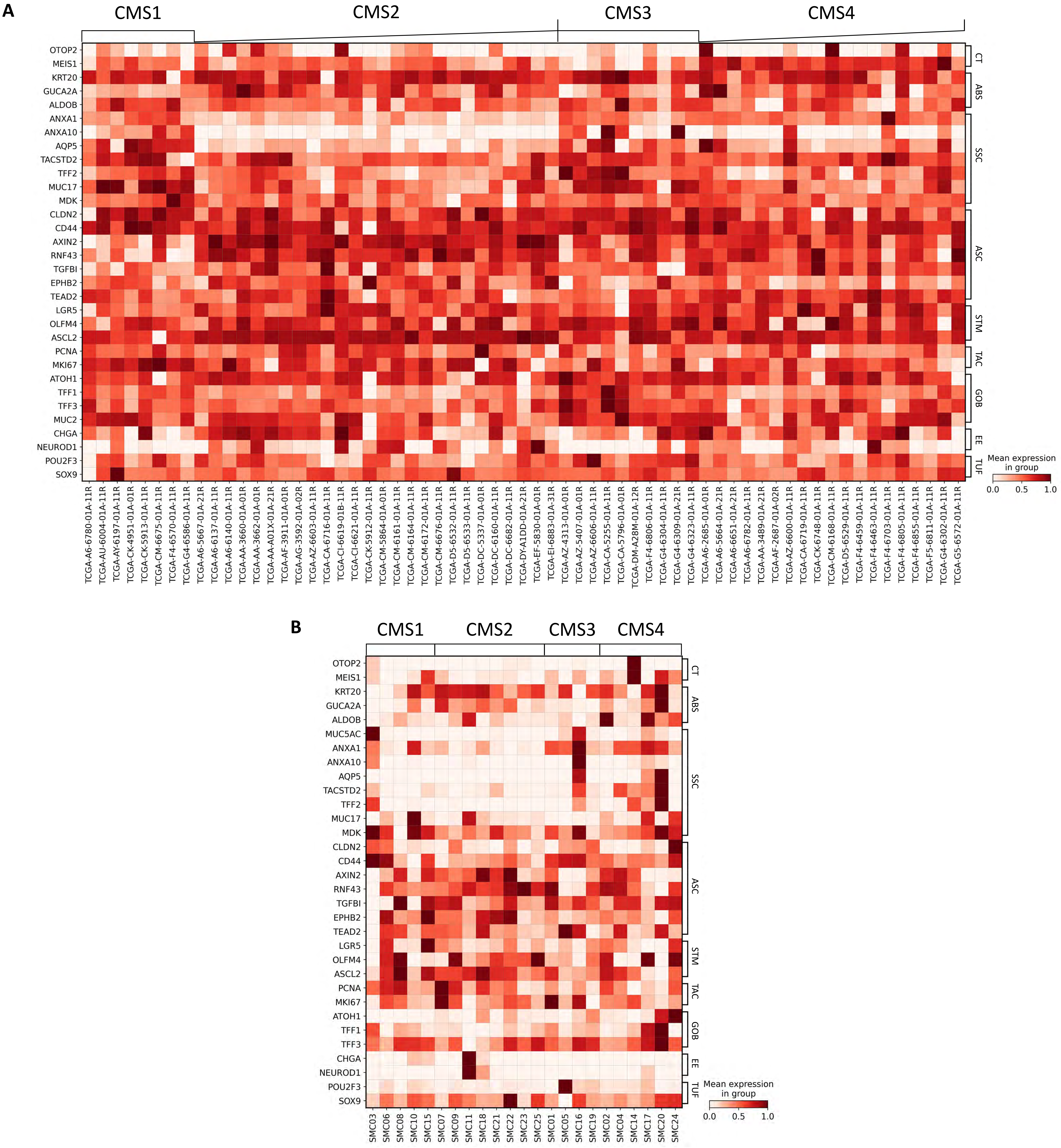
A
CMS1
CMS2
CMS3
CMS4
OTOP2
MEIS1
KRT20
GUCA2A
ALDOB
ANXA1
ANXA10
AQP5
TACSTD2
TFF2
MUC17
MDK
CLDN2
CD44
AXIN2
RNF43
TGFBI
EPHB2
TEAD2
LGR5
OLFM4
ASCL2
PCNA
MKI67
ATOH1
TFF1
TFF3
MUC2
CHGA
NEUROD1
POU2F3
SOX9
CT
ABS
SSC
ASC
STM
TAC
GOB
EE
TUF
Mean expression in group
0.0
0.5
1.0
TCGA-A6-6780-01A-11R
TCGA-AU-6004-01A-11R
TCGA-AY-6197-01A-11R
TCGA-CK-4951-01A-01R
TCGA-CK-5913-01A-11R
TCGA-CM-6675-01A-11R
TCGA-F4-6570-01A-11R
TCGA-G4-6586-01A-11R
TCGA-A6-5667-01A-21R
TCGA-A6-6137-01A-11R
TCGA-A6-6140-01A-11R
TCGA-AA-3660-01A-01R
TCGA-AA-3662-01A-01R
TCGA-AA-A01X-01A-21R
TCGA-AF-3911-01A-01R
TCGA-AG-3592-01A-02R
TCGA-AZ-6603-01A-11R
TCGA-CA-6716-01A-11R
TCGA-CI-6619-01B-11R
TCGA-CI-6621-01A-11R
TCGA-CK-5912-01A-11R
TCGA-CM-5864-01A-01R
TCGA-CM-6161-01A-11R
TCGA-CM-6164-01A-11R
TCGA-CM-6172-01A-11R
TCGA-CM-6676-01A-11R
TCGA-D5-6532-01A-11R
TCGA-D5-6533-01A-11R
TCGA-DC-5337-01A-01R
TCGA-DC-6160-01A-11R
TCGA-DC-6682-01A-11R
TCGA-DY-A1DD-01A-21R
TCGA-EF-5830-01A-01R
TCGA-EI-6883-01A-31R
TCGA-AZ-4313-01A-01R
TCGA-AZ-5407-01A-01R
TCGA-AZ-6606-01A-11R
TCGA-CA-5255-01A-11R
TCGA-CA-5796-01A-01R
TCGA-DM-A28M-01A-12R
TCGA-F4-6806-01A-11R
TCGA-G4-6304-01A-11R
TCGA-G4-6309-01A-21R
TCGA-G4-6323-01A-11R
TCGA-A6-2685-01A-01R
TCGA-A6-5664-01A-21R
TCGA-A6-6651-01A-21R
TCGA-A6-6782-01A-11R
TCGA-AA-3489-01A-21R
TCGA-AF-2687-01A-02R
TCGA-AZ-6600-01A-11R
TCGA-CA-6719-01A-11R
TCGA-CK-6748-01A-11R
TCGA-CM-6168-01A-11R
TCGA-D5-6529-01A-11R
TCGA-F4-6459-01A-11R
TCGA-F4-6463-01A-11R
TCGA-F4-6703-01A-11R
TCGA-F4-6805-01A-11R
TCGA-F4-6855-01A-11R
TCGA-F5-6811-01A-11R
TCGA-G4-6302-01A-11R
TCGA-G5-6572-01A-11R
B
CMS1
CMS2
CMS3
CMS4
OTOP2
MEIS1
KRT20
GUCA2A
ALDOB
MUC5AC
ANXA1
ANXA10
AQP5
TACSTD2
TFF2
MUC17
MDK
CLDN2
CD44
AXIN2
RNF43
TGFBI
EPHB2
TEAD2
LGR5
OLFM4
ASCL2
PCNA
MKI67
ATOH1
TFF1
TFF3
CHGA
NEUROD1
POU2F3
SOX9
CT
ABS
SSC
ASC
STM
TAC
GOB
EE
TUF
Mean expression in group
0.0
0.5
1.0
SMC03
SMC06
SMC08
SMC10
SMC15
SMC07
SMC09
SMC11
SMC18
SMC21
SMC22
SMC23
SMC25
SMC01
SMC05
SMC16
SMC19
SMC02
SMC04
SMC14
SMC17
SMC20
SMC24

Figure S21

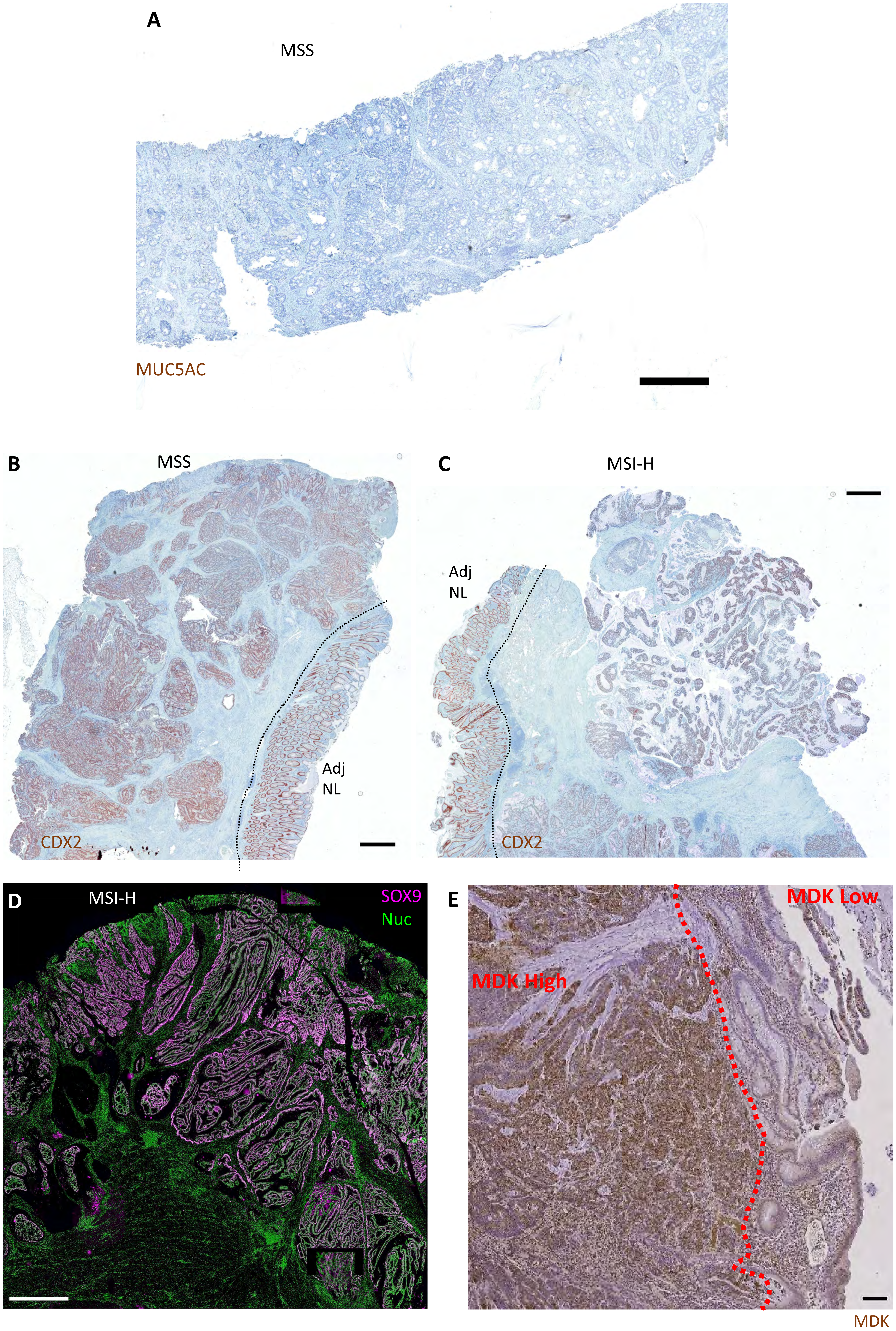
A
MSS
MUC5AC
B
MSS
Adj
NL
CDX2
C
MSI-H
Adj
NL
CDX2
D
MSI-H
SOX9
Nuc
E
MDK Low
MDK High
MDK

**Figure S22**

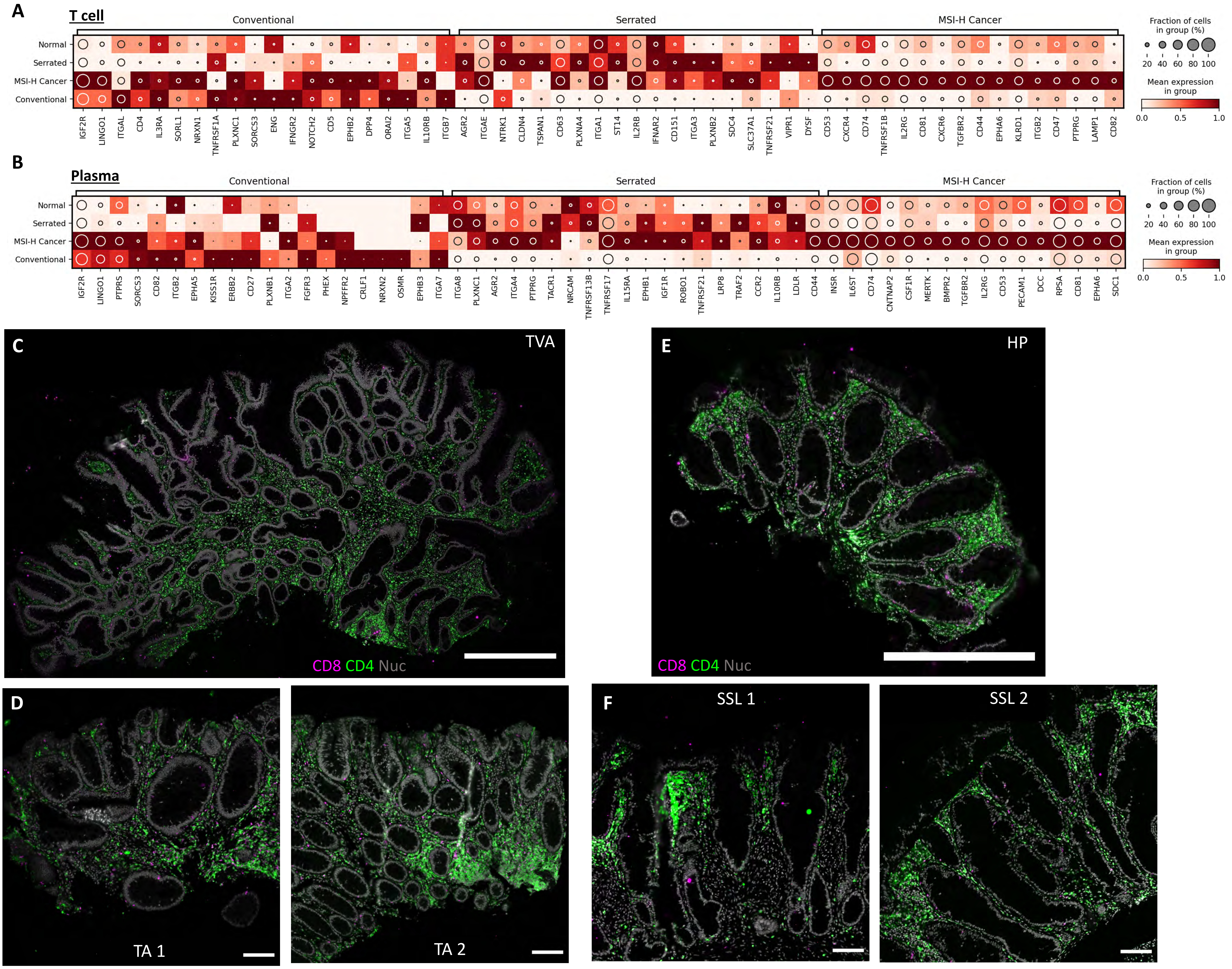
A
T cell
Conventional
Serrated
MSI-H Cancer
Normal
Serrated
MSI-H Cancer
Conventional
IGF2R LINGO1 ITGAL CD4 IL3RA SORL1 NRXN1 TNFRSF1A PLXNC1 SORCS3 ENG IFNGR2 NOTCH2 CD5 EPHB2 DPP4 ORAI2 ITGA5 IL10RB ITGB7 AGR2 ITGAE NTRK1 CLDN4 TSPAN1 CD63 PLXNA4 ITGA1 ST14 IL2RB IFNAR2 CD151 ITGA3 PLXNB2 SDC4 SLC37A1 TNFRSF21 VIPR1 DYSF CD53 CXCR4 CD74 TNFRSF1B IL2RG CD81 CXCR6 TGFBR2 CD44 EPHA6 KLRD1 ITGB2 CD47 PTPRG LAMP1 CD82
Fraction of cells in group (%)
20 40 60 80 100
Mean expression in group
0.0 0.5 1.0
B
Plasma
Conventional
Serrated
MSI-H Cancer
Normal
Serrated
MSI-H Cancer
Conventional
IGF2R LINGO1 PTPRS SORCS3 CD82 ITGB2 EPHA5 KISS1R ERBB2 CD27 PLXNB1 ITGA2 FGFR3 PHEX NPFFR2 CRLF1 NRXN2 OSMR EPHB3 ITGA7 ITGA8 PLXNC1 AGR2 ITGA4 PTPRG TACR1 NRCAM TNFRSF13B TNFRSF17 IL15RA EPHB1 IGF1R ROBO1 TNFRSF21 LRP8 TRAF2 CCR2 IL10RB LDLR CD44 INSR IL6ST CD74 CNTNAP2 CSF1R MERTK BMPR2 TGFBR2 IL2RG CD53 PECAM1 DCC RPSA CD81 EPHA6 SDC1
Fraction of cells in group (%)
20 40 60 80 100
Mean expression in group
0.0 0.5 1.0
C
TVA
CD8 CD4 Nuc
E
HP
CD8 CD4 Nuc
D
TA 1
TA 2
F
SSL 1
SSL 2

**Figure S23**

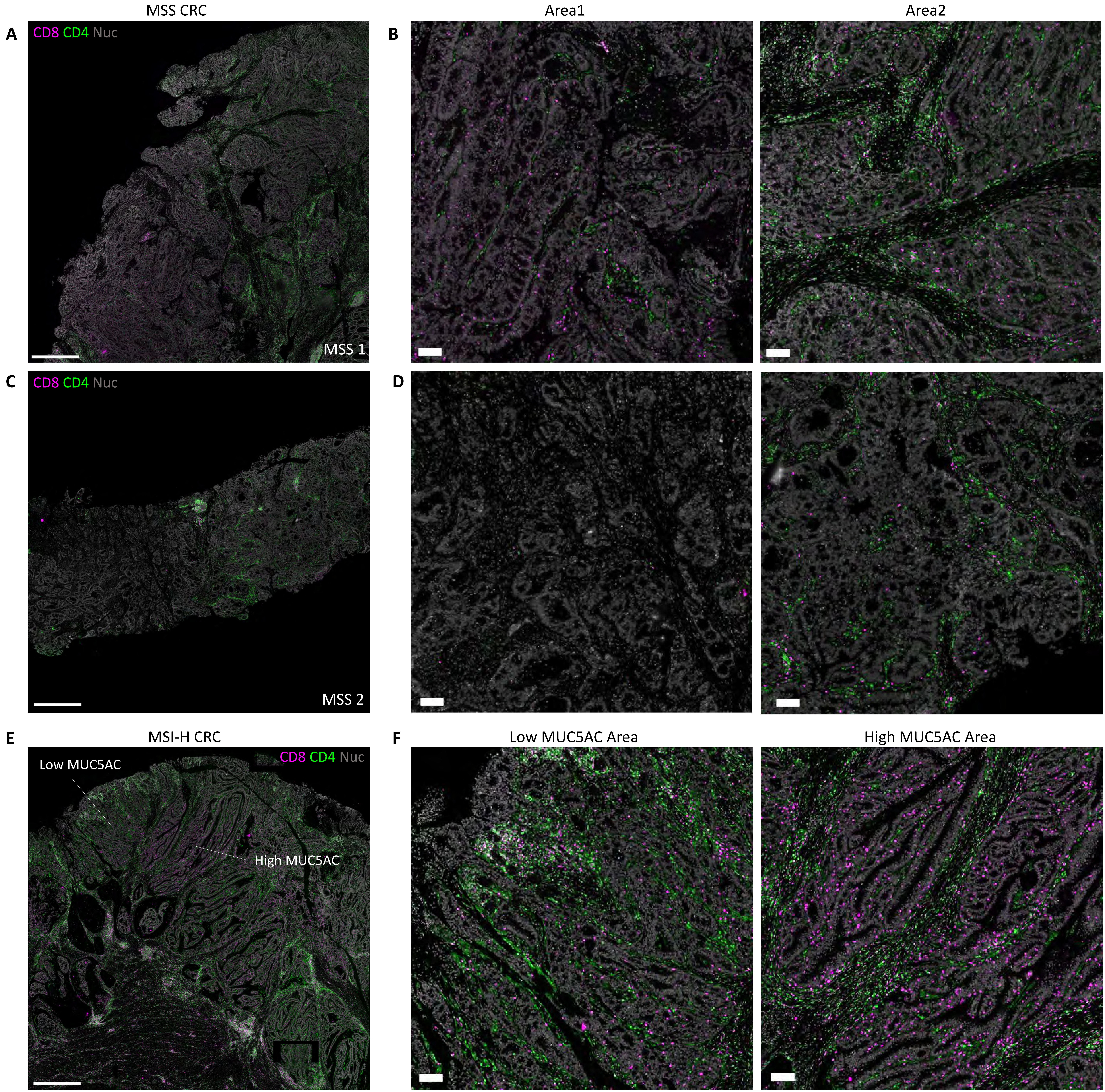

MSS CRC
Area1
Area2
A
CD8 CD4 Nuc
MSS 1
B
C
CD8 CD4 Nuc
MSS 2
D
E
MSI-H CRC
CD8 CD4 Nuc
Low MUC5AC
High MUC5AC
F
Low MUC5AC Area
High MUC5AC Area
