## Supplemental Tables 1-22 for "Human colorectal pre-cancer atlas identifies distinct molecular programs underlying two major subclasses of pre-malignant tumors"

#### Table S1 Tennessee Colorectal Polyp Study participants

| | Included in targeted gene sequencing (n=197) | Included in RNA Sequencing (n=66) |
|---|---|---|
| **Participant characteristics at enrollment** | | |
| Age, y, mean ± std | 59.5 (7.0) | 58.3 (7.0) |
| Gender | | |
| Female, n (%) | 62 (31.5) | 28 (42.4) |
| Male, n (%) | 135 (68.5) | 38 (57.6) |
| Race | | |
| White, n(%) | 170 (86.3) | 54 (81.8) |
| ….Black, n(%) | 20 (10.2) | 11 (16.7) |
| Other, n(%) | 4 (2.0) | 1 (1.5) |
| Unknown, n(%) | 3 (1.5) | |
| Polyp types at enrollment procedure, n (%) | | |
| Any AD | | |
| Tubular | 193 (97.8) | 61 (92.4) |
| Tubulovillous/Villous | 170 (86.2) | 47 (71.2) |
| Any SSL | 50 (25.4) | 24 (36.4) |
| Any HP | 10 (5.1) | 12 (18.2) |
| | 37 (18.8) | 10 (15.2) |
| Subtype of the sequenced polyps, n | | |
| Tubular AD | 217 | 36 |
| Tubulovillous AD | 50 | 22 |
| SSL | 7 | 8 |

[a] Large defined as ≥ 1 cm; Advanced defined as large, tubulovillous, villous, or high grade dysplasia; Proximal defined as cecum, ascending colon, hepatic flexure, transverse colon; Distal defined as splenic flexure, descending colon, sigmoid colon, or rectum

Table S2. Characteristics of COLON MAP study participants and polyp samples

| | | COLON MAP SAMPLED POLYP CASE GROUP | | | | |
|---|---|---|---|---|---|---|
| **Characteristic** | **Total (n=30)** | **No Sampled Polyp (n=3)** | **Hyperplastic Polyp (HP) (n=8)** | **Conventional Adenoma (AD) (n=14)** | **Sessile Serrated Lesion (SSL) (n=3)** | **SSL/ Tubular (n=2)** |
| Age, mean ± std | 60.6 (8.0) | 57.0 (7.0) | 61.1 (9.1) | 60.4 (8.3) | 61.0 (9.6) | 64.5 (3.5) |
| Female, n (%) | 12 (40%) | 2 (67%) | 3 (38%) | 4 (29%) | 2 (67%) | 1 (50%) |
| Race/Ethnicity, n (%) | | | | | | |
| White, non-Latinx | 21 (70%) | 2 (67%) | 5 (63%) | 10 (71%) | 2 (67%) | 2 (100%) |
| Black, non-Latinx | 6 (20%) | 1 (33%) | 2 (25%) | 3 (21%) | 0 | 0 |
| White, Latinx | 1 (3%) | 0 | 0 | 0 | 1 (33%) | 0 |
| More than one race, non-Latinx | 1 (3%) | 0 | 0 | 1 (7%) | 0 | 0 |
| Unknown, Latinx | 1 (3%) | 0 | 1 (13%) | 0 | 0 | 0 |
| Colonoscopy indication, n(%) | | | | | | |
| Screening | 15 (50%) | 2 (67%) | 4 (50%) | 5 (36%) | 3 (100%) | 1 (50%) |
| Surveillance | 15 (50%) | 1 (33%) | 4 (50%) | 9 (64%) | 0 | 1 (50%) |
| Family history of colorectal cancer in a first-degree relative, n (%) | 2 (7%) | 0 | 0 | 1 (7%) | 1 (33%) | 0 |
| Yes | 21 (70%) | 3 (100%) | 7 (88%) | 8 (57%) | 1 (33%) | 2 (100%) |
| No | 7 (23%) | 0 | 1 (13%) | 5 (36%) | 1 (33%) | 0 |
| Unknown | | | | | | |
| Personal history of polyps, % | | | | | | |
| No history | 12 (40%) | 2 (67%) | 4 (50%) | 5 (36%) | 1 (33%) | 0 |
| AD | 14 (47%) | 1 (33%) | 4 (50%) | 8 (57%) | 0 | 1 (50%) |
| SSL | 3 (10%) | 1 (33%) | 0 | 1 (7%) | 0 | 1 (50%) |
| HP | 5 (17%) | 1 (33%) | 2 (25%) | 0 | 0 | 2 (100%) |
| Unknown | 3 (10%) | 0 | 0 | 1 (7%) | 2 (67%) | 0 |
| Polyp types at enrollment procedure, n (%) | | | | | | |
| Any AD | 23 (77%) | 2 (67%) | 3 (38%) | 14 (100%) | 2 (67%) | 2 (100%) |
| Tubular | 21 (70%) | 2 (67%) | 3 (38%) | 13 (93%) | 1 (33%) | 2 (100%) |
| Tubulovillous | 4 (13%) | 0 | 0 | 3 (21%) | 1 (33%) | 0 |
| Villous | 0 | 0 | 0 | 0 | 0 | 0 |
| Large | 5 (17%) | 0 | 0 | 4 (29%) | 1 (33%) | 0 |
| High-grade dysplasia | 1 (3%) | 0 | 0 | 1 (7%) | 0 | 0 |
| Advanced | 6 (20%) | 0 | 0 | 5 (36%) | 1 (33%) | 0 |

| Characteristic | Total (n=30) | COLON MAP SAMPLED POLYP CASE GROUP<br>No Sampled Polyp (n=3) | Hyperplastic Polyp (HP) (n=8) | Conventional Adenoma (AD) (n=14) | Sessile Serrated Lesion (SSL) (n=3) | SSL/ Tubular (n=2) |
|---|---|---|---|---|---|---|
| Proximal | 16 (53%) | 1 (33%) | 2 (25%) | 11 (79%) | 0 | 2 (100%) |
| Distal | 14 (47%) | 1 (33%) | 2 (25%) | 9 (64%) | 2 (67%) | 0 |
| Any SSL | 6 (20%) | 1 (33%) | 1 (13%) | 0 | 3 (100%) | 1 (50%) |
| Large | 2 (7%) | 0 | 0 | 0 | 2 (67%) | 0 |
| High-grade dysplasia | 0 | 0 | 0 | 0 | 0 | 0 |
| Proximal | 4 (13%) | 0 | 1 (13%) | 0 | 2 (67%) | 1 (50%) |
| Distal | 3 (10%) | 1 (33%) | 0 | 0 | 1 (33%) | 1 (50%) |
| Any HP | 16 (53%) | 1 (33%) | 8 (100%) | 5 (36%) | 1 (33%) | 1 (50%) |
| Goblet cell HP | 5 (17%) | 1 (33%) | 1 (13%) | 3 (21%) | 0 | 0 |
| Microvesicular HP | 9 (30%) | 0 | 5 (63%) | 3 (21%) | 1 (33%) | 0 |
| Large | 0 | 0 | 0 | 0 | 0 | 0 |
| Proximal | 4 (13%) | 0 | 1 (13%) | 2 (14%) | 1 (33%) | 0 |
| Distal | 13 (43%) | 1 (33%) | 7 (88%) | 3 (21%) | 1 (33%) | 1 (50%) |
| Features of the index polyp, n (%) | | | | | | |
| Polyp location | | | | | | |
| Proximal colon | 14 (47%) | | 1 (13%) | 9 (64%) | 2 (67%) | 2 (100%) |
| Distal colon | 11 (37%) | | 6 (75%) | 4 (29%) | 1 (33%) | 0 |
| Rectum | 2 (7%) | | 1 (13%) | 1 (7%) | 0 | 0 |
| Polyp size | | | | | | |
| < 1.0 cm | 23 (77%) | | 7 (88%) | 13 (93%) | 1 (33%) | 2 (100%) |
| ≥ 1.0 cm (large) | 3 (10%) | | 0 | 1 (7%) | 2 (67%) | 0 |
| High-grade dysplasia | 1 (3%) | | 0 | 1 (7%) | 0 | 0 |
| Tubular | 12 (40%) | | 0 | 12 (86%) | 0 | 0 |
| Tubulovillous | 2 (7%) | | 0 | 2 (14%) | 0 | 0 |
| Advanced | 4 (13%) | | 0 | 2 (14%) | 2 (67%) | 0 |
| Microvesicular HP | 3 (10%) | | 3 (38%) | 0 | 0 | 0 |
| Goblet cell rich HP | 1 (3%) | | 1 (13%) | 0 | 0 | 0 |
| Normal colorectal mucosa biopsy, n (%) | | | | | | |
| Ascending colon | 18 (60%) | 2 (67%) | 5 (63%) | 9 (64%) | 1 (33%) | 1 (50%) |
| Descending colon | 16 (53%) | 1 (33%) | 7 (88%) | 7 (50%) | 1 (33%) | 0 |

[a] Large defined as ≥ 1 cm; Advanced defined as large, tubulovillous, villous, or high-grade dysplasia; Proximal defined as cecum, ascending colon, hepatic flexure, transverse colon; Distal defined as splenic flexure, descending colon, sigmoid colon, or rectum

Table S3

| Tested Pair, CytoTRACE | | MWU-statistic | P-value, α' = 0.0169 |
|---|---|---|---|
| ASC | SSC | **20442988.5** | **0.00E+00** |
| ASC | NM | **259958857.5** | **0.00E+00** |
| SSC | NM | **81294745.0** | **4.0136e-72** |

Table S4

| ID | Source | Term ID | Term Name | $p_{adj}$ (query_1) |
|---|---|---|---|---|
| 1 | GO:MF | GO:0005005 | transmembrane-ephrin receptor activity | $3.648 \times 10^{-2}$ |
| 2 | GO:BP | GO:0006959 | humoral immune response | $3.619 \times 10^{-4}$ |
| 3 | GO:BP | GO:0010273 | detoxification of copper ion | $5.805 \times 10^{-3}$ |
| 4 | GO:BP | GO:0019730 | antimicrobial humoral response | $1.930 \times 10^{-4}$ |
| 5 | GO:BP | GO:0044419 | interspecies interaction between organisms | $1.096 \times 10^{-7}$ |
| 6 | GO:BP | GO:0046688 | response to copper ion | $3.083 \times 10^{-2}$ |
| 7 | GO:BP | GO:0048584 | positive regulation of response to stimulus | $1.617 \times 10^{-2}$ |
| 8 | GO:BP | GO:0071280 | cellular response to copper ion | $3.206 \times 10^{-3}$ |
| 9 | GO:BP | GO:0061687 | detoxification of inorganic compound | $1.043 \times 10^{-2}$ |
| 10 | GO:BP | GO:0071294 | cellular response to zinc ion | $4.870 \times 10^{-2}$ |
| 11 | GO:BP | GO:0097501 | stress response to metal ion | $1.043 \times 10^{-2}$ |
| 12 | GO:BP | GO:1990169 | stress response to copper ion | $5.805 \times 10^{-3}$ |
| 13 | GO:CC | GO:0005925 | focal adhesion | $1.182 \times 10^{-5}$ |
| 14 | GO:CC | GO:0030055 | cell-substrate junction | $1.516 \times 10^{-5}$ |
| 15 | GO:CC | GO:0030054 | cell junction | $4.753 \times 10^{-4}$ |
| 16 | GO:CC | GO:0070161 | anchoring junction | $1.857 \times 10^{-3}$ |
| 17 | GO:CC | GO:1903561 | extracellular vesicle | $1.635 \times 10^{-2}$ |
| 18 | GO:CC | GO:0070701 | mucus layer | $1.228 \times 10^{-2}$ |
| 19 | GO:CC | GO:0070062 | extracellular exosome | $5.588 \times 10^{-3}$ |
| 20 | GO:CC | GO:0043230 | extracellular organelle | $1.720 \times 10^{-2}$ |
| 21 | REAC | REAC:R-HSA-8953897 | Cellular responses to external stimuli | $1.932 \times 10^{-6}$ |
| 22 | REAC | REAC:R-HSA-2262752 | Cellular responses to stress | $1.053 \times 10^{-4}$ |
| 23 | REAC | REAC:R-HSA-5663205 | Infectious disease | $8.518 \times 10^{-5}$ |
| 24 | REAC | REAC:R-HSA-5660526 | Response to metal ions | $1.059 \times 10^{-2}$ |
| 25 | REAC | REAC:R-HSA-5661231 | Metallothioneins bind metals | $3.599 \times 10^{-3}$ |
| 26 | GO:BP | GO:0007166 | cell surface receptor signaling pathway | $1.979 \times 10^{-3}$ |
| 27 | GO:CC | GO:0005615 | extracellular space | $3.039 \times 10^{-2}$ |
| 28 | GO:CC | GO:0005576 | extracellular region | $1.104 \times 10^{-2}$ |
| 29 | GO:CC | GO:0031256 | leading edge membrane | $3.686 \times 10^{-2}$ |
| 30 | GO:CC | GO:0071944 | cell periphery | $1.315 \times 10^{-2}$ |

| | |
|---|---|
| **version** | e101_eg48_p14_baf17f0 |
| **date** | 11/15/2020, 11:36:52 PM |
| **organism** | hsapiens |

#### Table S5

| ID | Source | Term ID | Term Name | $p_{adj}$ (query_1) |
|---|---|---|---|---|
| 1 | GO:MF | GO:0045296 | cadherin binding | $1.046\times10^{-12}$ |
| 2 | GO:MF | GO:0044548 | S100 protein binding | $4.072\times10^{-6}$ |
| 3 | GO:MF | GO:0045295 | gamma-catenin binding | $9.316\times10^{-5}$ |
| 4 | GO:BP | GO:0043312 | neutrophil degranulation | $3.076\times10^{-6}$ |
| 5 | GO:BP | GO:0002446 | neutrophil mediated immunity | $5.207\times10^{-6}$ |
| 6 | GO:BP | GO:0036230 | granulocyte activation | $6.977\times10^{-6}$ |
| 7 | GO:BP | GO:0046903 | secretion | $1.625\times10^{-5}$ |
| 8 | GO:BP | GO:0032879 | regulation of localization | $3.592\times10^{-3}$ |
| 9 | GO:BP | GO:0034332 | adherens junction organization | $3.906\times10^{-3}$ |
| 10 | GO:BP | GO:0098609 | cell-cell adhesion | $5.743\times10^{-3}$ |
| 11 | GO:BP | GO:0002252 | immune effector process | $2.816\times10^{-2}$ |
| 12 | GO:BP | GO:0034097 | response to cytokine | $2.717\times10^{-2}$ |
| 13 | GO:CC | GO:0070062 | extracellular exosome | $8.355\times10^{-22}$ |
| 14 | GO:CC | GO:0030141 | secretory granule | $8.258\times10^{-10}$ |
| 15 | GO:CC | GO:0031252 | cell leading edge | $1.529\times10^{-9}$ |
| 16 | GO:CC | GO:0070369 | beta-catenin-TCF7L2 complex | $2.930\times10^{-2}$ |
| 17 | KEGG | KEGG:05100 | Bacterial invasion of epithelial cells | $4.691\times10^{-2}$ |
| 18 | KEGG | KEGG:04520 | Adherens junction | $9.901\times10^{-6}$ |
| 19 | KEGG | KEGG:00190 | Oxidative phosphorylation | $6.673\times10^{-8}$ |
| 20 | REAC | REAC:R-HSA-67... | Neutrophil degranulation | $3.298\times10^{-3}$ |
| 21 | REAC | REAC:R-HSA-44... | VEGFA-VEGFR2 Pathway | $1.469\times10^{-2}$ |
| 22 | REAC | REAC:R-HSA-39... | EPHB-mediated forward signaling | $2.369\times10^{-2}$ |
| 23 | REAC | REAC:R-HSA-19... | Signaling by VEGF | $3.261\times10^{-2}$ |
| 24 | REAC | REAC:R-HSA-56... | RHO GTPases activate IQGAPs | $9.474\times10^{-6}$ |
| 25 | GO:BP | GO:0045321 | leukocyte activation | $2.430\times10^{-3}$ |
| 26 | GO:BP | GO:0030029 | actin filament-based process | $1.977\times10^{-4}$ |
| 27 | GO:CC | GO:0016342 | catenin complex | $2.449\times10^{-4}$ |
| 28 | REAC | REAC:R-HSA-41... | Adherens junctions interactions | $2.384\times10^{-4}$ |
| 29 | REAC | REAC:R-HSA-56... | RHO GTPases activate KTN1 | $6.561\times10^{-3}$ |
| 30 | WP | WP:WP623 | Oxidative phosphorylation | $9.265\times10^{-6}$ |

| | |
|---|---|
| **version** | e101_eg48_p14_baf17f0 |
| **date** | 11/23/2020, 1:22:17 PM |
| **organism** | hsapiens |

### Table S6

| ID | Source | Term ID | Term Name | $p_{adj}$ (query_1) |
|---|---|---|---|---|
| 1 | GO:MF | GO:0003735 | structural constituent of ribosome | $3.389\times10^{-82}$ |
| 2 | REAC | REAC:R-HSA-21... | TGF-beta receptor signaling in EMT (epithelial to m... | $7.603\times10^{-3}$ |
| 3 | REAC | REAC:R-HSA-12... | Downregulation of ERBB4 signaling | $2.922\times10^{-2}$ |
| 4 | REAC | REAC:R-HSA-89... | Regulation of PTEN localization | $4.345\times10^{-2}$ |
| 5 | REAC | REAC:R-HSA-88... | PTK6 Regulates RTKs and Their Effectors AKT1 and ... | $4.345\times10^{-2}$ |
| 6 | REAC | REAC:R-HSA-96... | Response of EIF2AK4 (GCN2) to amino acid deficie... | $1.105\times10^{-83}$ |
| 7 | REAC | REAC:R-HSA-90... | Regulation of expression of SLITs and ROBOs | $5.225\times10^{-69}$ |
| 8 | GO:CC | GO:0101002 | ficolin-1-rich granule | $1.267\times10^{-3}$ |
| 9 | GO:CC | GO:0042588 | zymogen granule | $1.813\times10^{-3}$ |
| 10 | GO:CC | GO:0030141 | secretory granule | $3.357\times10^{-3}$ |
| 11 | GO:CC | GO:0070062 | extracellular exosome | $1.871\times10^{-28}$ |
| 12 | GO:BP | GO:0042119 | neutrophil activation | $2.562\times10^{-3}$ |
| 13 | GO:BP | GO:0036230 | granulocyte activation | $3.057\times10^{-3}$ |
| 14 | GO:MF | GO:0045296 | cadherin binding | $5.484\times10^{-9}$ |
| 15 | GO:MF | GO:0050839 | cell adhesion molecule binding | $2.545\times10^{-8}$ |
| 16 | GO:BP | GO:0051649 | establishment of localization in cell | $3.288\times10^{-30}$ |
| 17 | GO:CC | GO:0030054 | cell junction | $9.813\times10^{-13}$ |
| 18 | GO:CC | GO:0099503 | secretory vesicle | $9.399\times10^{-3}$ |
| 19 | REAC | REAC:R-HSA-22... | Cellular responses to stress | $7.961\times10^{-40}$ |
| 20 | REAC | REAC:R-HSA-96... | Modulation by Mtb of host immune system | $1.075\times10^{-3}$ |

| | |
|---|---|
| **version** | e101_eg48_p14_baf17f0 |
| **date** | 11/23/2020, 1:26:14 PM |
| **organism** | hsapiens |

Table S7

| ID | Source | Term ID | Term Name | $p_{adj}$ (query_1) |
|---|---|---|---|---|
| 1 | GO:MF | GO:0008270 | zinc ion binding | $4.218\times10^{-2}$ |
| 2 | GO:MF | GO:0015103 | inorganic anion transmembrane transporter activity | $3.941\times10^{-2}$ |
| 3 | GO:BP | GO:0010273 | detoxification of copper ion | $4.828\times10^{-9}$ |
| 4 | GO:BP | GO:1990169 | stress response to copper ion | $4.828\times10^{-9}$ |
| 5 | GO:BP | GO:0061687 | detoxification of inorganic compound | $1.281\times10^{-8}$ |
| 6 | GO:BP | GO:0097501 | stress response to metal ion | $1.281\times10^{-8}$ |
| 7 | GO:BP | GO:0071294 | cellular response to zinc ion | $1.587\times10^{-7}$ |
| 8 | GO:BP | GO:0071280 | cellular response to copper ion | $4.608\times10^{-7}$ |
| 9 | GO:BP | GO:0071276 | cellular response to cadmium ion | $3.530\times10^{-6}$ |
| 10 | GO:BP | GO:0006882 | cellular zinc ion homeostasis | $3.530\times10^{-6}$ |
| 11 | GO:BP | GO:0055069 | zinc ion homeostasis | $4.930\times10^{-6}$ |
| 12 | GO:BP | GO:0046688 | response to copper ion | $7.868\times10^{-6}$ |
| 13 | GO:BP | GO:0010043 | response to zinc ion | $4.210\times10^{-5}$ |
| 14 | KEGG | KEGG:04978 | Mineral absorption | $2.441\times10^{-4}$ |
| 15 | REAC | REAC:R-HSA-56... | Metallothioneins bind metals | $3.726\times10^{-9}$ |
| 16 | WP | WP:WP3529 | Zinc homeostasis | $4.180\times10^{-5}$ |
| 17 | WP | WP:WP3286 | Copper homeostasis | $7.216\times10^{-3}$ |
| 18 | GO:CC | GO:0070701 | mucus layer | $3.471\times10^{-3}$ |
| 19 | GO:CC | GO:0031225 | anchored component of membrane | $1.937\times10^{-7}$ |
| 20 | GO:BP | GO:0046686 | response to cadmium ion | $1.170\times10^{-4}$ |

| | |
|---|---|
| **version** | e101_eg48_p14_baf17f0 |
| **date** | 11/23/2020, 1:32:22 PM |
| **organism** | hsapiens |

Table S8

| ID | Source | Term ID | Term Name | $p_{adj}$ (query_1) |
|---|---|---|---|---|
| 1 | GO:MF | GO:0046332 | SMAD binding | $1.271\times10^{-3}$ |
| 2 | GO:MF | GO:0050839 | cell adhesion molecule binding | $9.284\times10^{-8}$ |
| 3 | GO:MF | GO:0045296 | cadherin binding | $2.512\times10^{-6}$ |
| 4 | GO:MF | GO:0005003 | ephrin receptor activity | $3.868\times10^{-2}$ |
| 5 | GO:MF | GO:0005540 | hyaluronic acid binding | $3.072\times10^{-2}$ |
| 6 | GO:BP | GO:0051674 | localization of cell | $1.231\times10^{-5}$ |
| 7 | GO:BP | GO:0038018 | Wnt receptor catabolic process | $4.018\times10^{-3}$ |
| 8 | GO:BP | GO:0007166 | cell surface receptor signaling pathway | $3.954\times10^{-3}$ |
| 9 | GO:BP | GO:0060828 | regulation of canonical Wnt signaling pathway | $1.963\times10^{-4}$ |
| 10 | GO:BP | GO:0071560 | cellular response to transforming growth factor be... | $4.085\times10^{-5}$ |
| 11 | GO:CC | GO:0070062 | extracellular exosome | $5.631\times10^{-11}$ |
| 12 | GO:CC | GO:0071148 | TEAD-1-YAP complex | $2.725\times10^{-2}$ |
| 13 | REAC | REAC:R-HSA-39... | EPHB-mediated forward signaling | $5.214\times10^{-3}$ |
| 14 | REAC | REAC:R-HSA-46... | Regulation of FZD by ubiquitination | $9.883\times10^{-3}$ |
| 15 | REAC | REAC:R-HSA-90... | Regulation of expression of SLITs and ROBOs | $4.493\times10^{-15}$ |
| 16 | REAC | REAC:R-HSA-96... | Response of EIF2AK4 (GCN2) to amino acid deficie... | $4.060\times10^{-21}$ |
| 17 | KEGG | KEGG:04310 | Wnt signaling pathway | $1.708\times10^{-2}$ |
| 18 | GO:BP | GO:0060389 | pathway-restricted SMAD protein phosphorylation | $3.751\times10^{-3}$ |
| 19 | GO:BP | GO:0007179 | transforming growth factor beta receptor signalin... | $1.095\times10^{-4}$ |
| 20 | GO:BP | GO:0030111 | regulation of Wnt signaling pathway | $4.781\times10^{-6}$ |

| | |
|---|---|
| **version** | e101_eg48_p14_baf17f0 |
| **date** | 11/23/2020, 1:45:38 PM |
| **organism** | hsapiens |

Table S9

| ID | Source | Term ID | Term Name | $p_{adj}$ (query_1) |
|---|---|---|---|---|
| 1 | GO:MF | GO:0045296 | cadherin binding | $2.509\times10^{-14}$ |
| 2 | GO:MF | GO:0044548 | S100 protein binding | $6.279\times10^{-8}$ |
| 3 | GO:MF | GO:0003779 | actin binding | $2.821\times10^{-7}$ |
| 4 | GO:MF | GO:0008092 | cytoskeletal protein binding | $1.006\times10^{-4}$ |
| 5 | GO:MF | GO:0048306 | calcium-dependent protein binding | $3.122\times10^{-4}$ |
| 6 | GO:BP | GO:0043312 | neutrophil degranulation | $9.897\times10^{-21}$ |
| 7 | GO:BP | GO:0002283 | neutrophil activation involved in immune response | $1.235\times10^{-20}$ |
| 8 | GO:BP | GO:0036230 | granulocyte activation | $4.856\times10^{-20}$ |
| 9 | GO:BP | GO:0043299 | leukocyte degranulation | $4.008\times10^{-19}$ |
| 10 | GO:BP | GO:0002275 | myeloid cell activation involved in immune respon... | $8.194\times10^{-19}$ |
| 11 | GO:BP | GO:0002444 | myeloid leukocyte mediated immunity | $1.281\times10^{-18}$ |
| 12 | GO:BP | GO:0002252 | immune effector process | $1.111\times10^{-11}$ |
| 13 | GO:BP | GO:0002443 | leukocyte mediated immunity | $2.835\times10^{-13}$ |
| 14 | GO:BP | GO:0042060 | wound healing | $3.751\times10^{-2}$ |
| 15 | GO:BP | GO:0071345 | cellular response to cytokine stimulus | $2.707\times10^{-2}$ |
| 16 | GO:BP | GO:0009611 | response to wounding | $1.995\times10^{-3}$ |
| 17 | GO:CC | GO:0070062 | extracellular exosome | $7.626\times10^{-34}$ |
| 18 | GO:CC | GO:0042582 | azurophil granule | $4.746\times10^{-8}$ |
| 19 | GO:CC | GO:0000323 | lytic vacuole | $6.410\times10^{-7}$ |
| 20 | GO:CC | GO:0030141 | secretory granule | $9.840\times10^{-20}$ |
| 21 | GO:CC | GO:0005903 | brush border | $4.539\times10^{-6}$ |
| 22 | GO:CC | GO:0101002 | ficolin-1-rich granule | $6.236\times10^{-4}$ |
| 23 | GO:CC | GO:0031528 | microvillus membrane | $1.219\times10^{-3}$ |
| 24 | GO:CC | GO:0045177 | apical part of cell | $3.088\times10^{-3}$ |
| 25 | KEGG | KEGG:05130 | Pathogenic Escherichia coli infection | $3.262\times10^{-2}$ |
| 26 | KEGG | KEGG:04810 | Regulation of actin cytoskeleton | $1.705\times10^{-2}$ |
| 27 | REAC | REAC:R-HSA-16... | Innate Immune System | $3.285\times10^{-9}$ |
| 28 | REAC | REAC:R-HSA-67... | Neutrophil degranulation | $4.980\times10^{-14}$ |
| 29 | REAC | REAC:R-HSA-11... | Platelet degranulation | $5.915\times10^{-3}$ |
| 30 | WP | WP:WP2272 | Pathogenic Escherichia coli infection | $2.303\times10^{-2}$ |

| | |
|---|---|
| **version** | e101_eg48_p14_baf17f0 |
| **date** | 11/23/2020, 1:50:45 PM |
| **organism** | hsapiens |

Table S10

| ID | Source | Term ID | Term Name | $p_{adj}$ (query_1) |
|---|---|---|---|---|
| 1 | GO:MF | GO:0031730 | CCR5 chemokine receptor binding | $3.762\times10^{-2}$ |
| 2 | GO:MF | GO:0070888 | E-box binding | $6.497\times10^{-4}$ |
| 3 | GO:CC | GO:0000790 | nuclear chromatin | $1.193\times10^{-42}$ |
| 4 | KEGG | KEGG:05210 | Colorectal cancer | $1.091\times10^{-2}$ |
| 5 | KEGG | KEGG:04310 | Wnt signaling pathway | $9.918\times10^{-3}$ |
| 6 | KEGG | KEGG:05202 | Transcriptional misregulation in cancer | $9.035\times10^{-3}$ |
| 7 | KEGG | KEGG:04659 | Th17 cell differentiation | $2.132\times10^{-2}$ |
| 8 | KEGG | KEGG:05321 | Inflammatory bowel disease | $2.936\times10^{-2}$ |
| 9 | REAC | REAC:R-HSA-44... | Binding of TCF/LEF:CTNNB1 to target gene promo... | $6.595\times10^{-4}$ |
| 10 | REAC | REAC:R-HSA-89... | RUNX3 regulates WNT signaling | $6.595\times10^{-4}$ |
| 11 | REAC | REAC:R-HSA-89... | RUNX3 regulates YAP1-mediated transcription | $6.595\times10^{-4}$ |
| 12 | REAC | REAC:R-HSA-15... | Signaling by NOTCH | $4.827\times10^{-3}$ |
| 13 | REAC | REAC:R-HSA-67... | Interleukin-4 and Interleukin-13 signaling | $3.610\times10^{-3}$ |
| 14 | REAC | REAC:R-HSA-68... | TP53 Regulates Transcription of Genes Involved in ... | $5.360\times10^{-3}$ |
| 15 | REAC | REAC:R-HSA-21... | SMAD2/SMAD3:SMAD4 heterotrimer regulates tra... | $1.475\times10^{-2}$ |
| 16 | REAC | REAC:R-HSA-15... | G0 and Early G1 | $1.569\times10^{-2}$ |
| 17 | REAC | REAC:R-HSA-20... | TCF dependent signaling in response to WNT | $4.105\times10^{-2}$ |
| 18 | REAC | REAC:R-HSA-13... | Activation of PUMA and translocation to mitocho... | $4.478\times10^{-2}$ |
| 19 | REAC | REAC:R-HSA-88... | Transcriptional regulation by RUNX3 | $8.596\times10^{-6}$ |
| 20 | WP | WP:WP127 | IL-5 Signaling Pathway | $2.405\times10^{-3}$ |
| 21 | WP | WP:WP3972 | PDGFR-beta pathway | $2.824\times10^{-3}$ |
| 22 | WP | WP:WP399 | Wnt Signaling Pathway and Pluripotency | $2.869\times10^{-3}$ |
| 23 | WP | WP:WP710 | DNA Damage Response (only ATM dependent) | $4.079\times10^{-3}$ |
| 24 | WP | WP:WP49 | IL-2 Signaling Pathway | $2.928\times10^{-3}$ |
| 25 | WP | WP:WP205 | IL-7 Signaling Pathway | $1.630\times10^{-2}$ |
| 26 | WP | WP:WP4216 | Chromosomal and microsatellite instability in colo... | $1.189\times10^{-2}$ |
| 27 | WP | WP:WP560 | TGF-beta Receptor Signaling | $1.888\times10^{-2}$ |
| 28 | WP | WP:WP61 | Notch Signaling Pathway Netpath | $3.893\times10^{-2}$ |
| 29 | GO:BP | GO:0009790 | embryo development | $2.099\times10^{-14}$ |
| 30 | GO:BP | GO:0008283 | cell population proliferation | $3.332\times10^{-8}$ |

| | |
|---|---|
| **version** | e101_eg48_p14_baf17f0 |
| **date** | 11/23/2020, 2:01:23 PM |
| **organism** | hsapiens |

Table S11

| ID | Source | Term ID | Term Name | $p_{adj}$ (query_1) |
|---|---|---|---|---|
| 1 | GO:MF | GO:0046332 | SMAD binding | $3.575 \times 10^{-4}$ |
| 2 | GO:MF | GO:0070412 | R-SMAD binding | $8.051 \times 10^{-3}$ |
| 3 | GO:MF | GO:0070644 | vitamin D response element binding | $7.375 \times 10^{-3}$ |
| 4 | GO:MF | GO:0035497 | cAMP response element binding | $9.436 \times 10^{-6}$ |
| 5 | GO:BP | GO:0033554 | cellular response to stress | $5.603 \times 10^{-7}$ |
| 6 | GO:BP | GO:0002521 | leukocyte differentiation | $8.855 \times 10^{-10}$ |
| 7 | GO:BP | GO:0030099 | myeloid cell differentiation | $7.764 \times 10^{-10}$ |
| 8 | GO:BP | GO:0001890 | placenta development | $3.422 \times 10^{-7}$ |
| 9 | GO:CC | GO:0035976 | transcription factor AP-1 complex | $4.981 \times 10^{-9}$ |
| 10 | KEGG | KEGG:04668 | TNF signaling pathway | $1.405 \times 10^{-8}$ |
| 11 | KEGG | KEGG:04659 | Th17 cell differentiation | $5.695 \times 10^{-5}$ |
| 12 | KEGG | KEGG:04657 | IL-17 signaling pathway | $7.724 \times 10^{-3}$ |
| 13 | KEGG | KEGG:04310 | Wnt signaling pathway | $1.038 \times 10^{-2}$ |
| 14 | KEGG | KEGG:05321 | Inflammatory bowel disease | $1.876 \times 10^{-2}$ |
| 15 | KEGG | KEGG:04152 | AMPK signaling pathway | $1.766 \times 10^{-2}$ |
| 16 | KEGG | KEGG:04010 | MAPK signaling pathway | $4.583 \times 10^{-2}$ |
| 17 | KEGG | KEGG:04022 | cGMP-PKG signaling pathway | $4.617 \times 10^{-2}$ |
| 18 | REAC | REAC:R-HSA-88... | CREB3 factors activate genes | $1.248 \times 10^{-5}$ |
| 19 | REAC | REAC:R-HSA-38... | ATF4 activates genes in response to endoplasmic r... | $2.113 \times 10^{-3}$ |
| 20 | REAC | REAC:R-HSA-21... | SMAD2/SMAD3:SMAD4 heterotrimer regulates tra... | $3.905 \times 10^{-3}$ |
| 21 | REAC | REAC:R-HSA-38... | PERK regulates gene expression | $4.258 \times 10^{-3}$ |
| 22 | REAC | REAC:R-HSA-17... | Signaling by TGF-beta Receptor Complex | $4.353 \times 10^{-2}$ |
| 23 | REAC | REAC:R-HSA-21... | Transcriptional activity of SMAD2/SMAD3:SMAD4 ... | $1.032 \times 10^{-2}$ |
| 24 | WP | WP:WP366 | TGF-beta Signaling Pathway | $2.180 \times 10^{-4}$ |
| 25 | WP | WP:WP364 | IL-6 signaling pathway | $8.602 \times 10^{-4}$ |
| 26 | WP | WP:WP4754 | IL-18 signaling pathway | $3.309 \times 10^{-2}$ |
| 27 | WP | WP:WP615 | Senescence and Autophagy in Cancer | $3.499 \times 10^{-2}$ |
| 28 | WP | WP:WP382 | MAPK Signaling Pathway | $3.913 \times 10^{-2}$ |
| 29 | WP | WP:WP4342 | Vitamins A and D - action mechanisms | $1.945 \times 10^{-5}$ |
| 30 | WP | WP:WP4482 | Vitamin D in inflammatory diseases | $7.067 \times 10^{-4}$ |

| | |
|---|---|
| **version** | e101_eg48_p14_baf17f0 |
| **date** | 11/23/2020, 1:54:22 PM |
| **organism** | hsapiens |

#### Table S12

| ID | Source | Term ID | Term Name | $p_{adj}$ (query_1) |
|---|---|---|---|---|
| 1 | GO:MF | GO:0008270 | zinc ion binding | $4.218\times10^{-2}$ |
| 2 | GO:MF | GO:0015103 | inorganic anion transmembrane transporter activity | $3.941\times10^{-2}$ |
| 3 | GO:BP | GO:0010273 | detoxification of copper ion | $4.828\times10^{-9}$ |
| 4 | GO:BP | GO:1990169 | stress response to copper ion | $4.828\times10^{-9}$ |
| 5 | GO:BP | GO:0061687 | detoxification of inorganic compound | $1.281\times10^{-8}$ |
| 6 | GO:BP | GO:0097501 | stress response to metal ion | $1.281\times10^{-8}$ |
| 7 | GO:BP | GO:0071294 | cellular response to zinc ion | $1.587\times10^{-7}$ |
| 8 | GO:BP | GO:0071280 | cellular response to copper ion | $4.608\times10^{-7}$ |
| 9 | GO:BP | GO:0071276 | cellular response to cadmium ion | $3.530\times10^{-6}$ |
| 10 | GO:BP | GO:0006882 | cellular zinc ion homeostasis | $3.530\times10^{-6}$ |
| 11 | GO:BP | GO:0055069 | zinc ion homeostasis | $4.930\times10^{-6}$ |
| 12 | GO:BP | GO:0046688 | response to copper ion | $7.868\times10^{-6}$ |
| 13 | GO:BP | GO:0010043 | response to zinc ion | $4.210\times10^{-5}$ |
| 14 | KEGG | KEGG:04978 | Mineral absorption | $2.441\times10^{-4}$ |
| 15 | REAC | REAC:R-HSA-56... | Metallothioneins bind metals | $3.726\times10^{-9}$ |
| 16 | WP | WP:WP3529 | Zinc homeostasis | $4.180\times10^{-5}$ |
| 17 | WP | WP:WP3286 | Copper homeostasis | $7.216\times10^{-3}$ |
| 18 | GO:CC | GO:0070701 | mucus layer | $3.471\times10^{-3}$ |
| 19 | GO:CC | GO:0031225 | anchored component of membrane | $1.937\times10^{-7}$ |
| 20 | GO:BP | GO:0046686 | response to cadmium ion | $1.170\times10^{-4}$ |

| | |
|---|---|
| **version** | e101_eg48_p14_baf17f0 |
| **date** | 11/23/2020, 1:32:22 PM |
| **organism** | hsapiens |

Table S13

| ID | Source | Term ID | Term Name | $p_{adj}$ (query_1) |
|---|---|---|---|---|
| 1 | GO:MF | GO:0003735 | structural constituent of ribosome | $1.536\times10^{-130}$ |
| 2 | GO:MF | GO:0017134 | fibroblast growth factor binding | $1.105\times10^{-2}$ |
| 3 | GO:MF | GO:0031625 | ubiquitin protein ligase binding | $1.027\times10^{-2}$ |
| 4 | GO:MF | GO:0023026 | MHC class II protein complex binding | $9.175\times10^{-3}$ |
| 5 | WP | WP:WP3888 | VEGFA-VEGFR2 Signaling Pathway | $3.935\times10^{-3}$ |
| 6 | REAC | REAC:R-HSA-90... | Regulation of expression of SLITs and ROBOs | $1.191\times10^{-112}$ |
| 7 | REAC | REAC:R-HSA-56... | Infectious disease | $1.506\times10^{-63}$ |
| 8 | REAC | REAC:R-HSA-22... | Cellular responses to stress | $3.114\times10^{-70}$ |
| 9 | GO:CC | GO:0070062 | extracellular exosome | $1.178\times10^{-33}$ |
| 10 | GO:CC | GO:0005925 | focal adhesion | $3.541\times10^{-40}$ |
| 11 | GO:CC | GO:0030054 | cell junction | $3.069\times10^{-17}$ |
| 12 | GO:CC | GO:0101002 | ficolin-1-rich granule | $9.651\times10^{-4}$ |
| 13 | GO:BP | GO:0044403 | symbiotic process | $1.705\times10^{-74}$ |
| 14 | GO:BP | GO:0045047 | protein targeting to ER | $1.140\times10^{-148}$ |
| 15 | GO:BP | GO:0072594 | establishment of protein localization to organelle | $9.442\times10^{-87}$ |
| 16 | GO:MF | GO:0045296 | cadherin binding | $1.041\times10^{-8}$ |
| 17 | GO:MF | GO:0050839 | cell adhesion molecule binding | $3.184\times10^{-8}$ |
| 18 | GO:BP | GO:0044419 | interspecies interaction between organisms | $6.668\times10^{-49}$ |
| 19 | GO:BP | GO:1901575 | organic substance catabolic process | $7.840\times10^{-49}$ |
| 20 | GO:BP | GO:0071705 | nitrogen compound transport | $2.078\times10^{-47}$ |

| | |
|---|---|
| **version** | e101_eg48_p14_baf17f0 |
| **date** | 11/23/2020, 1:33:42 PM |
| **organism** | hsapiens |

Table S14

| ID | Source | Term ID | Term Name | $p_{adj}$ (query_1) |
|---|---|---|---|---|
| 1 | GO:MF | GO:0045296 | cadherin binding | $4.817 \times 10^{-11}$ |
| 2 | GO:MF | GO:0044548 | S100 protein binding | $1.032 \times 10^{-6}$ |
| 3 | GO:BP | GO:0002446 | neutrophil mediated immunity | $2.195 \times 10^{-17}$ |
| 4 | GO:BP | GO:0043312 | neutrophil degranulation | $8.544 \times 10^{-17}$ |
| 5 | GO:BP | GO:0042119 | neutrophil activation | $2.375 \times 10^{-16}$ |
| 6 | GO:BP | GO:0045055 | regulated exocytosis | $2.958 \times 10^{-16}$ |
| 7 | GO:BP | GO:0036230 | granulocyte activation | $3.451 \times 10^{-16}$ |
| 8 | GO:BP | GO:0002252 | immune effector process | $1.141 \times 10^{-14}$ |
| 9 | GO:BP | GO:0002443 | leukocyte mediated immunity | $2.103 \times 10^{-13}$ |
| 10 | GO:BP | GO:0002274 | myeloid leukocyte activation | $1.660 \times 10^{-12}$ |
| 11 | GO:BP | GO:0006915 | apoptotic process | $7.093 \times 10^{-6}$ |
| 12 | GO:BP | GO:0038093 | Fc receptor signaling pathway | $1.475 \times 10^{-4}$ |
| 13 | GO:BP | GO:0071345 | cellular response to cytokine stimulus | $2.329 \times 10^{-4}$ |
| 14 | GO:BP | GO:0001666 | response to hypoxia | $1.004 \times 10^{-3}$ |
| 15 | GO:CC | GO:0070062 | extracellular exosome | $9.947 \times 10^{-51}$ |
| 16 | GO:CC | GO:0101002 | ficolin-1-rich granule | $5.616 \times 10^{-7}$ |
| 17 | KEGG | KEGG:05100 | Bacterial invasion of epithelial cells | $3.618 \times 10^{-7}$ |
| 18 | KEGG | KEGG:04810 | Regulation of actin cytoskeleton | $1.859 \times 10^{-5}$ |
| 19 | KEGG | KEGG:05130 | Pathogenic Escherichia coli infection | $3.388 \times 10^{-5}$ |
| 20 | KEGG | KEGG:04670 | Leukocyte transendothelial migration | $3.342 \times 10^{-4}$ |
| 21 | REAC | REAC:R-HSA-16... | Innate Immune System | $1.554 \times 10^{-15}$ |
| 22 | REAC | REAC:R-HSA-67... | Neutrophil degranulation | $1.072 \times 10^{-14}$ |
| 23 | REAC | REAC:R-HSA-26... | EPH-Ephrin signaling | $1.500 \times 10^{-9}$ |
| 24 | REAC | REAC:R-HSA-39... | EPHB-mediated forward signaling | $1.368 \times 10^{-8}$ |
| 25 | REAC | REAC:R-HSA-19... | RHO GTPase Effectors | $1.738 \times 10^{-8}$ |
| 26 | WP | WP:WP3888 | VEGFA-VEGFR2 Signaling Pathway | $4.637 \times 10^{-4}$ |
| 27 | WP | WP:WP2272 | Pathogenic Escherichia coli infection | $3.373 \times 10^{-8}$ |
| 28 | REAC | REAC:R-HSA-56... | MAPK6/MAPK4 signaling | $4.107 \times 10^{-2}$ |
| 29 | REAC | REAC:R-HSA-19... | Signaling by Rho GTPases | $2.115 \times 10^{-6}$ |
| 30 | REAC | REAC:R-HSA-44... | Interleukin-12 family signaling | $2.527 \times 10^{-2}$ |

| | |
|---|---|
| **version** | e101_eg48_p14_baf17f0 |
| **date** | 11/23/2020, 1:40:31 PM |
| **organism** | hsapiens |

Table S15

| ID | WD 79442 | WD 69814 | WD 69753 |
|---|---|---|---|
| Age | 40 | 54 | 69 |
| Sex | M | F | M |
| Ethnicity | B | W | W |
| Patient Height | 177.8 cm | 154.9 cm | 185.4 cm |
| Patient Weight | 86.4 kg | 45.5kg | 83.5 kg |
| Patient BMI | 27.3 | 19 | 24.3 |
| Tumor location | Hepatic flexure | Sigmoid colon | Cecum |
| Tumor greatest dimension | 5.8cm | 3.6cm | 6.2cm |
| Grade | G2 Moderately differentiated | G2 Moderately differentiated | G3 Poorly differentiated |
| Staging | I | IIIB | IIIB |
| Tumor extension | Tumor invades muscularis propria | Tumor invades through the muscularis propria into pericolorectal tissue | Tumor invades through the muscularis propria into pericolorectal tissue |
| Metastasis | None | Lymph nodes | Lymph nodes |
| Microsatellite status | MSS | MSS | MSI-H |

Table S16

| Tested Pair, CytoTRACE | | MWU-statistic | P-value, $\alpha' = 0.0085$ |
|---|---|---|---|
| MSS | ASC | 4112859.5 | 8.19E-02 |
| MSI | SSC | **2519966** | **4.45E-278** |
| MSI | ASC | **4137738** | **4.26E-145** |
| SSC | ASC | **6454505** | **9.47E-199** |
| SSC | MSS | **1394002** | **2.24E-68** |
| MSI | MSS | **806408** | **9.32E-79** |

Table S17

| ID | Source | Term ID | Term Name | $p_{adj}$ (query_1) |
|---|---|---|---|---|
| 1 | GO:MF | GO:0031730 | CCR5 chemokine receptor binding | $3.762\times10^{-2}$ |
| 2 | GO:MF | GO:0070888 | E-box binding | $6.497\times10^{-4}$ |
| 3 | GO:CC | GO:0000790 | nuclear chromatin | $1.193\times10^{-42}$ |
| 4 | KEGG | KEGG:05210 | Colorectal cancer | $1.091\times10^{-2}$ |
| 5 | KEGG | KEGG:04310 | Wnt signaling pathway | $9.918\times10^{-3}$ |
| 6 | KEGG | KEGG:05202 | Transcriptional misregulation in cancer | $9.035\times10^{-3}$ |
| 7 | KEGG | KEGG:04659 | Th17 cell differentiation | $2.132\times10^{-2}$ |
| 8 | KEGG | KEGG:05321 | Inflammatory bowel disease | $2.936\times10^{-2}$ |
| 9 | REAC | REAC:R-HSA-44... | Binding of TCF/LEF:CTNNB1 to target gene promo... | $6.595\times10^{-4}$ |
| 10 | REAC | REAC:R-HSA-89... | RUNX3 regulates WNT signaling | $6.595\times10^{-4}$ |
| 11 | REAC | REAC:R-HSA-89... | RUNX3 regulates YAP1-mediated transcription | $6.595\times10^{-4}$ |
| 12 | REAC | REAC:R-HSA-15... | Signaling by NOTCH | $4.827\times10^{-3}$ |
| 13 | REAC | REAC:R-HSA-67... | Interleukin-4 and Interleukin-13 signaling | $3.610\times10^{-3}$ |
| 14 | REAC | REAC:R-HSA-68... | TP53 Regulates Transcription of Genes Involved in ... | $5.360\times10^{-3}$ |
| 15 | REAC | REAC:R-HSA-21... | SMAD2/SMAD3:SMAD4 heterotrimer regulates tra... | $1.475\times10^{-2}$ |
| 16 | REAC | REAC:R-HSA-15... | G0 and Early G1 | $1.569\times10^{-2}$ |
| 17 | REAC | REAC:R-HSA-20... | TCF dependent signaling in response to WNT | $4.105\times10^{-2}$ |
| 18 | REAC | REAC:R-HSA-13... | Activation of PUMA and translocation to mitocho... | $4.478\times10^{-2}$ |
| 19 | REAC | REAC:R-HSA-88... | Transcriptional regulation by RUNX3 | $8.596\times10^{-6}$ |
| 20 | WP | WP:WP127 | IL-5 Signaling Pathway | $2.405\times10^{-3}$ |
| 21 | WP | WP:WP3972 | PDGFR-beta pathway | $2.824\times10^{-3}$ |
| 22 | WP | WP:WP399 | Wnt Signaling Pathway and Pluripotency | $2.869\times10^{-3}$ |
| 23 | WP | WP:WP710 | DNA Damage Response (only ATM dependent) | $4.079\times10^{-3}$ |
| 24 | WP | WP:WP49 | IL-2 Signaling Pathway | $2.928\times10^{-3}$ |
| 25 | WP | WP:WP205 | IL-7 Signaling Pathway | $1.630\times10^{-2}$ |
| 26 | WP | WP:WP4216 | Chromosomal and microsatellite instability in colo... | $1.189\times10^{-2}$ |
| 27 | WP | WP:WP560 | TGF-beta Receptor Signaling | $1.888\times10^{-2}$ |
| 28 | WP | WP:WP61 | Notch Signaling Pathway Netpath | $3.893\times10^{-2}$ |
| 29 | GO:BP | GO:0009790 | embryo development | $2.099\times10^{-14}$ |
| 30 | GO:BP | GO:0008283 | cell population proliferation | $3.332\times10^{-8}$ |

| | |
|---|---|
| **version** | e101_eg48_p14_baf17f0 |
| **date** | 11/23/2020, 2:01:23 PM |
| **organism** | hsapiens |

Table S18

| ID | Source | Term ID | Term Name | $p_{adj}$ (query_1) |
|---|---|---|---|---|
| 1 | GO:MF | GO:0046332 | SMAD binding | $3.575\times10^{-4}$ |
| 2 | GO:MF | GO:0070412 | R-SMAD binding | $8.051\times10^{-3}$ |
| 3 | GO:MF | GO:0070644 | vitamin D response element binding | $7.375\times10^{-3}$ |
| 4 | GO:MF | GO:0035497 | cAMP response element binding | $9.436\times10^{-6}$ |
| 5 | GO:BP | GO:0033554 | cellular response to stress | $5.603\times10^{-7}$ |
| 6 | GO:BP | GO:0002521 | leukocyte differentiation | $8.855\times10^{-10}$ |
| 7 | GO:BP | GO:0030099 | myeloid cell differentiation | $7.764\times10^{-10}$ |
| 8 | GO:BP | GO:0001890 | placenta development | $3.422\times10^{-7}$ |
| 9 | GO:CC | GO:0035976 | transcription factor AP-1 complex | $4.981\times10^{-9}$ |
| 10 | KEGG | KEGG:04668 | TNF signaling pathway | $1.405\times10^{-8}$ |
| 11 | KEGG | KEGG:04659 | Th17 cell differentiation | $5.695\times10^{-5}$ |
| 12 | KEGG | KEGG:04657 | IL-17 signaling pathway | $7.724\times10^{-3}$ |
| 13 | KEGG | KEGG:04310 | Wnt signaling pathway | $1.038\times10^{-2}$ |
| 14 | KEGG | KEGG:05321 | Inflammatory bowel disease | $1.876\times10^{-2}$ |
| 15 | KEGG | KEGG:04152 | AMPK signaling pathway | $1.766\times10^{-2}$ |
| 16 | KEGG | KEGG:04010 | MAPK signaling pathway | $4.583\times10^{-2}$ |
| 17 | KEGG | KEGG:04022 | cGMP-PKG signaling pathway | $4.617\times10^{-2}$ |
| 18 | REAC | REAC:R-HSA-88... | CREB3 factors activate genes | $1.248\times10^{-5}$ |
| 19 | REAC | REAC:R-HSA-38... | ATF4 activates genes in response to endoplasmic r... | $2.113\times10^{-3}$ |
| 20 | REAC | REAC:R-HSA-21... | SMAD2/SMAD3:SMAD4 heterotrimer regulates tra... | $3.905\times10^{-3}$ |
| 21 | REAC | REAC:R-HSA-38... | PERK regulates gene expression | $4.258\times10^{-3}$ |
| 22 | REAC | REAC:R-HSA-17... | Signaling by TGF-beta Receptor Complex | $4.353\times10^{-2}$ |
| 23 | REAC | REAC:R-HSA-21... | Transcriptional activity of SMAD2/SMAD3:SMAD4 ... | $1.032\times10^{-2}$ |
| 24 | WP | WP:WP366 | TGF-beta Signaling Pathway | $2.180\times10^{-4}$ |
| 25 | WP | WP:WP364 | IL-6 signaling pathway | $8.602\times10^{-4}$ |
| 26 | WP | WP:WP4754 | IL-18 signaling pathway | $3.309\times10^{-2}$ |
| 27 | WP | WP:WP615 | Senescence and Autophagy in Cancer | $3.499\times10^{-2}$ |
| 28 | WP | WP:WP382 | MAPK Signaling Pathway | $3.913\times10^{-2}$ |
| 29 | WP | WP:WP4342 | Vitamins A and D - action mechanisms | $1.945\times10^{-5}$ |
| 30 | WP | WP:WP4482 | Vitamin D in inflammatory diseases | $7.067\times10^{-4}$ |

| | |
|---|---|
| **version** | e101_eg48_p14_baf17f0 |
| **date** | 11/23/2020, 1:54:22 PM |
| **organism** | hsapiens |

Table S19

| Tested Pair, CMS1 Score | | T-statistic | P-value, $\alpha' = 0.0085$ |
|---|---|---|---|
| MSI | MSS | **59.440057** | **0.00E+00** |
| MSI | SSC | **34.037399** | **8.38E-182** |
| MSI | ASC | **112.394144** | **0.00E+00** |
| MSS | SSC | **-42.701015** | **5.25E-270** |
| MSS | ASC | **19.224249** | **7.02E-74** |
| SSC | ASC | **133.158047** | **0.00E+00** |

Table S20

| Tested Pair, CMS2 Score | | T-statistic | P-value, α' = 0.0085 |
|---|---|---|---|
| MSI | MSS | **-79.926606** | **0.00E+00** |
| MSI | SSC | **-17.6641** | **2.25E-62** |
| MSI | ASC | **-103.464824** | **0.00E+00** |
| MSS | SSC | **82.560561** | **0.00E+00** |
| MSS | ASC | **8.471719** | **5.30E-17** |
| SSC | ASC | **-149.026347** | **0.00E+00** |

Table S21

| Tested Pair, CMS3 Score | | T-statistic | P-value, α’ = 0.0085 |
|---|---|---|---|
| MSI | MSS | **71.255748** | **0.00E+00** |
| MSI | SSC | **7.817476** | **1.36E-14** |
| MSI | ASC | **46.633286** | **3.91E-259** |
| MSS | SSC | **-113.638842** | **0.00E+00** |
| MSS | ASC | **-49.301489** | **0.00E+00** |
| SSC | ASC | **92.148023** | **0.00E+00** |

Table S22

| Tested Pair, CMS4 Score | | T-statistic | P-value, α' = 0.0085 |
|---|---|---|---|
| MSI | MSS | **-24.110683** | **2.12E-113** |
| MSI | SSC | **-17.2609** | **1.71E-59** |
| MSI | ASC | **-30.223984** | **6.54E-147** |
| MSS | SSC | **14.527264** | **4.16E-45** |
| MSS | ASC | 2.145397 | 3.21E-02 |
| SSC | ASC | **-25.071478** | **8.98E-134** |
