## Supplementary material for "Human colorectal pre-cancer atlas identifies distinct molecular programs underlying two major subclasses of pre-malignant tumors": Key Resources Table

| **Reagent or Resource** | **Source** | **Identifier** |
| --- | --- | --- |
| **Biological Samples** | | |
| COLON MAP (Polyp) | See Experimental Model and Subject Details; Data and Code Availability | Synapse scRNA-seq Level 1: [10.7303/syn23564801](javascript:;)  Synapse scRNA-seq Level 2: 10.7303/syn23630431  Synapse scRNA-seq Level 3: 10.7303/syn23520239 |
| CHTN TMA (CRC) | See Experimental Model and Subject Details; Data and Code Availability | Synapse DOI: 10.7303/syn21050481 |
| TCPS (Polyp) | See Experimental Model and Subject Details; Data and Code Availability | Synapse DOI: 10.7303/syn21050481 |
| **Critical Commercial Assays** | | |
| 7900HT Fast RT-PCR | Applied Biosystems | Catalog: 4329001 |
| Agilent 2100 Bioanalyzer | Agilent | Catalog: G2939BA |
| Qubit 2.0 Fluorometer | Invitrogen | Catalog: Q32866 |
| cBot and HiSeq2500 | Illumina | Catalog: SY–401–2501 |
| TruSeq RNA Access Library Prep Kit | Illumina | Catalog: RS-301-2001 |
| TruSeq v4 SBS kit | Illumina | Catalog: FC-401-4002 |
| TruSeq v4 PE Cluster kit | Illumina | Catalog: PE-401-4001 |
| KAPA Library quantification kit | KAPA Biosystems | Catalog: 07960140001 |
| Qubit DNA BR Assay kit | Invitrogen | Catalog: Q32850 |
| NovaSeq 6000 Reagent Kit | Illumina | Catalog: 20039236 |
| TruDrop primers | Illumina, In-house | Catalog: RS-122-2001 |
| QIAmp DNA kit | Qiagen | Catalog: 80234 |
| Lysing Matrix E | MP Bio | Catalog: 116914100 |
| RNeasy Mini | Qiagen | Catalog: 74104 |
| AllPrep DNA/RNA/miRNA Universal Kit | Qiagen | Catalog: 80224 |
| QIAamp DNA FFPE Tissue Kit | Qiagen | Catalog: 56404 |
| IN Cell Analyzer 2500 | Cytiva/GE Healthcare Life Sciences | Catalog: 29240356 |
| Cell DIVe | Cytiva/GE Healthcare Life Sciences | Catalog: 29262872 |
| Muc2, F-2 clone, A488 dye | Santa Cruz | Catalog: sc-515032 AF488 |
| Collagen Peptide, R-CHP clone, Cy3 dye | 3Helix | Catalog: RED300 |
| SNA, Lectin clone, Cy5 dye | Vector | Catalog: CL-1305-1 |
| CD11B, C67F154 clone, A488 dye | Thermo Fisher | Catalog: 53-0196-82 |
| CD45, 2D-1 clone, A546 dye | Santa Cruz | Catalog: sc-1187 AF546 |
| CD20, D-10 clone, A647 dye | Santa Cruz | Catalog: sc-393894 AF647 |
| PCNA, PC-10 clone, A488 dye | Cell Signaling | Catalog: 8580S |
| B-catenin, 12F751 clone, 550 dye | Vanderbilt Antibody and Protein Resource | Catalog: In-House |
| p-STAT3, D3A7 clone, A647 dye | Cell Signaling | Catalog: 4324S |
| pEGFR, EP774Y clone, A488 dye | Abcam | Catalog: ab205827 |
| CgA, C-12 clone, A546 dye | Santa Cruz | Catalog: sc-393941 |
| CD4, EPR6855 clone, A647 dye | Abcam | Catalog: ab196147 |
| Cox2, D5H5 clone, A488 dye | Cell Signaling | Catalog: 13596S |
| CD3d, EP4426 clone, A555 dye | Abcam | Catalog: ab208514 |
| HLA-A, EP1395Y clone, A647 dye | Abcam | Catalog: ab199837 |
| PanCK, AE1/AE3 clone, A488 dye | Thermo Fisher | Catalog: 53-9003-82 |
| OLFM4, D1E4M clone, A555 dye | Cell Signaling | Catalog: 14369S |
| CD8, C8/114B clone, A647 dye | Biolegend | Catalog: 372906 |
| Alpha-actinin, EPR2533(2) clone, A488 dye | Abcam | Catalog: ab198608 |
| CD68, KP1 clone, A546 dye | Santa Cruz | Catalog: sc-20060 AF546 |
| NaKATPase, EP1845Y clone, A647 dye | Abcam | Catalog: ab198367 |
| Vimentin, E-5 clone, A488 dye | Santa Cruz | Catalog: sc-373717 AF488 |
| Sox9, EPR14335 clone, A555 dye | Abcam | Catalog: ab202516 |
| FOXP3, 206D clone, A647 dye | Biolegend | Catalog: 320114 |
| Lysozyme, E-5 clone, A488 dye | Santa Cruz | Catalog: sc-518012 AF488 |
| SMA, 1A4 clone, Cy3 dye | Millipore Sigma | Catalog: C6198-100UL |
| ERBB2, EPR19547 clone, A647 dye | Abcam | Catalog: ab225510 |
| P-p44/42 MAPK, Rabbit Monoclonal | Cell Signaling | Catalog: 4370 |
| MUC5AC, Rabbit Monoclonal | Cell Signaling | Catalog: 61193 |
| CDX2, Rabbit Monoclonal | Cell Signaling | Catalog: 12306 |
| Midkine, Rabbit Monoclonal | Abcam | Catalog: ab52637 |
| YAP, Rabbit Monoclonal | Cell Signaling | Catalog: 14074 |
| MLH1, Rabbit Monoclonal | Abcam | Catalog: Ab92312 |
| AEC+ Substrate-Chromogen | Agilent Technologies | Catalog: K3461 |
| EnVision+ System HRP, Labeled Polymer Anti-Rabbit | Dako | Catalog: K4002 |
| **Deposited Data** | | |
| TCGA (CRC) | (Muzny et al., 2012) | <https://gdac.broadinstitute.org/>; TCGA GDAC Firehose: [2016_01_28](http://gdac.broadinstitute.org/runs/stddata__2016_01_28/data) |
| SMC (CRC) | (Lee et al., 2020) | <https://www.ncbi.nlm.nih.gov/geo/>; GEO: GSE132465 |
| **Software and Algorithms** | | |
| pCreode | (Herring et al., 2018) | <https://github.com/KenLauLab/pCreode> |
| Scanpy | (Wolf et al., 2018) | <https://github.com/theislab/scanpy> |
| Pegasus | Klarman Cell Observatory | <https://github.com/klarman-cell-observatory/pegasus> |
| pySCENIC | (Aibar et al., 2017) | <https://github.com/aertslab/pySCENIC> |
| CytoTRACE | (Gulati et al., 2020) | <https://cytotrace.stanford.edu/> |
| CMScaller | (Eide et al., 2017) | <https://github.com/peterawe/CMScaller> |
| CMSclassifier | (Guinney et al., 2015) | <https://github.com/Sage-Bionetworks/CMSclassifier> |
| Seaborn | (Waskom et al., 2020) | <https://github.com/mwaskom/seaborn> |
| Matplotlib | (Caswell et al., 2019) | <https://github.com/matplotlib/matplotlib> |
| cBioPortal | (Cerami et al., 2012; Gao et al., 2013) | <https://www.cbioportal.org/> |
| GATK4 | (Poplin et al., 2017) | <https://gatk.broadinstitute.org/hc/en-us> |
| DENDRO | (Zhou et al., 2020) | <https://github.com/zhouzilu/DENDRO> |
| Dropkick | (Heiser et al., 2020) | <https://github.com/KenLauLab/dropkick> |
| DropEst | (Petukhov et al., 2018) | <https://github.com/hms-dbmi/dropEst> |
| STAR | (Dobin et al., 2013) | <https://github.com/alexdobin/STAR> |
| Cytoscape | (Shannon et al., 2003) | <https://cytoscape.org/> |
| g:Profiler | (Raudvere et al., 2019) | <https://biit.cs.ut.ee/gprofiler/> |
| Scipy | (Virtanen et al., 2020) | <https://www.scipy.org/> |
| Sinto | (Tim Stuart, 2018) | <https://github.com/timoast/sinto> |
| Dendextend | (Galili, 2015) | <https://github.com/talgalili/dendextend> |
| Numpy | (Harris et al., 2020) | <https://numpy.org/> |
| Pandas | (McKinney, 2010) | <https://pandas.pydata.org/> |
| BWA | (Li and Durbin, 2009) | <https://sourceforge.net/projects/maq/> |
| Genome Studio | Illumina | <https://www.illumina.com/techniques/microarrays/array-data-analysis-experimental-design/genomestudio.html> |
| ANNOVAR | (Wang et al., 2010; Yang and Wang, 2015) | <https://github.com/WGLab/doc-ANNOVAR> |
| Picard | (Broad Institute, 2019) | <https://broadinstitute.github.io/picard/> |
| Sambamba | (Tarasov et al., 2015) | <https://github.com/biod/sambamba> |
| Python 3.8 | (van Rossum and Drake, 2009) | <https://www.python.org/> |
| R 4.0.2 | (R Core Team (2020), 2020) | <https://www.r-project.org/> |
| lme4 | (Bates et al., 2020) | <https://github.com/lme4/lme4> |
| lmerTest | (Kuznetsova et al., 2017) | <https://github.com/runehaubo/lmerTestR> |
| emmeans | (Lenth et al., 2020) | <https://github.com/rvlenth/emmeans> |

Aibar, S., González-Blas, C.B., Moerman, T., Huynh-Thu, V.A., Imrichova, H., Hulselmans, G., Rambow, F., Marine, J.C., Geurts, P., Aerts, J., et al. (2017). SCENIC: Single-cell regulatory network inference and clustering. Nat. Methods.

Bates, K., Bolker, B., Walker, S., Christiensen, R.H., Singmann, H., Dai, B., and Scheipl, F. (2020). Ime4.

Broad Institute (2019). Picard toolkit.

Caswell, T.A., Droettboom, M., Hunter, J., Lee, A., Firing, E., Stansby, D., Klymak, J., Andrade, E.S. de, Nielsen, J.H., Varoquaux, N., et al. (2019). matplotlib/matplotlib: REL: v3.1.1.

Cerami, E., Gao, J., Dogrusoz, U., Gross, B.E., Sumer, S.O., Aksoy, B.A., Jacobsen, A., Byrne, C.J., Heuer, M.L., Larsson, E., et al. (2012). The cBio Cancer Genomics Portal: An Open Platform for Exploring Multidimensional Cancer Genomics Data. Cancer Discov. *2*, 401 LP – 404.

Dobin, A., Davis, C.A., Schlesinger, F., Drenkow, J., Zaleski, C., Jha, S., Batut, P., Chaisson, M., and Gingeras, T.R. (2013). STAR: Ultrafast universal RNA-seq aligner. Bioinformatics.

Eide, P.W., Bruun, J., Lothe, R.A., and Sveen, A. (2017). CMScaller: an R package for consensus molecular subtyping of colorectal cancer pre-clinical models. Sci. Rep. *7*, 16618.

Galili, T. (2015). dendextend: an R package for visualizing, adjusting and comparing trees of hierarchical clustering. Bioinformatics *31*, 3718–3720.

Gao, J., Aksoy, B.A., Dogrusoz, U., Dresdner, G., Gross, B., Sumer, S.O., Sun, Y., Jacobsen, A., Sinha, R., Larsson, E., et al. (2013). Integrative Analysis of Complex Cancer Genomics and Clinical Profiles Using the cBioPortal. Sci. Signal. *6*, pl1 LP-pl1.

Guinney, J., Dienstmann, R., Wang, X., De Reyniès, A., Schlicker, A., Soneson, C., Marisa, L., Roepman, P., Nyamundanda, G., Angelino, P., et al. (2015). The consensus molecular subtypes of colorectal cancer. Nat. Med.

Gulati, G.S., Sikandar, S.S., Wesche, D.J., Manjunath, A., Bharadwaj, A., Berger, M.J., Ilagan, F., Kuo, A.H., Hsieh, R.W., Cai, S., et al. (2020). Single-cell transcriptional diversity is a hallmark of developmental potential. Science (80-. ).

Harris, C.R., Millman, K.J., van der Walt, S.J., Gommers, R., Virtanen, P., Cournapeau, D., Wieser, E., Taylor, J., Berg, S., Smith, N.J., et al. (2020). Array programming with NumPy. Nature *585*, 357–362.

Heiser, C.N., Wang, V.M., Chen, B., Hughey, J.J., and Lau, K.S. (2020). Automated quality control and cell identification of droplet-based single-cell data using dropkick. BioRxiv.

Herring, C.A., Banerjee, A., McKinley, E.T., Simmons, A.J., Ping, J., Roland, J.T., Franklin, J.L., Liu, Q., Gerdes, M.J., Coffey, R.J., et al. (2018). Unsupervised Trajectory Analysis of Single-Cell RNA-Seq and Imaging Data Reveals Alternative Tuft Cell Origins in the Gut. Cell Syst. *6*, 37-51.e9.

Kuznetsova, A., Brockhoff, P.B., and Christensen, R.H.B. (2017). lmerTest Package: Tests in Linear Mixed Effects Models. J. Stat. Software; Vol 1, Issue 13 .

Lee, H.O., Hong, Y., Etlioglu, H.E., Cho, Y.B., Pomella, V., Van den Bosch, B., Vanhecke, J., Verbandt, S., Hong, H., Min, J.W., et al. (2020). Lineage-dependent gene expression programs influence the immune landscape of colorectal cancer. Nat. Genet.

Lenth, R., Singmann, H., Love, J., Buerkner, P., and Herve, M. (2020). Package ‘ emmeans .’

Li, H., and Durbin, R. (2009). Fast and accurate short read alignment with Burrows-Wheeler transform. Bioinformatics.

McKinney, W. (2010). {D}ata {S}tructures for {S}tatistical {C}omputing in {P}ython. In {P}roceedings of the 9th {P}ython in {S}cience {C}onference, S. van der Walt, and J. Millman, eds. pp. 56–61.

Muzny, D.M., Bainbridge, M.N., Chang, K., Dinh, H.H., Drummond, J.A., Fowler, G., Kovar, C.L., Lewis, L.R., Morgan, M.B., Newsham, I.F., et al. (2012). Comprehensive molecular characterization of human colon and rectal cancer. Nature *487*, 330–337.

Petukhov, V., Guo, J., Baryawno, N., Severe, N., Scadden, D.T., Samsonova, M.G., and Kharchenko, P. V. (2018). dropEst: Pipeline for accurate estimation of molecular counts in droplet-based single-cell RNA-seq experiments. Genome Biol.

Poplin, R., Ruano-Rubio, V., DePristo, M.A., Fennell, T.J., Carneiro, M.O., Van der Auwera, G.A., Kling, D.E., Gauthier, L.D., Levy-Moonshine, A., Roazen, D., et al. (2017). Scaling accurate genetic variant discovery to tens of thousands of samples. BioRxiv.

R Core Team (2020) (2020). R: A language and environment for statistical computing. R A Lang. Environ. Stat. Comput. R Found. Stat. Comput. Vienna, Austria.

Raudvere, U., Kolberg, L., Kuzmin, I., Arak, T., Adler, P., Peterson, H., and Vilo, J. (2019). g:Profiler: a web server for functional enrichment analysis and conversions of gene lists (2019 update). Nucleic Acids Res. *47*, W191–W198.

van Rossum, G., and Drake, F.L. (2009). Python 3 Reference Manual.

Shannon, P., Markiel, A., Ozier, O., Baliga, N.S., Wang, J.T., Ramage, D., Amin, N., Schwikowski, B., and Ideker, T. (2003). Cytoscape: A software Environment for integrated models of biomolecular interaction networks. Genome Res.

Tarasov, A., Vilella, A.J., Cuppen, E., Nijman, I.J., and Prins, P. (2015). Sambamba: Fast processing of NGS alignment formats. Bioinformatics.

Tim Stuart, W.K. (2018). Sinto: single-cell analysis tools.

Virtanen, P., Gommers, R., Oliphant, T.E., Haberland, M., Reddy, T., Cournapeau, D., Burovski, E., Peterson, P., Weckesser, W., Bright, J., et al. (2020). SciPy 1.0: fundamental algorithms for scientific computing in Python. Nat. Methods *17*, 261–272.

Wang, K., Li, M., and Hakonarson, H. (2010). ANNOVAR: Functional annotation of genetic variants from high-throughput sequencing data. Nucleic Acids Res.

Waskom, M., Botvinnik, O., Gelbart, M., Ostblom, J., Hobson, P., Lukauskas, S., Gemperline, D.C., Augspurger, T., Halchenko, Y., Warmenhoven, J., et al. (2020). mwaskom/seaborn: v0.11.0 (Sepetmber 2020).

Wolf, F.A., Angerer, P., and Theis, F.J. (2018). SCANPY: Large-scale single-cell gene expression data analysis. Genome Biol.

Yang, H., and Wang, K. (2015). Genomic variant annotation and prioritization with ANNOVAR and wANNOVAR. Nat. Protoc.

Zhou, Z., Xu, B., Minn, A., and Zhang, N.R. (2020). DENDRO: genetic heterogeneity profiling and subclone detection by single-cell RNA sequencing. Genome Biol. *21*, 10.
